# Integrative Multi-omics Analysis of the Human Skeletal Muscle Response to Endurance or Resistance Exercise: Findings from the Molecular Transducers of Physical Activity Consortium (MoTrPAC)

**DOI:** 10.64898/2026.03.04.705181

**Authors:** Hasmik Keshishian, Gina M. Many, Gregory Smith, Natalie M. Clark, Gayatri Iyer, Patrick Hart, Malene E. Lindholm, Samuel Montalvo, Zidong Zhang, Christopher Jin, James A. Sanford, Steven A. Carr, Joshua N. Adkins, D.R. Mani, Sue C. Bodine, Scott Trappe, Joseph A. Houmard, Nicolas Musi, Kim M. Huffman, William E. Kraus, Lauren M. Sparks, Anna E. Thalacker-Mercer, Stuart C. Sealfon, Ashley Y. Xia, Daniel H. Katz, Christopher B. Newgard, Charles F. Burant, Paul M. Coen, Bret H. Goodpaster, MoTrPAC Study Group

## Abstract

The Molecular Transducers of Physical Activity Consortium (MoTrPAC) was established to systematically characterize the molecular basis of the health benefits of exercise. Here, we present the integrative, multi-omics response of human skeletal muscle to acute endurance (EE) or resistance (RE) exercise. Distinct temporal omic responses were observed, with RE resulting in a greater number of differentially abundant (DA) features across omes and timepoints. Temporal multi-omic dynamics to EE or RE were used to identify transcriptional regulatory hubs converging around MEF2A and NFIC regulation of autophagy, angiogenesis and metabolism in the acute exercise response. Further, we propose a model where early RE-specific phosphoproteome changes override epigenetic modifications to downregulate E3 ubiquitin-protein ligase transcripts involved in protein turnover. Additional findings include suppression of HIPK3 kinase signature linked to the acute exercise regulation of sarcomeric proteins TTN, AKTN2 and LMOD2. Our data demonstrate distinct temporal regulation across the multi-omic landscape of human skeletal muscle, with EE and RE eliciting both common and unique dynamic molecular signatures.

## Introduction

Regular exercise elicits a myriad of health benefits, including maintaining mobility, reducing risk of chronic diseases including type 2 diabetes, cancer, and cardiovascular diseases and increasing healthy lifespan.^1,2^ Skeletal muscle is essential for movement, contributes to systemic metabolic homeostasis, and functions as an endocrine organ to facilitate intercellular, tissue and organ crosstalk^3^. Skeletal muscle is also highly plastic and adapts to regular exercise by increasing mitochondrial biogenesis, vascularization, muscle mass and strength. These training effects are the aggregate result of coordinated molecular responses in muscle during individual acute bouts of exercise.

While much is known about the molecular underpinnings of exercise, prior studies have generally limited the description of acute changes to single-ome profiles, including transcriptome,^4^ phospho/proteome,^5,6^ lipidome,^7^ metabolome or epigenome.^8^ Prior studies have also been limited by small numbers of subjects ^4^ or have used meta analyses across various assorted studies, resulting in incomplete information about the temporal responses to exercise.^4^ As such, our knowledge about the integrated, dynamic and coordinated molecular responses to acute exercise has been limited. Endurance and resistance exercise elicit distinct adaptations that improve muscle function and health.^9^ Endurance exercise, typically defined as continuous, repetitive movement of major muscle groups, generally increases mitochondrial fatty acid oxidation (FAO) and alters redox state resulting in downstream signal transduction (HIF-1, PGC-1, CREB) that drives mitochondrial and oxidative metabolism adaptations. Resistance exercise involves lower repetitions of higher load which provides greater neuromuscular recruitment of type II fibers, non-oxidative glycolytic flux, and a greater mechanical/neuronal stimulus. The resulting mechanotransduction and activation of mTOR and Akt and inhibition of FOXOs stimulate greater protein synthesis and hypertrophic and structural adaptations.^10–12^

The Molecular Transducers of Physical Activity Consortium (MoTrPAC) was established to systematically characterize the molecular basis of the health benefits of exercise.^13^ Here we profile the integrative, temporal multi-omics response to exercise in skeletal muscle obtained from sedentary human participants who were randomized to either acute endurance exercise (EE) or resistance exercise (RE) and relative to a non-exercised control (CON) groups.^14^ *Vastus lateralis* muscle biopsies were collected before and at multiple time points after a sub-maximal acute bout of EE, RE or CON, and were analyzed using epigenomics (ATAC-seq, methylCap), transcriptomics (RNA-seq), proteomics, phosphoproteomics, and metabolomics profiling. The multi-omic and temporal nature of this study, benchmarked to time-matched controls, provides insight into time-resolved, multi-omic exercise responses in human skeletal muscle, highlighting common and specific signatures between EE and RE. The study represents a fundamentally important step towards understanding the molecular mechanisms for the health benefits of exercise and developing personalized exercise prescriptions.

## Results

### Study design and experimental overview

Muscle biopsies were analyzed from a total of 174 healthy sedentary adults who met eligibility criteria,^13^ and were randomized to non-exercise control (CON, n = 37), endurance exercise (EE, n = 64), or resistance exercise (RE, n = 73). These participants were enrolled prior to the MoTrPAC study suspension due to the COVID-19 pandemic, and the results from this cohort will guide analysis of the larger post-suspension cohort of ∼1,540 participants. A detailed description of the study design, screening, baseline phenotyping of the Pre-COVID-19 cohort as well as details of the acute exercise protocol can be found in the clinical landscape ^15^ and protocol papers.^14^ The CON group was included to account for time of day, fasting, stress of tissue biopsies, and other non-exercise factors that might confound multi-omic responses to exercise. Participants in this cohort were predominantly female (72%), with a mean age of 41 ± 15 years. Subject randomization resulted in similar age, weight, BMI, cardiorespiratory fitness, and strength between groups (Table 1 and Supplementary Table 1). A *vastus lateralis* biopsy was obtained from all participants before exercise. EE participants completed a session of cycling exercise (40 minutes at 65% VO_2_peak), while RE participants completed a circuit of eight resistance exercises, and the CON group rested in a supine position for 40 minutes. A *vastus lateralis* biopsy was obtained from all participants before exercise or control, and subsequently participants were randomized to have post-exercise biopsies at either 15 minutes (*Early*), 3.5 hours (*Middle*), 24 hours (*Late*), or at all post-exercise time points (*All*) (Fig. 1a). The number of samples profiled for epigenome, transcriptome, metabolome, proteome and phosphoproteome by time point and group are described in Extended Data Fig. 1a. The overarching goal of the analysis was to define temporal multi-omic responses and molecular networks modulated by acute exercise, with emphasis on unveiling both shared and unique molecular signals driving modality-specific responses.

**Figure 1:**
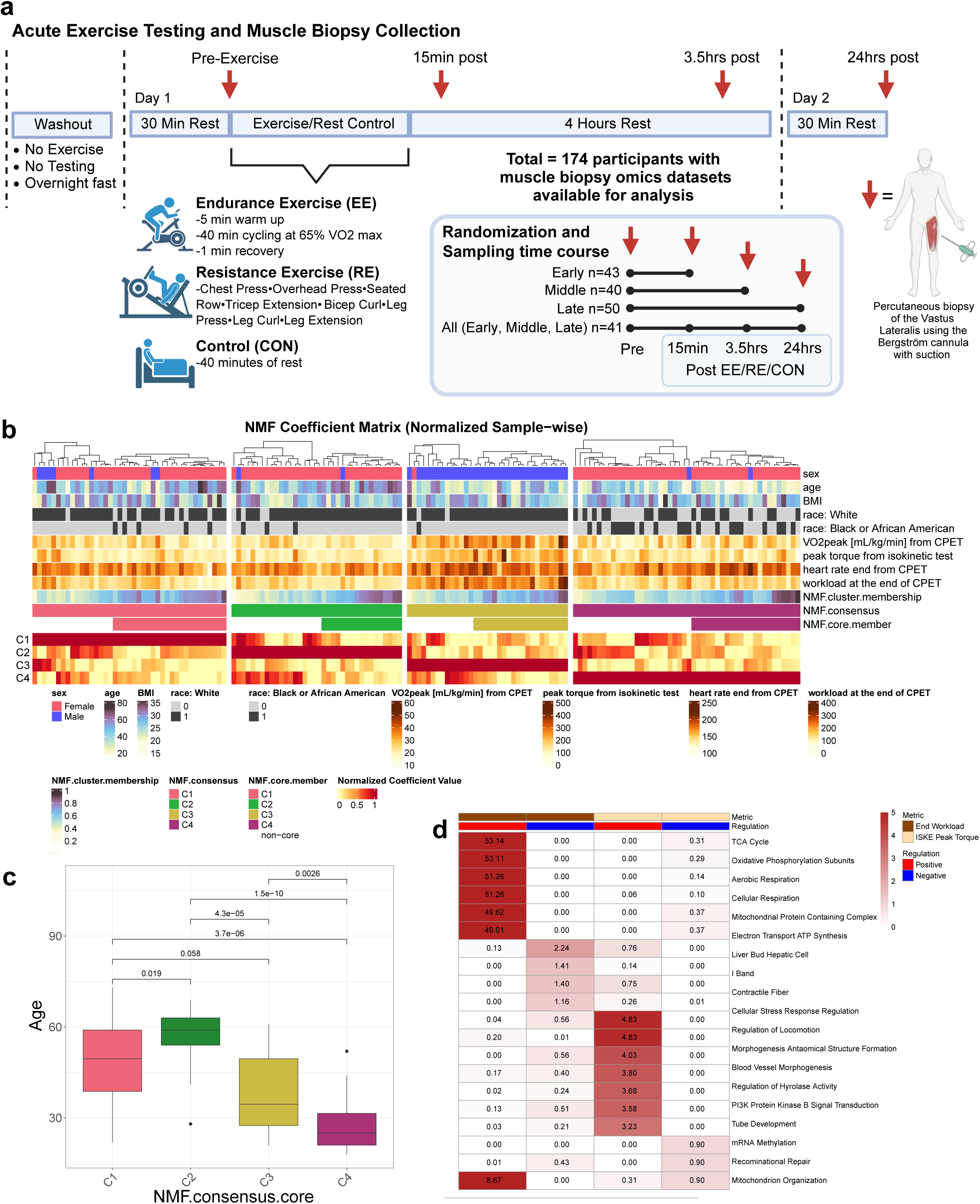
Study design and integrative multi-omics analysis. **a.** Schematic overview of MoTrPAC acute exercise protocol. Patients arrived at the clinical site in the morning following an overnight fast and complete 30 minutes of rest. After baseline biopsies of the *vastus lateralis skeletal muscle biopsies*, participants were randomized to complete endurance exercise (EE), resistance exercise (RE) or controlled rest (CON). Post exercise muscle biopsies were obtained at 15 min, 3.5 hours and/or 24 hours post-exercise based on randomization to early, middle, late or all timepoint collections per subject. The number of subjects per group is indicated. **b.** NMF coefficient matrix showing the results of NMF clustering on muscle pre-exercise RNA, proteome, and phosphoproteome of n = 159 samples. **c.** Boxplot of NMF core samples showing significant age differences. Reported P-values are from two-sample t-tests on each pairwise comparison. **d.** Heatmap of overrepresentation analysis (ORA) of genes identified by linear model to be significantly associated with select clinical metrics. The first two columns reflect the significant genes whose pre-exercise expression is positively or negatively correlated, respectively, with workload (Watts). The last two columns reflect the genes whose 3.5 hour exercise response are positively or negatively correlated, respectively, with isometric test (ISKE) Peak Torque. Representative pathways among the top ten most significantly associated are included for each ORA comparison. Colors and the corresponding numbers of each box reflect the -log10(adj p-value) of ORA pathway enrichment.

**Table 1:**
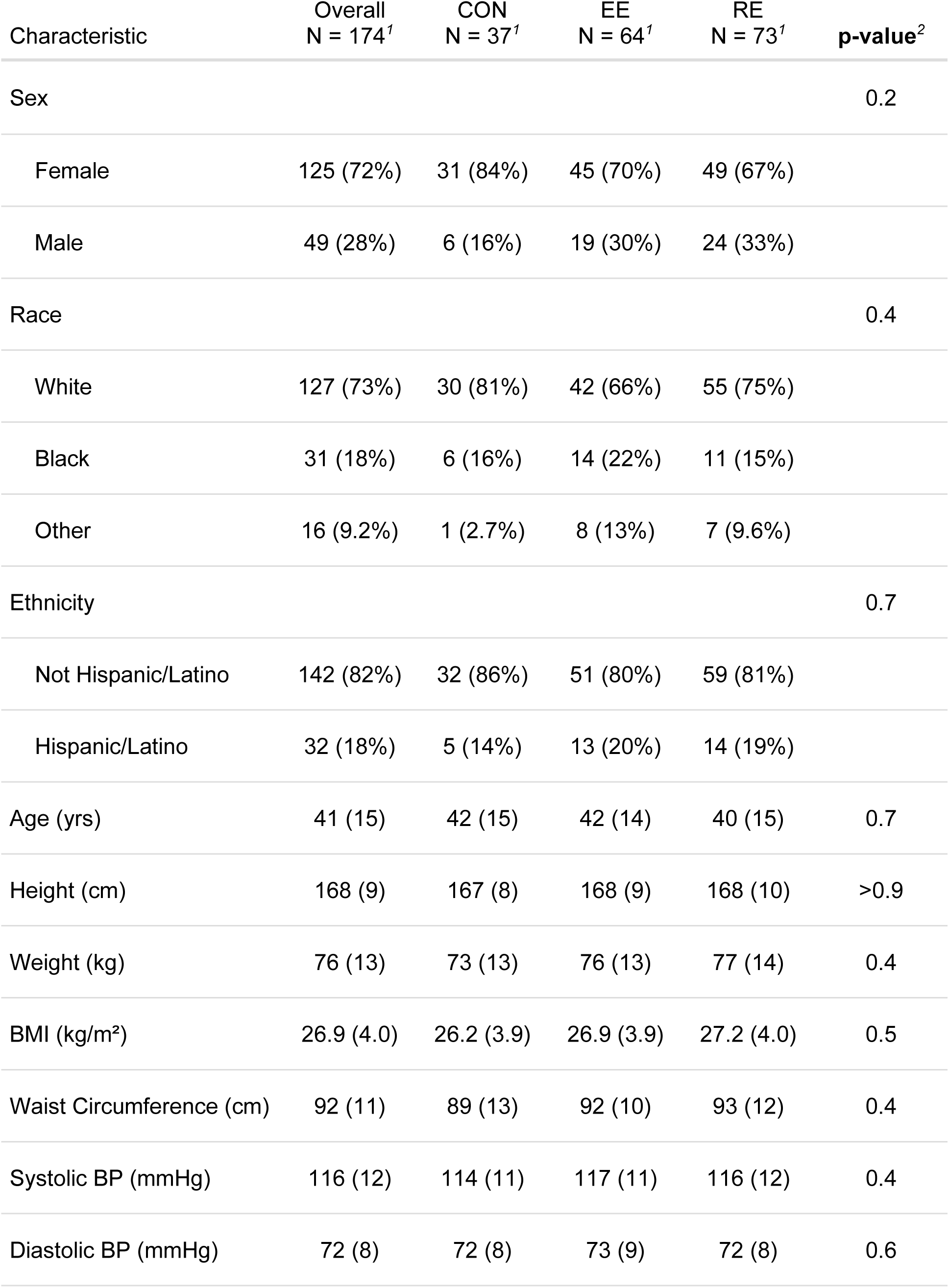
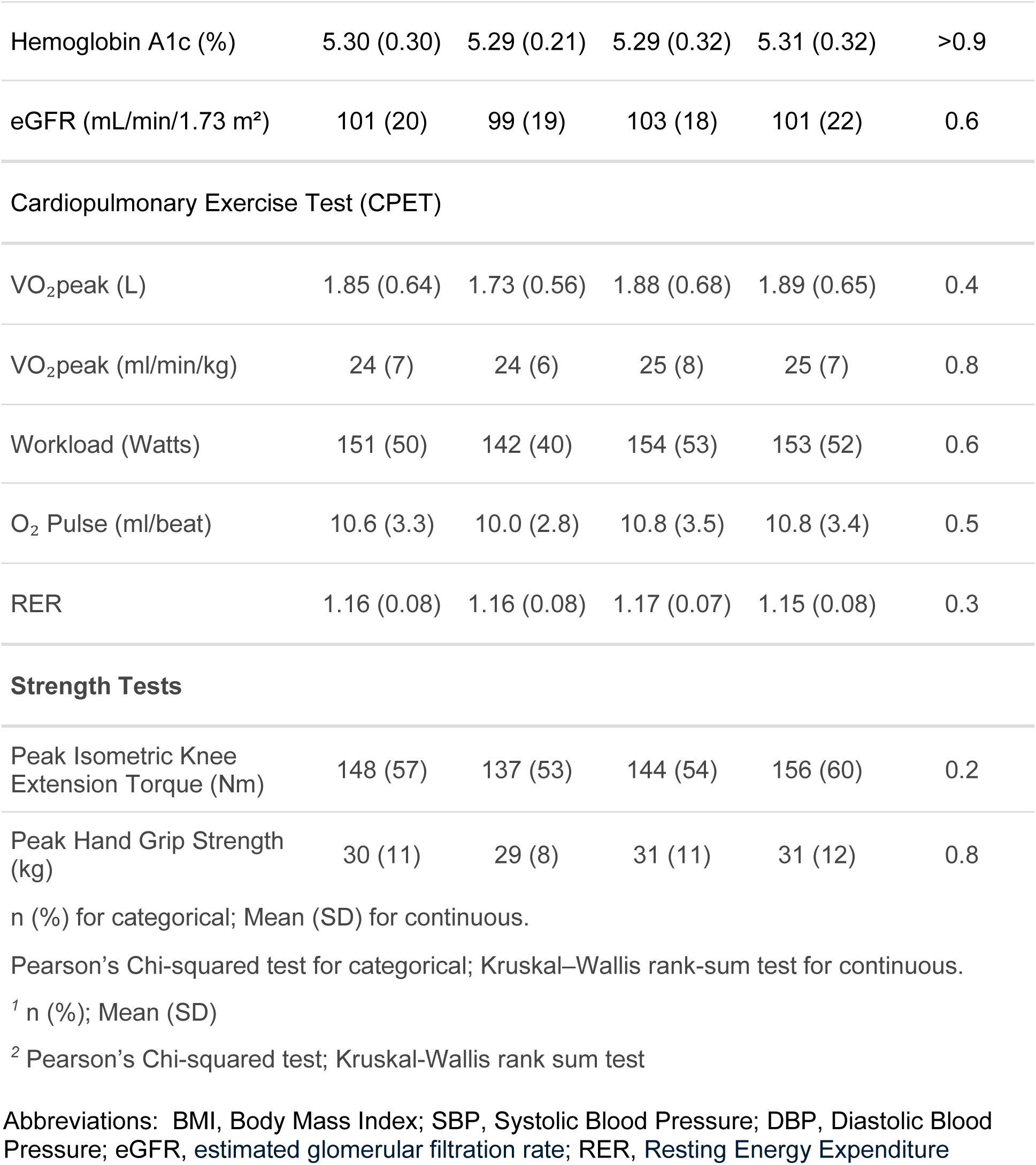
Baseline characteristics of study groups.

### Multi-omic associations with baseline participant characteristics

We first explored relationships between participant characteristics and muscle molecular data. The goal was twofold; to understand whether baseline muscle molecular data explained interindividual variability in participant phenotype including those related to fitness and strength, and to identify phenotypes that may affect subsequent analysis of the acute exercise data. To do this, we performed non-negative matrix factorization (NMF) clustering on pre-exercise transcriptomics, proteomics, and phosphoproteomics data of all participants. Metabolomics data were excluded from NMF clustering because 20 participants had not been assayed across all metabolomic platforms (targeted and untargeted), and would have reduced the sample size by approximately 10% (see Methods). Therefore, metabolomics data were evaluated post hoc instead.

Multi-omics NMF clustering stratified the cohort into 4 clusters (Supplementary Table 2), which differed by sex and age (Fig. 1b). Male participants were predominantly represented in cluster 3, while the other clusters were female-dominant. Further, clusters 3 and 4 skewed younger in age (Fig. 1c). Additionally, while cluster 4 was significantly enriched for Black women (Fisher test, p-value < 0.01), we did not have sufficient statistical power to investigate further, nor adjust statistical models without introducing inestimable coefficients. Baseline physiological measurements were linked to cluster separations by sex and age. Muscle strength (Peak torque) and cardiorespiratory fitness (VO_2_peak, peak power output and peak heart rate) were significantly higher in male-dominant cluster 3 (Extended Data Fig. 1b). Using Pre-ranked Correlation-Adjusted MEan-RAnk (CAMERA-PR)^16^, distinct enrichments were observed across the clusters and included enrichments in oxidative phosphorylation in the younger age clusters 3 and 4 (Supplementary Results, Extended Data Fig. 1c). Examination of baseline metabolomics data also revealed distinct patterns of energy-associated metabolites, including lower levels of triglycerides and higher levels of amino acids strongly associating with male-skewed cluster 3 (Supplementary Results, Extended Data Fig. 1d). Overall, these results suggest that NMF clusters capture distinct baseline metabolic states that may partly reflect the age- and sex-associated structure of the cohort. These results also underscore the importance of controlling for sex and age in statistical models.

Next, we sought to investigate associations of baseline physiological metrics with temporal changes in omics features, while adjusting for age and sex. Physiological metrics were divided into separate correlation clusters (Extended Data Fig. 2a) with strong positive correlation between VO_2_ peak and peak power output (Watts) attained during the Cardiopulmonary Exercise Test (CPET), and a weaker positive correlation between CPET peak power and knee extension peak torque (Extended Data Fig. 2b,c). Of all phenotypes assessed, CPET peak power displayed the most frequent correlation to pre-exercise proteins (n=448) and the second most frequent correlation to transcripts (n=254) in sedentary muscle (Extended Data Fig. 2d,e). Post-acute exercise-induced muscle transcripts and proteins were also correlated to baseline phenotypes, with peak torque displaying the most frequent associations with muscle transcripts at 3.5 hours (n=244), CPET rating of perceived exertion (RPE) most frequently associated with phosphopeptides at 15 minutes (n=263) (Extended Data Fig. 2d,f), and systolic blood pressure (BP) most frequently associated with phosphopeptides (n=346) and metabolites (n=25) at 24 hours (Extended Data Fig. 2f,g). Genes whose baseline expression was positively correlated with Watts were highly enriched for cellular respiration and mitochondrial pathways (Fig. 1d). Thirty two percent of these were also positively correlated with Watts in the proteomics data (Extended Data Fig. 2e), including ATP5PB and other genes associated with aerobic respiration (Extended Data Fig. 2h). Locomotion regulation and circulatory system-associated pathways were enriched for genes whose 3.5 hour exercise was positively correlated with peak torque (Fig. 1d), including the proteoglycan CSPG4, with roles in microvascular development^17^(Extended Data Figure 2i), while negatively correlated genes were enriched for mitochondrial organization and DNA repair pathways including AP5S1 (Extended Data Fig. 2j). Together, these findings demonstrate that baseline physiological phenotypes are reflected in the skeletal muscle molecular landscape and are associated with interindividual variation in the molecular response to acute exercise.

### RE elicits a broader and more sustained multi-omic response than EE

We assessed the temporal molecular responses by ome and exercise modality using a difference-in-change model, by comparing changes per ome in response to EE or RE relative to time-matched changes in CON (Methods). The greatest number of molecular changes, defined as differentially abundant (DA) features (adjusted p-value <0.05), were observed in the transcriptome and phosphoproteome, followed by the epigenome, metabolome, and proteome (Fig. 2a, Extended Data Fig. 3a, Supplementary Table 3). RE consistently resulted in a greater number of DA features than EE at all time points (Fig. 2a).

**Figure 2:**
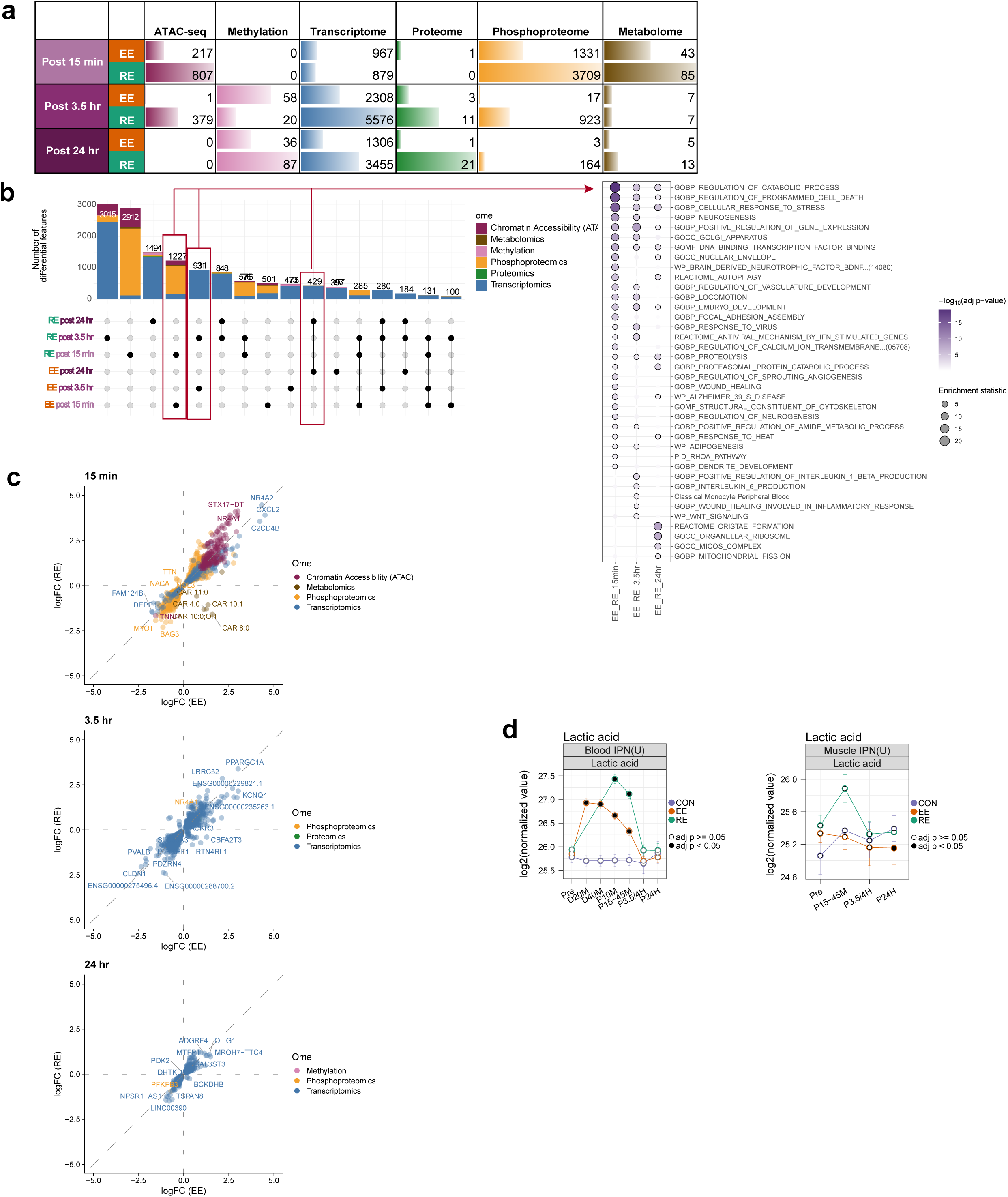
Differential regulation of multi-omics dataset. **a.** Total number of differentially abundant (DA) features for each assay by time points and exercise modality; different omes in each bar are represented by ome-specific colors. **b.** Left panel: Upset plot showing overlap of DA features across the exercise modalities and time points. Different omes in each bar are represented by ome-specific colors. Numbers on the bars indicate the total number of features per intersection, where a vertical line indicates shared omic features between modalities and/or timepoints. Right panel: Heatmap illustrating selected pathways from Overrepresentation analysis (ORA) of shared features between EE and RE at 15 minute, 3.5 hour, and 24 hour time points. **c.** Scatter plot of LogFC of shared features between EE and RE at each time point depicting concordance and divergence between EE and RE; omes are represented by ome-specific colors. **d.** Temporal trajectory of Lactic acid in blood (see multi-omics Landscape; in preparation/submission; for sampling details). Where EE = Endurance exercise (orange line), RE = resistance exercise (green line), CON = control group (purple), Pre = pre-exercise, D20M = during 20 minutes, D40M = during 40 minutes, P10M = 10 minutes post exercise, P15-45M = 15, 30, 45 minutes post exercise (depending on tissue), P3.5-4H = 3.5 or 4 hours post exercise (depending on tissue), P24H = 24 hours post exercise. Y-axis shows Log2 transformed raw means. Significance (adj. P value < 0.05) indicated by black dot per timepoint per group.

To further investigate the acute exercise response, we examined directionality agnostic time-point-specific DA feature overlap between exercise modalities and associated pathway enrichments of overlapping features (Fig. 2b, Supplementary Results). While overlapping DA features were broadly directionally concordant across omes and modalities (Fig. 2c), medium to long-chain acylcarnitines (>6 carbons) increased at 15 min with EE and decreased with RE (Extended Data Fig. 3b)^18^. These fatty acid conjugates are required for mitochondrial import and β-oxidation–supporting increased mitochondrial lipid flux following EE. Alternatively, increased reliance on glycolysis with RE, results in lower mitochondrial lipid import in muscle via malonyl-CoA production^19^ and elevated peak blood lactate (Fig. 2d), which inhibits lipolysis in adipose tissue^20^. At 3.5 and 24 hours post-exercise, shared DA features remained broadly concordant between exercise modalities (Fig. 2c). Among several DA proteins only one was shared between EE and RE; muscle specific E3 ubiquitin ligase (TRIM63), which regulates proteasomal degradation, was upregulated at 3.5 hours (Extended Data Fig. 3c). Overall, these findings demonstrate that acute EE and RE induce largely concordant molecular responses while also revealing distinct temporal and modality-specific signatures. RE was associated with a broader and more sustained multi-omic response (Fig. 2a,b), whereas early metabolic differences, including divergent acylcarnitine responses, highlighted distinct patterns of substrate utilization between exercise modalities. These differences became more pronounced during recovery, particularly at the proteomic level (Extended Data Fig. 3c), demonstrating that the molecular responses to the intensity and regimen of EE and RE used in this study progressively diverge over the post-exercise period. Modality-specific responses to acute exercise are linked to metabolic and mechanical stimuli

### Modality-specific responses to acute exercise are linked to metabolic and mechanical stimuli

We next performed gene set enrichment analysis utilizing CAMERA-PR^16^ to identify biological processes affected by exercise. The transcriptomic response revealed several early shared enrichments between EE and RE, including pathways related to stem cell fate commitment, hypertrophy, MAPK, IL-6, and nerve growth factor (NGF) (Fig. 3a). RE generally resulted in a greater magnitude and duration of pathway enrichments. EE-specific transcriptomics enrichments were most pronounced at 24 hours and included pathways related to mitochondrial fission, FAO, and mitochondrial gene expression. Expanding pathway annotation to the MitoCarta 3.0 database revealed overlapping mitochondrial enrichments across modalities, to include OXPHOS subunits and cristae formation, albeit EE induced more robust or isolated effects at 24 hours (Fig. 3b, Supplementary Table 4). While some muscle remodeling pathways were shared between modalities, RE often triggered a prolonged expression of genes key to muscle remodeling processes, including inflammation (*IL18*, *IL1R1*), fibroblast growth factor signaling (*FGF6/12*, *FGFR1/4*), and myogenesis (*MYF6*, *MYOD1*, *MEF2C*) (Extended Data Fig. 4a,b), emphasizing modality divergent effects on muscle structural remodeling (Extended Data Fig. 4a,b).^21^ RE uniquely resulted in enrichment of autophagy, translation and contractile fiber pathways at 3.5 hours followed by positive regulation of telomerase at 24 hours, altogether supporting RE-biased induction of structural remodeling and tissue repair pathways.

**Figure 3:**
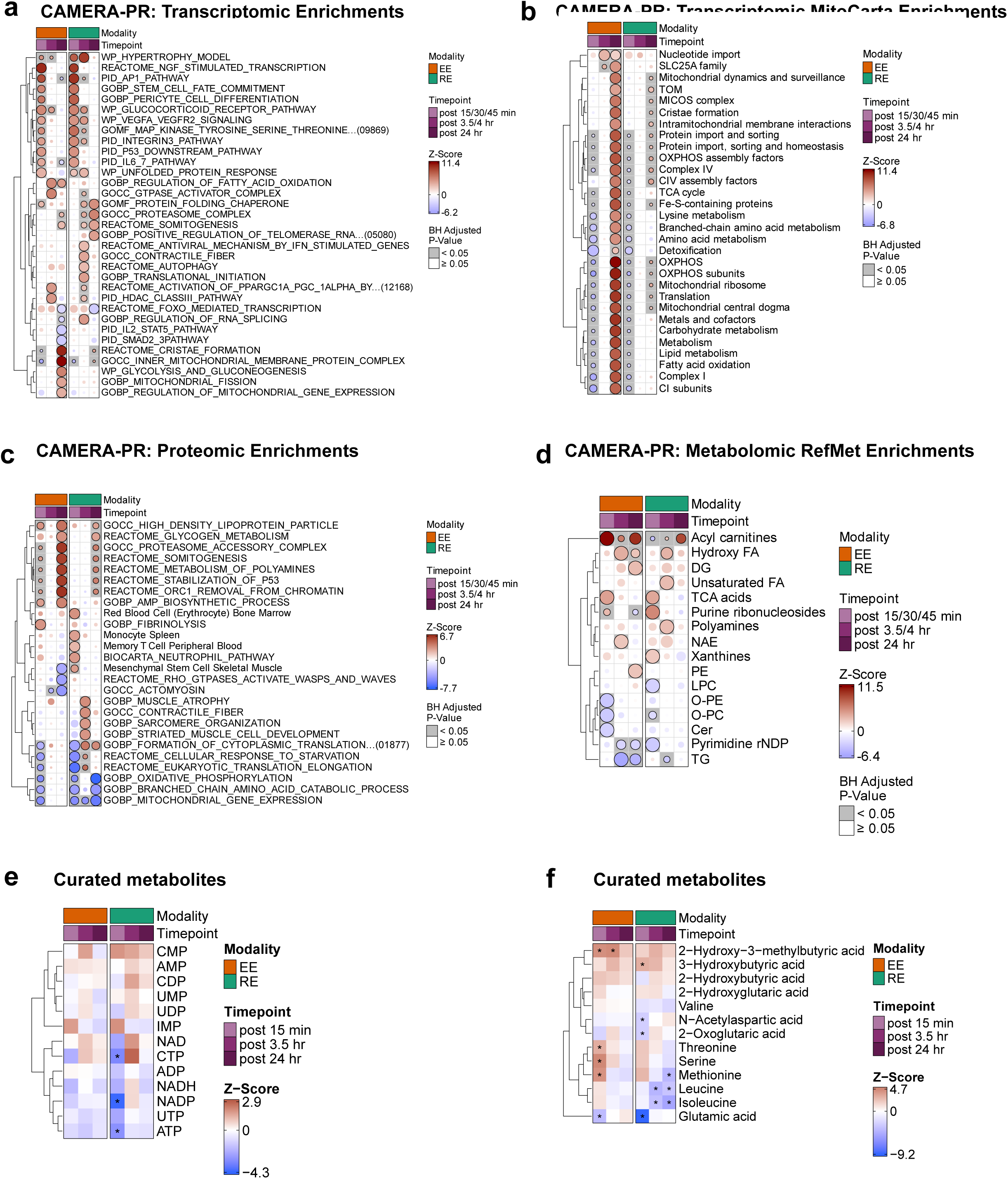
Pathway enrichment analysis. **a.** Heatmap of CAMERA-PR (Correlation Adjusted MEan RAnk gene set test—PreRanked) pathway enrichment analysis of transcriptomics data across the exercise modalities and time points using MSigDB C2 and C5 databases and the CellMarker 2.0 database. Shown are top most significant non-redundant manually curated pathways of biological relevance per timepoint that are significant (adj. P-value < 0.05). **b.** Heatmap of CAMERA-PR enrichment of transcriptomics dataset using MitoCarta 3.0 database displaying significantly enriched terms (adj. P-value < 0.05). **c.** Heatmap of CAMERA-PR pathway enrichment analysis of proteomics data across the exercise modalities and time points using MSigDB C2 and C5 databases and the CellMarker 2.0 database. Shown are pathways that are significant (adj. P-value < 0.05) per contrast, and manually curated to reduce term redundancy and display the broad biological effects of exercise on the skeletal muscle. **d.** Heatmap of RefMet (Reference analysis of Metabolite names) analysis of metabolomics dataset; displaying significantly enriched (adj. P-value < 0.05) RefMet subclasses. **e.** Heatmap of manually curated energy-carrying nucleotides and their metabolites across timepoints and modalities; * represents statistical significance at adj. P-value < 0.05. **f.** Heatmap of manually curated metabolites, relevant to skeletal muscle energy production, across timepoints and modalities; * represents statistical significance at adj. P-value < 0.05.

At the proteome level, EE induced an early upregulation of fibrinolysis-associated proteins, whereas RE upregulated proteomic terms related to mesenchymal cells, erythrocytes, and monocytes at 15 minutes post-exercise (Fig. 3c). RE additionally upregulated proteomic pathways related to muscle atrophy and sarcomeric components at 3.5 hours post-exercise, consistent with eccentric loading induced muscle damage, immune cell chemotaxis, stem cell activation and remodeling^22^. By 24 hours, modality-specific differences were even more pronounced–EE induced unique upregulation of AMP biosynthetic processes and downregulation of terms related to mesenchymal stem cells, whereas RE upregulated proteomic pathways related to translation and prolonged suppression of mitochondrial proteins (Fig. 3c; Extended Data Fig. 4c).

Among shared metabolomic enrichments included upregulation of the TCA cycle acids (Fig. 3d), including increases in citrate and isocitrate (Supplementary Table 3). RE led to a greater early depletion of the nucleotides ATP and CTP (Fig. 3e), and increase in the ketone 3−hydroxybutyric acid at 15 minutes (Fig. 3f) following RE. RE increased levels of hypoxanthine, inosine and xanthosine, consistent with greater nucleotide degradation under energetic stress (Extended Data Fig. 4d). In contrast, EE resulted in an early downregulation of ceramides, which are associated with skeletal muscle insulin resistance (Fig. 3d).^23^ Additionally, EE induced a strong upregulation of 2−hydroxy−3−methylbutyric acid at early and middle timepoints(Fig. 3f). Accumulation of 2−hydroxy−3−methylbutyric acid, a BCAA catabolite, along with increases in methionine, serine, and threonine early post-EE, support an increase in amino acid turnover.^24^ RE induced significant reductions in isoleucine and leucine across the exercise bout, perhaps reflecting increased utilization for *de novo* protein synthesis following RE. These findings are supported by studies showing substantial protein turnover (synthesis and breakdown) following RE,^25^ coupled with greater induction of myofibrillar protein synthesis compared to EE.^26^ Together, our findings highlight modality-divergent metabolomic responses to acute exercise with EE resulting in increased mitochondrial and lipid metabolic signatures and RE linked to greater contractile stress and remodeling signatures.

### Integrated multi-omic analysis reveals molecular coordination within skeletal muscle during acute exercise

To define exercise-regulated coordinated temporal relationships we applied Pathway-Level Information ExtractoR (PLIER) to three muscle omic datasets (ATAC-seq, RNA-seq and proteomics) (Fig. 4a). This approach identified 100 Latent Variables (LVs) representing features with shared patterns of expression across three selected omes in response to both EE and RE, with 14 LVs reflecting significant exercise responses (Fig. 4b, Extended Data Fig. 5a). Twelve of the fourteen highlighted LVs were significantly enriched for at least one biological pathway (Fig. 4c, Supplementary Table 5).

**Figure 4:**
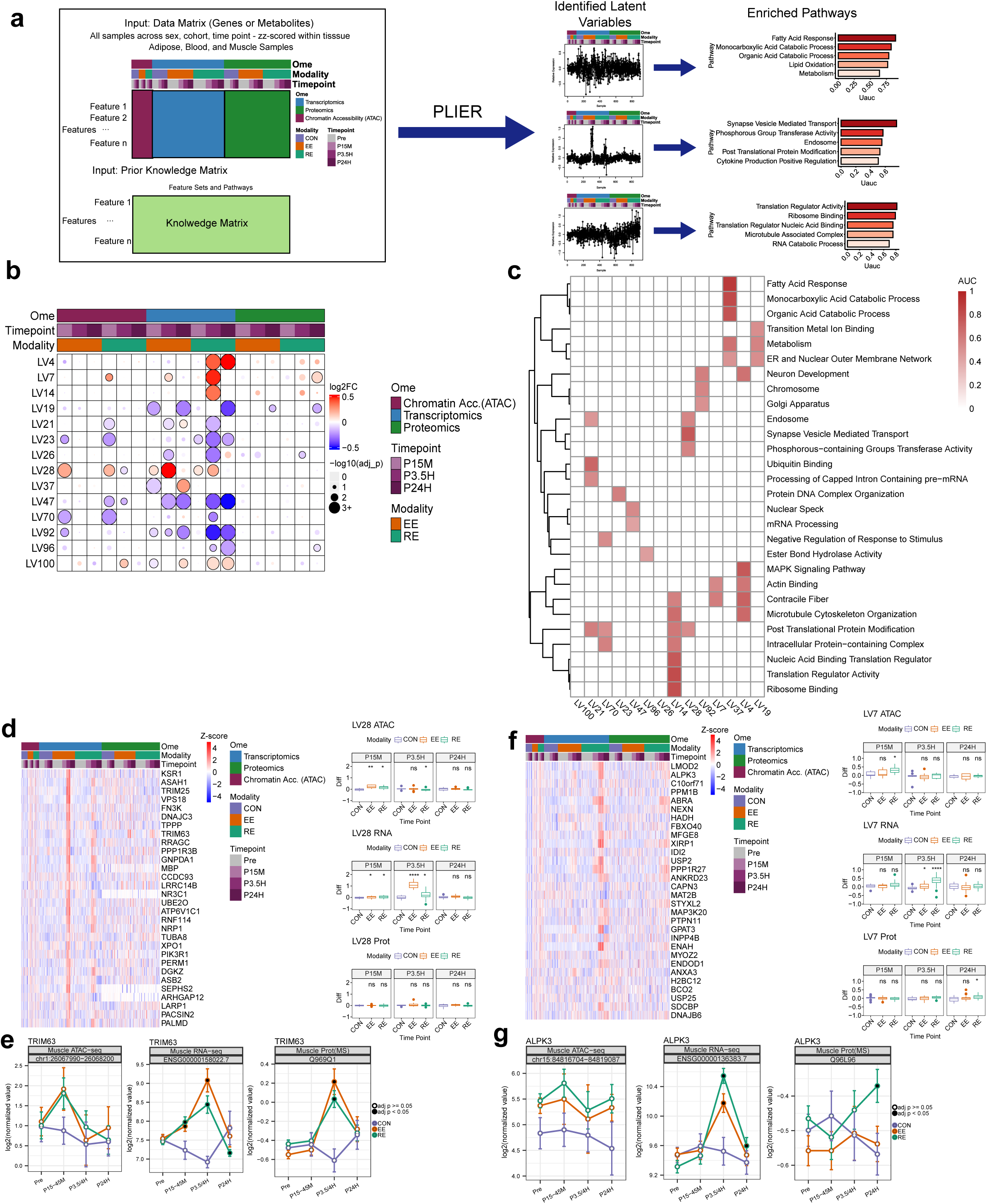
Cross-omic coordinated responses to acute exercise. **a.** Schematic of PLIER method design. PLIER used as input Z-scored normalized muscle ATACseq, RNAseq, and protein abundance data concatenated into a single input matrix where rows were proteins that also had transcriptomic and epigenetic data and columns were the samples in each of the three omes, as well as a matrix of prior knowledge for the inputted feature set. ATACseq data was merged into a cumulative promoter accessibility score for each row. PLIER outputs a series of Latent Variables (LVs) reflecting distinct patterns of behavior across the measured samples with a set of associated features that align with the pattern and specific pathway enrichments based upon the prior knowledge and associated feature set. **b.** Bubble heatmap of the significance (size = adjusted p-value) and directionality (color = log2 fold-change) of the EE or RE response at each time point for each ome described by a subset of LVs with the most significantly identified acute exercise responses. **c.** Top enriched pathways for the LVs described in (B). **d.** Heatmaps and associated boxplots for LV28. Heatmaps indicate within-ome z-scores of the top 30 features for each participant at every ome-time point for a given LV. Each heatmap is paired with boxplots detailing the within-subject expression level difference relative to pre-exercise of each LV in ATAC, RNA, and protein at each post-exercise time point within each modality. Significance is calculated by a t-test measuring EE or RE post- vs pre-exercise differences relative to control subject post- vs pre-exercise differences in LV level. Significance labels represent: ns: p > 0.05, *: 5e-03 < p <= 0.05, **: 5e-04 < p <= 5e-03, ***: 5e-05 < p <= 5e-04, ****: p < 5e-05. **e.** Line plots describing TRIM63 response at the ATACseq (left panel), RNAseq (middle panel) and protein (right panel) abundance levels. TRIM63 ATAC response is highlighted by promoter peak chr1:26067990-26068200. Time points are colored black if differential analysis of a given feature is significant (ad.j P-val < 0.05). **f.** Heatmaps and associated boxplots for LV7. Heatmaps indicate within-ome z-scores of the top 30 features for each participant at every ome-time point for a given LV. Each heatmap is paired with boxplots detailing the within-subject expression level difference relative to pre-exercise of each LV in ATAC, RNA, and protein at each post-exercise time point within each modality. Significance is calculated by a t-test measuring EE or RE post- vs pre-exercise differences relative to control subject post- vs pre-exercise differences in LV level. Significance labels represent: ns: p > 0.05, *: 5e-03 < p <= 0.05, **: 5e-04 < p <= 5e-03, ***: 5e-05 < p <= 5e-04, ****: p < 5e-05. **g.** Line plots describing ALPK3 response at the ATACseq (left panel), RNAseq (middle panel) and protein (right panel) abundance levels. ALPK3 ATAC response is highlighted by promoter peak chr15:84816704-84819087. Time points are colored black if differential analysis of a given feature is significant (adj. P-val < 0.05).

Two LVs demonstrated an apparent coordinated time lag between chromatin accessibility and gene expression. LV28, enriched for endosome and synapse vesicle transport terms, displayed an early increase in accessibility in EE and RE, along with increased gene expression at 15 min and 3.5 hours (Fig. 4d). Top LV28-associated genes included acid ceramidase (*ASAH1)*, consistent with observed decreases in muscle ceramides^27^ (Fig. 3d), and two E3 ubiquitin-protein ligases, *TRIM25* and *TRIM63*. While TRIM25 is known for its involvement in the RIG-1 pathway, sensing viral RNA triggering interferon production,^28^ its role in exercise is less defined. *TRIM63 (MURF1)* is an E3 ubiquitin ligase involved in myosin heavy chain degradation and muscle protein turnover^29^ and a known response gene to both EE and RE.^30^ Accessibility to its promoter region appeared to increase at 15 minutes, but didn’t reach statistical significance, while gene expression increased at early and middle timepoints, and displayed a protein abundance increase at 3.5 hours post exercise, while expression attenuated below baseline levels with RE at 24 hours (Fig. 4e). LV7 displayed an RE-specific response across all three omes, with significant changes in chromatin accessibility at 15 minutes, increases in gene expression at 3.5 hours, and increase in protein abundance at 24 hours (Fig. 4f). LV7 displayed enrichment in actin binding and contractile fiber terms with *LMOD2*, discussed more in the following section, and Alpha Protein Kinase 3 (*ALPK3*) the top-associated genes (Fig. 4g). *ALPK3* is localized to the sarcomere and has been found able to restore impaired contraction in cardiomyopathy^31^.

PLIER was also able to detect time-point variable modality-specific responses to exercise. An EE-specific program was found in LV37 enriched for lipid metabolism (Fig. 4c; Extended Data Fig. 5b), with gene expression decreasing at 15 minutes, but increasing 24 hours post-EE. Genes following this pattern included *HADHA*, *MLYCD*, *CPT1B*, and *ACAA2* (Extended Data Fig. 5c), all involved in FAO, consistent with EE-dominant increases in metabolomic and transcriptomic pathways related to lipid metabolic flux, such as increases in acylcarnitines and FAO transcripts (Fig. 3a,b) ^15,32^. LV4 which showed the second highest correlation between transcript and protein abundance and for gene expression and chromatin accessibility (Extended Data Fig. 5a) displayed enrichment of cytoskeleton and MAPK signaling pathways (Extended Data Fig. 5d). It included genes with RE-specific increases in transcripts and proteins at 3.5 and 24 hours, including ANKRD1, a stress and contraction inducible gene^33^ (Extended Data Fig. 5e). These findings demonstrate the coordinated temporal dynamics of the skeletal muscle exercise response, and identify novel modality conserved and divergent patterns of cross-omic regulatory behavior.

### Exercise-induced HIPK3 dephosphorylation is associated with skeletal muscle remodeling

We employed Post Translational Modification Signature Enrichment Analysis (PTM-SEA) of the phosphoproteome to uncover how early phosphoproteomic changes may regulate subsequent transcriptional remodeling and signaling networks. We observed 52 and 77 significant signatures (adj. p-value < 0.05) across all time points in EE and RE, respectively. The majority of signatures were significant at 15 minutes post-exercise; some were shared between EE and RE (Fig. 5a, Supplementary Table 6, Supplementary Results), while others were EE or RE specific (Extended Fig. 6a).

**Figure 5:**
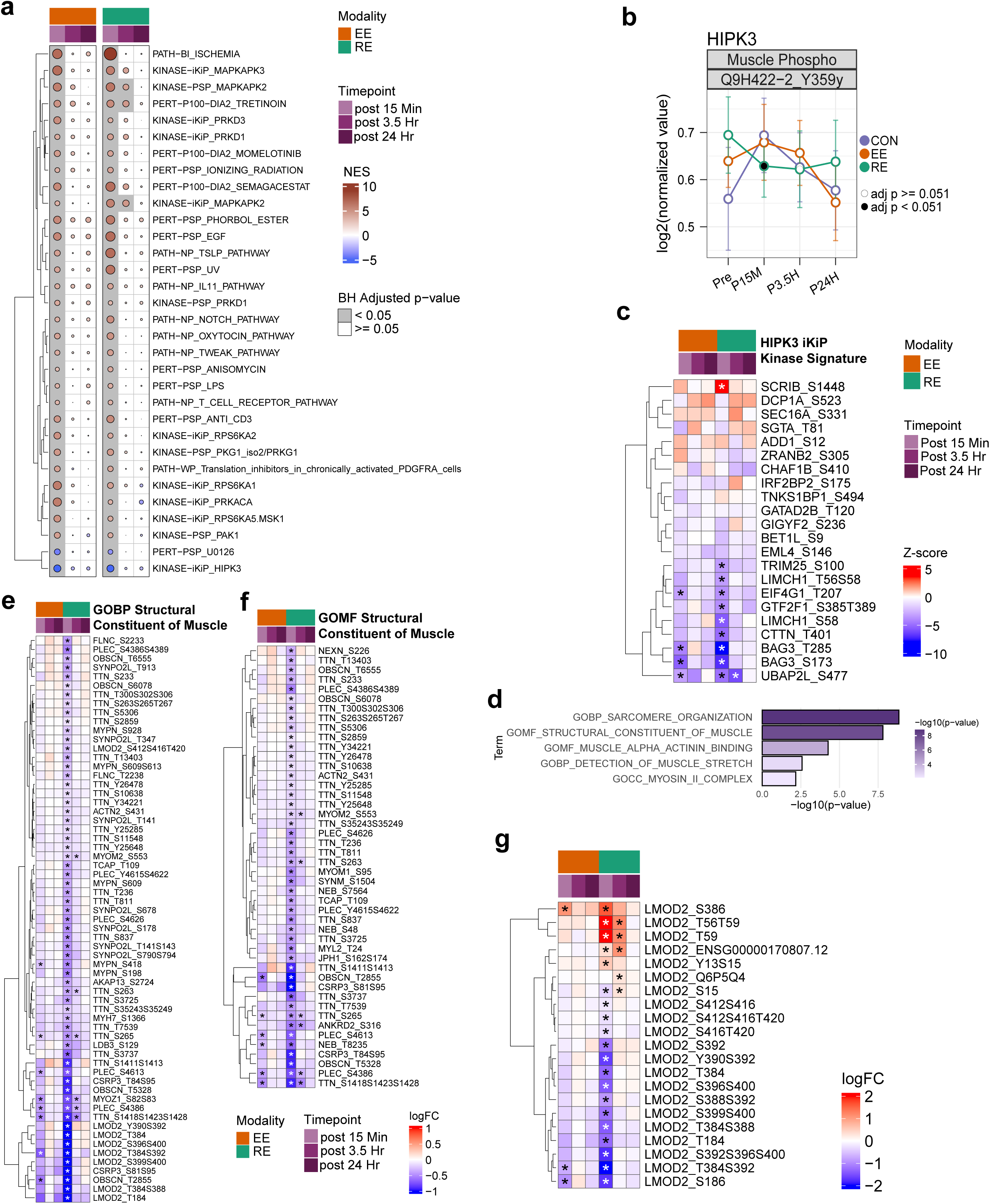
Phosphoproteomics response to acute exercise. **a.** Heatmap of PTM-SEA (Post Translational Modification Signature Enrichment Analysis) of phosphoproteomics dataset. Displayed are selected significantly enriched signatures (adj. P-value < 0.05) at 15 minute time point in both EE and RE. **b.** Temporal trajectory of peptide containing pY359 of HIPK3. EE = Endurance exercise (orange line), RE = resistance exercise (green line), CON = control group (purple), Pre = pre-exercise, D15M =15 minutes post exercise, P3.5H = 3.5 hours post exercise, P24H = 24 hours post exercise. **c.** Heatmap of all the phosphosites in Kinase_iKiP_HIPK3 signature detected in the phosphoproteome dataset. **d.** Bar plot representing HIPK3 pY359 nearest neighbor ORA analysis using C2 and C5 databases. Pathways shown were significant with adj. P-value < 0.05 and contained at least 5 proteins. **e.** Heatmap of GOBP Sarcomere organization pathway, one of the significantly enriched pathways in the ORA of HIPK3 pY359 nearest neighbor analysis. Asterisks indicate significance in the phosphoproteome dataset with adj. P-value < 0.05. **f.** Heatmap of GOMF Structural constituent of muscle pathway, one of the significantly enriched pathways in the ORA of HIPK3 pY359 nearest neighbor analysis. Asterisks indicate significance in the phosphoproteome dataset with adj. P-value < 0.05. **g.** Heatmap of differential LMOD2 (Leiomodin-2) features (transcript, protein and several phosphosites). Asterisks indicate individual feature significance with adj. P-value < 0.05.

As expected, MAPK related phospho-signatures were robustly upregulated with MAPKAPK signatures shared in response to EE and RE, while many others including MAPK3/ERK1, MAPK1/ERK2, MAPK11/p38, MAPK8/JNK1 displayed RE-specific upregulation throughout the post-exercise time course (Extended Data Fig. 6a), pairing with PLIER RE-specific LV4 enrichments (Fig. 4 b,c). Consistent with prior work, acute RE promotes a robust MAPK activation signal to help regulate cell growth and differentiation, stress responses, and the maintenance of muscle mass.^34–37^

Among the most robust and rapid downregulated kinase signatures in both EE and RE was HIPK3 (Fig. 5a). This is a homeodomain-interacting protein kinase belonging to the dual-specificity tyrosine (Y) phosphorylation-regulated kinase (DYRK) family that is autophosphorylated at the tyrosine residue in the activation loop,^38^and involved in transcriptional regulation and apoptosis.^39–41^ Here we detect significant downregulation of Y359, the autophosphorylation site on HIPK3, after RE (Fig. 5b). Leading edge analysis linked this HIPK3 signature to robust dephosphorylation of the molecular chaperone protein BAG3 at S173 and T285 in both EE and RE, suggesting BAG3 phosphorylation is a strong driver of the HIPK3 acute exercise enrichment signature (Fig. 5c, Supplementary Table 6). Transcript levels of BAG3 increased following RE, without observed changes in protein abundance, indicating that the down-regulation of BAG3 phosphorylation is related to phospho-signaling rather than protein abundance (Extended Data Fig. 6b). Further, contraction-induced dephosphorylation of BAG3 facilitates mitophagy, a key component of exercise-induced remodeling to maintain mitochondrial functional efficiency.^42^

To identify additional phosphosites which could be linked to HIPK3 dephosphorylation, we performed a nearest-neighbor analysis by correlating the fold change of all quantified phosphosites across the time points for EE and RE with the fold change of the HIPK3 autophosphorylation site Y359. ORA of the HIPK3 nearest-neighbor phosphosites (Pearson’s *r* ≥0.8), which were also DA for at least one time point highlighted pathways related to striated muscle contraction and sarcomere organization (Fig. 5d, Supplementary Table 6). For example, the Sarcomere Organization and Structural Constituent of Muscle pathways consist of rapidly downregulated phosphosites in response to RE predominantly in titin (TTN) and other muscle structural proteins, suggesting a possible link between HIPK3 activity, exercise-induced mechanical stress, and subsequent structural and functional remodeling of the sarcomere in a manner that is more pronounced with RE (Fig. 5e,f). All the proteins within these pathways displayed decreased phosphorylation at 15 min without changes in global protein abundance and included several sarcomeric proteins such as ACTN2, Myopalladin (MYPN), CSRP3, all localized at the Z-disk, and Leiomodin-2 (LMOD2) concentrated at the pointed ends of actin thin filament with the primary function of elongating and regulating sarcolemmic stability to promote contraction.^43^ Interestingly, this rapid dephosphorylation of LMOD2 (Fig. 5g) precedes its protein upregulation as one of the few differential proteins in RE only (Extended Data Fig. 3c). LMOD2 was also found to be a top feature in the RE-specific cross-omic regulatory LV7 discussed previously and linked to mechanosensitive pathways (Fig. 4f). Moreover, additional PLIER analysis of the combined phosphoproteomics and transcriptomics data identified an LV associated with cytoskeleton organization displaying a set of proteins that included BAG3 and LMOD2 (Extended Data Fig. 6c, Supplementary Table 5). Further CSRP3 phosphorylation was unique to RE. Given its role in facilitating autophagy during myogenesis,^44^ this is consistent with RE-specific transcriptomic enrichments in proteasome activation and myogenic transcripts (e.g. *MYOG*, Extended Data Fig. 4b). Both LMOD2 and CSRP3 displayed consistent de-phosphorylation at 15 minutes post EE and RE, followed by increase of their transcripts at 3.5 hours post-exercise in both modalities, albeit more pronounced and prolonged after RE. Together, our findings highlight several known and novel signal transduction pathways involved in the acute exercise response.

### Central regulatory nodes of the acute exercise response connect with modality divergent transcriptional effects

A key goal of MoTrPAC is to understand the central regulators of exercise responses, including transcription factors (TFs). To this end, we used Spatiotemporal Clustering and Inference of Omics Networks (SC-ION)^45^ to infer multi-omics networks of the acute exercise response and identify regulatory hubs by linking phosphorylation of TFs with the transcriptomic expression of putative downstream target genes (Fig. 6a, Methods). We used changes in protein phosphorylation to assess TF activation, as opposed to gene/protein expression given that phosphorylation occurs rapidly and strongly influences TF activity.^46^ In concert, we used MAGICAL^47^ to infer TF activity from chromatin accessibility (Extended Data Fig. 7a). The regulatory networks inferred by SC-ION and MAGICAL were combined to form the predicted regulatory network for downstream analysis. Using the Network Motif Score (NMS), we identified several TFs as important regulators in the combined EE+RE network (Supplementary Table 7). Several of these included known regulators of exercise-induced skeletal muscle adaptations including MEF2A (Fig. 6b-e; Extended Data Fig 7b; Supplementary Results).

**Figure 6:**
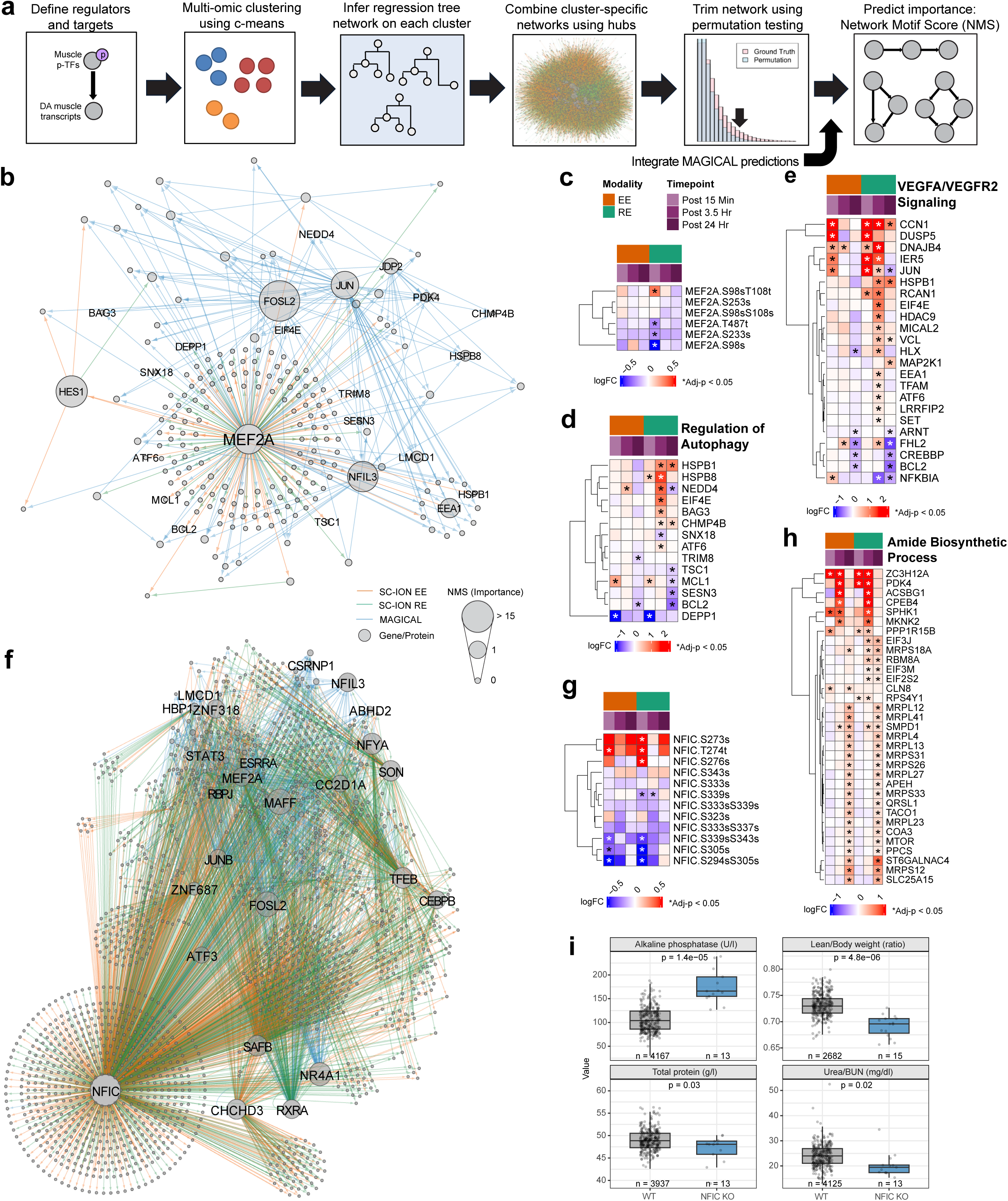
Integrative network analysis identifies NFIC and MEF2A as exercise response hubs. **a.** SC-ION workflow. MAGICAL predictions are integrated into the network prior to calculating the importance of each gene. **b.** Subnetwork of MEF2A and its ChIP-supported, predicted direct targets. TFs with high importance and autophagy-related genes are labeled. In (B) and (E), orange lines were inferred using SC-ION on EE data, green lines were inferred using SC-ION on RE data, and blue lines were inferred using MAGICAL. Node size represents the Normalized Motif Score (NMS; importance). **c.** Heatmap of all MEF2A phosphosites across the time course in response to exercise. In (C), (D), (F), (G), * denotes adjusted p-value < 0.05. **d.** Transcriptome expression of ChIP-supported MEF2A targets which are in the GOBP Regulation of Autophagy gene set. **e.** Transcriptome expression of ChIP-supported MEF2A targets which are in the Reactome VEGFA/VEGFR2 Signaling gene set. **f.** Subnetwork of NFIC and its ChIP-supported, predicted direct targets. TFs with high importance are labeled. **g.** Heatmap of all NFIC phosphosites across the time course in response to exercise. **h.** Transcriptome expression of ChIP-supported NFIC targets which are in the GO-BP Amide Biosynthetic Process gene set. **i.** Phenotypic measures in NFIC knockout mice on a C57/Bl6 background; all data was accessed through the International Mouse Phenotyping Consortium repository, where male and female mice wild-type and knockout mice are included in control and experimental groups, respectively. The p-value between wild-type and NFIC knockout mice is presented above the box plot of each phenotype.

Within our regulatory network analysis, the Nuclear Factor I (NFI) family of transcription factors, particularly NFIC emerged as a novel regulator of the exercise response (Fig. 6f). While the NFI family has suggested roles in regulating tissue development through transcriptional regulation, including myogenesis and atrophy in skeletal muscle,^48,49^ the role of NFIC in regulating exercise responses has not been identified. We filtered ChIP-seq predicted targets of NFIC in the combined network (1580/2326 predicted targets, 67.9%),^50,51^ which included MEF2A, a component of the mitochondrial MICOS complex, CHCHD3, and the estrogen-related receptor, ESRRA, that facilitates mitochondrial biogenesis with exercise training.^52^ A number of phosphosites used to predict NFIC regulation (pS273, pT274, and pS276) increased in response to exercise whereas pS305 and the multiply phosphorylated form pS339pS343 and pS294pS305 decreased (Fig. 6g). ORA on the NFIC targets revealed an enrichment in OXPHOS, catabolism, splicing, and mitochondrial translation and gene translation signatures (Extended Data Fig. 7c). NR4A1 emerged as a top differential NFIC targeted gene strongly induced in response to both EE and RE, which has previously been proposed as a central regulator for physical activity-responsive metabolic programming^4^. The amide biosynthesis signature enriched with both EE and RE included multiple genes related to pan and mitochondrial-localized translation (e.g. *MRPS* genes and *TACO1*) and anabolic processes (e.g. *EIF2,* EIF3, and *MTOR)* (Fig. 6h). EE also induced the unique upregulation of multiple NFIC target genes related to mitochondrial metabolism at 24 hours post-exercise, including multiple electron transport chain complexes (Extended Data Fig. 7d). Conversely, NFIC-targeted genes related to macromolecule catabolic processes more strongly responded to RE, with the strongest changes occurring at 15 min and 3.5 hours (Extended Data Fig. 7e). Within this pathway, several E3 Ubiquitin ligases (*RNF19A, CBLB, FBXL3*) displayed upregulation at 3.5 hours with RE but then attenuated by 24 hours, consistent with an impact of other co-regulators of NFIC-targeted gene expression in driving modality differential transcriptional programs.

Given our findings of NFIC being a central hub of the acute exercise response, we sought to probe data from the international mouse phenotyping consortium^53^ to examine the effects of NFIC deletion on skeletal muscle-related phenotypes. NFIC knockout mice displayed reduced lean to total body mass in males and females (Fig. 6i). Total circulating urea nitrogen and protein levels were also reduced and alkaline phosphatase was elevated in NFIC knockout mice, suggestive of systemically reduced protein synthesis, and pairing with our observation of increases in NFIC targeted genes related to translation, protein synthesis and metabolism.

Overall, our analysis elucidates coordinated phosphorylation and transcriptional regulation of BAG3 by HIPK3, NFIC and MEF2A as a novel candidate autophagy-induced remodeling hub in the acute response to exercise (Extended Data Fig. 7f). Together, these findings highlight known and novel exercise-induced transcriptional networks that largely display modality divergence linked to functional adaptations to exercise training.

### MAPK-driven phosphorylation overrides epigenetic signals to limit E3 ligase expression with RE

FOXO1 and FOXO3 emerged as additional regulatory hubs of the acute exercise response (Fig. 7a). Although the FOXO TF family is classically known for transcriptional regulation of muscle atrophy, it also plays key roles in myofiber regeneration and fiber type switching^54^. Adjacent to FOXO1 in the SC-ION regulatory hub was ZEB1, a putative inhibitor of FOXO3-mediated transcription of *FBXO32* and *TRIM63*, and hence of protein turnover.^55^ To further examine the ZEB1 regulatory hub, ORA of ZEB1 transcriptional targets previously validated by ChIP-seq^50,56^ revealed enrichments of pathways related to stress responses, starvation, autophagy, and transcriptional regulation (Extended Data Fig. 8a, Supplementary Table 8). ZEB1 target genes driving enrichment of the intracellular signal transduction included *MYC*, which increases following mechanical stress to promote muscle remodeling and hypertrophy, and *FOXO1*, with RE prolonging *MYC* expression (Extended Data Fig. 8a). Exploration of the FOXO-ZEB1 signaling axis also revealed modality divergent phosphorylation events. RE resulted in FOXO3 hyperphosphorylation at S425, S413 and S294, while EE only resulted in phosphorylation of S294 (Fig. 7b). Notably all of these sites are located within the FBXO32 transactivation domain, where S294 and S425 phosphorylation promote FOXO3 degradation in response to ERK/p38 MAPK signaling^57^, and S413 phosphorylation is mediated via AMPK,^58^ which is consistent with elevated p38 MAPK signaling (Fig. 3a) and a greater drop in skeletal muscle ATP levels following RE, respectively (Fig. 3e). Further, RE increased ZEB1 phosphorylation at S679, while EE decreased phosphorylation at S315;T324 (Fig. 7c). While the function of these phosphosites has not been identified, ablation of S322 on ZEB1, inhibits its nuclear translocation in response to ERK,^54^ suggesting that these divergent phosphorylation events may drive modality divergent regulation of ZEB1 target genes.

**Figure 7:**
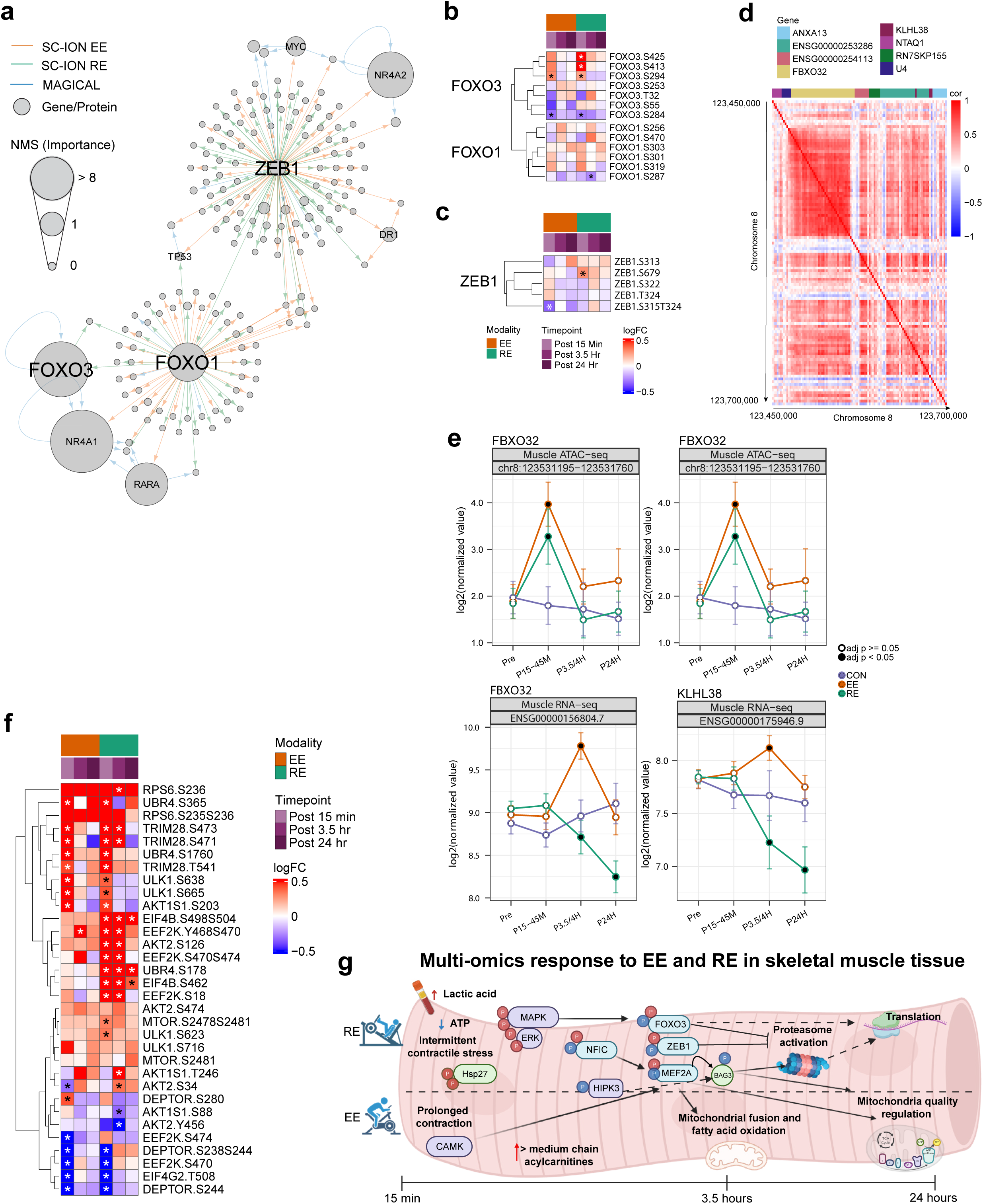
Coordinated Regulation of Muscle Protein Turnover. **a.** Subnetwork of ZEB1, FOXO1, and their ChIP-supported, predicted direct targets. TFs with high importance are labeled. Orange lines were inferred using SC-ION on EE data, green lines were inferred using SC-ION on RE data, and blue lines were inferred using MAGICAL. Node size represents the Normalized Motif Score (NMS; importance). **b.** Heatmap of all FOXO1 and FOXO3 phosphosites across the time course in response to exercise. * denotes adjusted p-value < 0.05. **c.** Heatmap of changes of all ZEB1 phosphosites in response to exercise. * denotes adjusted p-value < 0.05. **d.** Heatmap detailing correlation of chromatin accessibility over all muscle samples for ATACseq peaks on chromosome 8 between base pairs 123,450,000 and 123,700,000. Peaks are annotated by associated gene and gene-body regions. **e.** Temporal changes in genomic accessibility to regions containing the FBXO32 genes (top). Temporal changes in FBXO32 and the adjacent KLHL38 gene expression, where a black dot represents significance at adj. p<0.05. **f.** Heatmap displaying the effects of acute exercise on the phosphorylation of proteins that regulate protein turnover and E3 ligase activity; * represents significance at adj. p<0.05. **g.** Schematic overview of experimental findings displaying divergent acute signaling and downstream molecular multi-omic events following EE and RE in the skeletal muscle. Phosphosite colors represent response to exercise (red: up, blue: down). Dotted line is used to represent whether features are modality divergent, with shared features represented in the center of the pink myofiber (near dotted line). Here, individual differentially abundant features are represented as a circle, and summarized pathway enrichments by CAMERA-PR are written as terms as are metabolite species (e.g. acylcarnitines and lactic acid).

To further probe molecular regulators of protein turnover and remodeling, we examined epigenetic and transcriptional hubs. The most pronounced signal emerged at a genomic locus containing the FOXO-targeted FBXO32, and another regulator of ubiquitin-mediated protein turnover, KLHL38,^59^ where locus accessibility and gene expression were highly correlated (Fig. 7d and Extended Data Fig. 8b). Following both EE and RE, accessibility of this region increased, yet gene expression displayed modality-divergent patterns; *FBXO32* and *KLHL38* expression increased with EE and decreased with RE (Fig. 7e). We further interrogated transcriptional and phosphoproteomic regulators of protein turnover to better characterize modality-divergent signaling events. RE induced early unique protein synthesis phosphosignatures (Fig. 7f), such as RPS6 S236 and EEF2K S470;S474, where RPS6 S236 is phosphorylated in response to ERK/RSK and mTOR signaling.^60^ We additionally observed a strong early MAPK/ERK signal unique to RE likely contributing to repression of the *FBXO32* gene locus via FOXO transcriptional repression (Extended Data Fig. 8c).^57,61^ Together this suggests that the mechanical stress of RE combined with a pronounced drop in ATP (Fig. 3e), promote early anabolic signaling to result in previously-reported RE-specific increases in myofibrillar protein synthesis.^26^ Further, these signals, perhaps in combination with altered ZEB1 phosphorylation patterns, likely drive divergent *FBXO32* and *KLHL38* expression following acute exercise, despite increased gene accessibility, resulting in subsequent increases in skeletal muscle transcriptional activation and remodeling following RE (Extended Data Fig. 8d). Altogether, the divergent mechanical and metabolic demands of EE and RE appear to result in early epigenetic, phosphoproteomic and metabolomic signalling events that drive distinct transcription factor activities and transcriptional programing resulting in modality-divergent biological responses (Fig. 7g).

## Discussion

We have presented comprehensive multi-omics analysis of dynamic molecular responses within skeletal muscle to an acute bout of endurance (EE) or resistance (RE) exercise. Through investigation of shared and divergent molecular responses to EE and RE across time points and omes, we identified novel temporally-related signaling events and regulatory networks as key transducers of the acute exercise response. Overall, EE and RE displayed the greatest molecular overlap at early post-exercise time points, with greater divergence in molecular signatures as recovery progressed. RE induced a greater number of differential metabolites, phosphosites and differentially accessible ATAC peaks at 15 minutes, followed by a greater number of unique differential transcripts and proteins at 3.5 and 24 hours.

While early omic responses were more similar between modalities, RE produced broader and more sustained multi-omic pathway enrichments, encompassing sarcolemmal remodeling, myogenesis, telomere regulation, and translational regulatory pathways. RE also induced a broader suppression of multiple metabolite classes, to include early depletion of acylcarnitines, ATP, CTP and BCAAs. Although *PPARCG1A* expression was comparable between modalities, EE led to a greater induction of mitochondrial fatty acid oxidation transcriptional programs paired with greater triacylglycerol and ceramide depletion, consistent with enhanced mitochondrial gene expression and biogenesis programs following EE^9,62^. Our findings of baseline and acute exercise-induced ome associations with systemic muscle phenotypes highlights the applicability of these findings to understand training adaptations, and warrant future investigation of modality-specific responses.

Exploration of modality divergent signal transduction through phosphoproteomic analysis, revealed early EE-specific CAMK and IL-6 phosphoproteomic signaling activation as well as a more robust induction of AMPKA1 and repression of MTOR signaling (Supplementary Results). These findings paired with molecular indicators of increased systemic lipid oxidation following EE, including mobilization of plasma and skeletal muscle acylcarnitine species^32^. This is also supported by literature demonstrating increased fatty acid oxidation during EE as compared to RE.^63^ Given the peak in mitochondrial gene set enrichments at the transcriptomic level occur at 24hr, this suggests that the time course of proteomic adaptations is beyond 24 hrs, with changes at this timepoint likely being driven by mitochondrial turnover as suggested by others.^64^ These findings provide greater molecular insight into the role of EE on metabolic flexibility and insulin sensitivity.

The acute response to RE was dominated by changes indicative of increased glycolytic metabolism, decreased oxidative flux and stress-associated MAPK/ERK signaling activation. This was concomitant with an increase in proteomic immune cell enrichments and mesenchymal cells in the skeletal muscle at 15 minutes post RE, suggesting an RE-specific damage and remodeling response to the contractile stress. Further, we observed a more robust and prolonged induction of several transcriptomic pathways related to tissue remodeling in the skeletal muscle, including VEGFR signaling, integrin 3 signaling, and hypertrophy, inclusive of myogenic transcripts, following RE. Consistent with our model of greater stress- and hypoxia-induced remodeling following RE, we observed a larger increase in plasma lactic acid with RE (Fig. 2d). These findings are indicative of greater remodeling following RE, emphasizing the utility of this modality in conditions where muscle mass is reduced (e.g. cachexia, sarcopenia, calorie-restricted weight loss).

While MEF2A was observed to be central to both the EE and RE response, a greater number of MEF2A autophagy-related targets were upregulated following RE. Interestingly, these central regulatory hubs were also linked to NFIC-mediated translational events with RE, suggesting coincident degradation and remodeling that was most pronounced following RE. The observation of simultaneous enrichment of pathways and features related to protein degradation and synthesis suggests a complex coordination of molecular remodeling following an acute bout of exercise, particularly with RE. Some responses were also similar across modalities, e.g. increases in chromatin accessibility of *FBXO32* (Atrogin-1), a well-documented regulator of E3 ubiquitin ligase-induced protein turnover.^65^ Despite increased accessibility of this region, RE but not EE resulted in decreased *FBXO32* expression. Through SC-ION analysis, we identified a FOXO3/ZEB1 signaling axis as a potential regulator of divergent FOXO3-mediated gene expression, in addition to RE dominant AMPK and ERK/MAPK regulated FOXO3 phosphorylation events associated with its transcriptional repression. This, combined with phosphorylation events indicative of AKT1S1, MTOR and RPS6 activation, provide a compilation of molecular events resulting in increased translation and likely remodeling following an acute bout of RE. Our findings thus help to define the molecular regulators of simultaneous muscle protein synthesis and breakdown following an acute bout of RE, which is consistent with data from isotope-labeled studies.^25^ These findings also pair with RE-specific proteomic signatures that display simultaneous enrichment in muscle atrophy and developmental terms.

A novel finding of our integrative network analysis is NFIC transcriptional regulation, which has not previously been associated with exercise. While less well studied in the skeletal muscle than its Nuclear Factor I family member, NFIX, NFIC plays central roles in developmental programming through β-catenin/Wnt signaling in adipocytes and chondrocytes,^66^ and has been shown to decrease inflammation in rheumatoid arthritis.^67^ Moreover, we identify the acute dephosphorylation of HIPK2/3 as a novel exercise response, computationally linked to molecular transducers of exercise, such as MEF2A genetic programming and sarcomeric protein post-translational modifications. Prior studies found that HIPK2 acts as a corepressor of MEF2C gene expression in undifferentiated myocytes and dephosphorylation of HIPK2 in the activation loop promotes expression of muscle-specific genes.^41^ Since HIPK2 and HIPK3 share 56% homology in their homeobox-interacting domain that mediates binding to TFs, and MEF2A and MEF2C share over 90% homology in their N-terminal binding domain, we hypothesize that HIPK3 is acting as a transcriptional co-repressor of MEF2A. HIPK3 is downregulated with one acute bout of exercise, leading to de-repression of MEF2A and expression of skeletal muscle genes such as *PPARGC1A*, *NR4A1*, *CKM*, and *CSRP3.* Additionally, our analysis showed a link between HIPK3 activity and phosphorylation of many sarcomelic proteins, one of which was LMOD2 which has been previously shown in muscle,^68,69^ but the function of its phosphorylation is relatively unknown. Given the strong anti-correlation in our dataset between LMOD2 protein and phosphosite expression, it is likely that phosphorylation has a pivotal role in LMOD2 function.

We acknowledge that our analysis has important limitations related to temporal downsampling to limit the number of biopsies performed per participant and omic downsampling due to specific assay limitations and study design (Ref Multi-omic landscape; in preparation/submission). Nonetheless, within participant temporal sampling and the inclusion of a control group better resolves independent exercise effects. Further, while the sample size was among the largest in the field of acute exercise, it was too limited to rigorously assess the effects of sex, age and ethnicity. However, data suggests omic clustering by age, sex and race, which will be investigated in future MoTrPAC studies. Further, the larger number of RE-associated molecular changes should be interpreted cautiously in the context of varying protocol-specific demands on muscle, including greater mechanical loading, intermittent high-force contractions, and likely greater local metabolic stress following RE in sedentary participants.

Overall, this comprehensive multi-omic atlas of acute exercise delineates the temporal orchestration of modality-specific molecular responses and transforms our understanding of how distinct exercise stimuli can be employed to achieve specific long-term metabolic and structural adaptation. Collectively, our findings delineate molecular signaling events that govern acute responses to EE or RE, and provide a framework for understanding modality-divergent adaptations to exercise training.

## Supporting information

Supplementary Results

## Extended Data Figure Legends

**Extended Data Figure 1.**
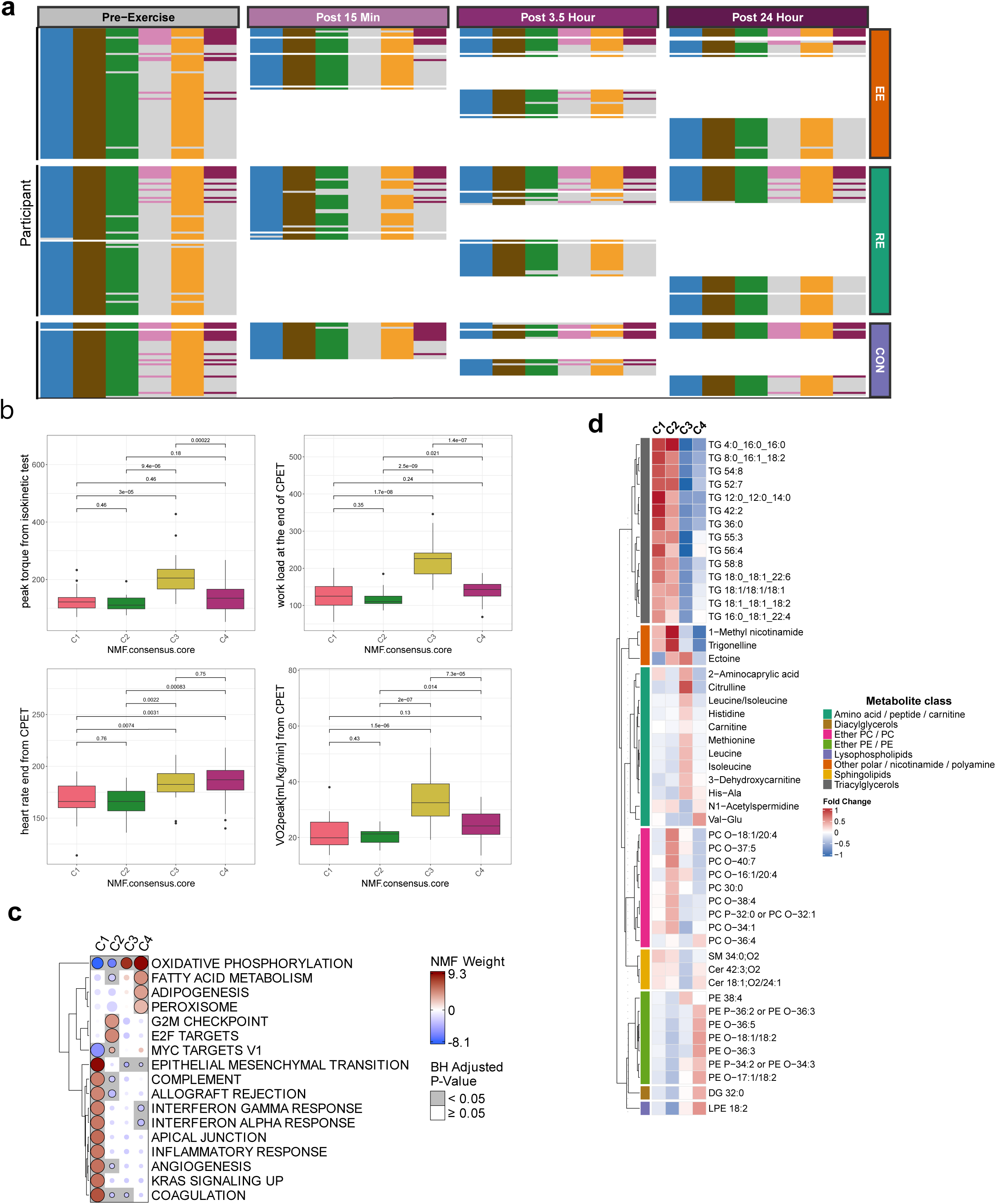
Multi-omics clustering at baseline, related to Figure 1. **a**, Omics Dataset Availability. Each row represents a participant. If data is available for a particular participant for a given tissue-ome-time point combination, this is indicated by a colored cell. A gray cell indicates that no omic data was available or that no sample was collected for that participant at that time point. **b**, Clinical exercise-related measurements from the NMF core samples which are statistically significantly different in at least one cluster. P-values are from pairwise, two-sample t-tests. **c**, CAMERA-PR results on the core samples in each NMF cluster. The input is the multi-omic weight matrix from NMF. The enrichment is run at gene symbol level on the MSigDb Hallmark gene set. Gray squares denote FDR < 0.05 in a one-vs-rest comparison (e.g., C1 denotes C1 vs all other clusters). **d**, Association analysis of metabolomics data using the NMF core samples. Comparisons were performed in a one-vs-rest structure. Plotted metabolites have an FDR < 0.05 across all four comparisons (F-test). Color represents the fold change of that metabolite in each comparison (e.g., a red square in C1-vs-rest means that metabolite is higher in C1 versus all other clusters). Metabolites are clustered by class.

**Extended Data Figure 2.**
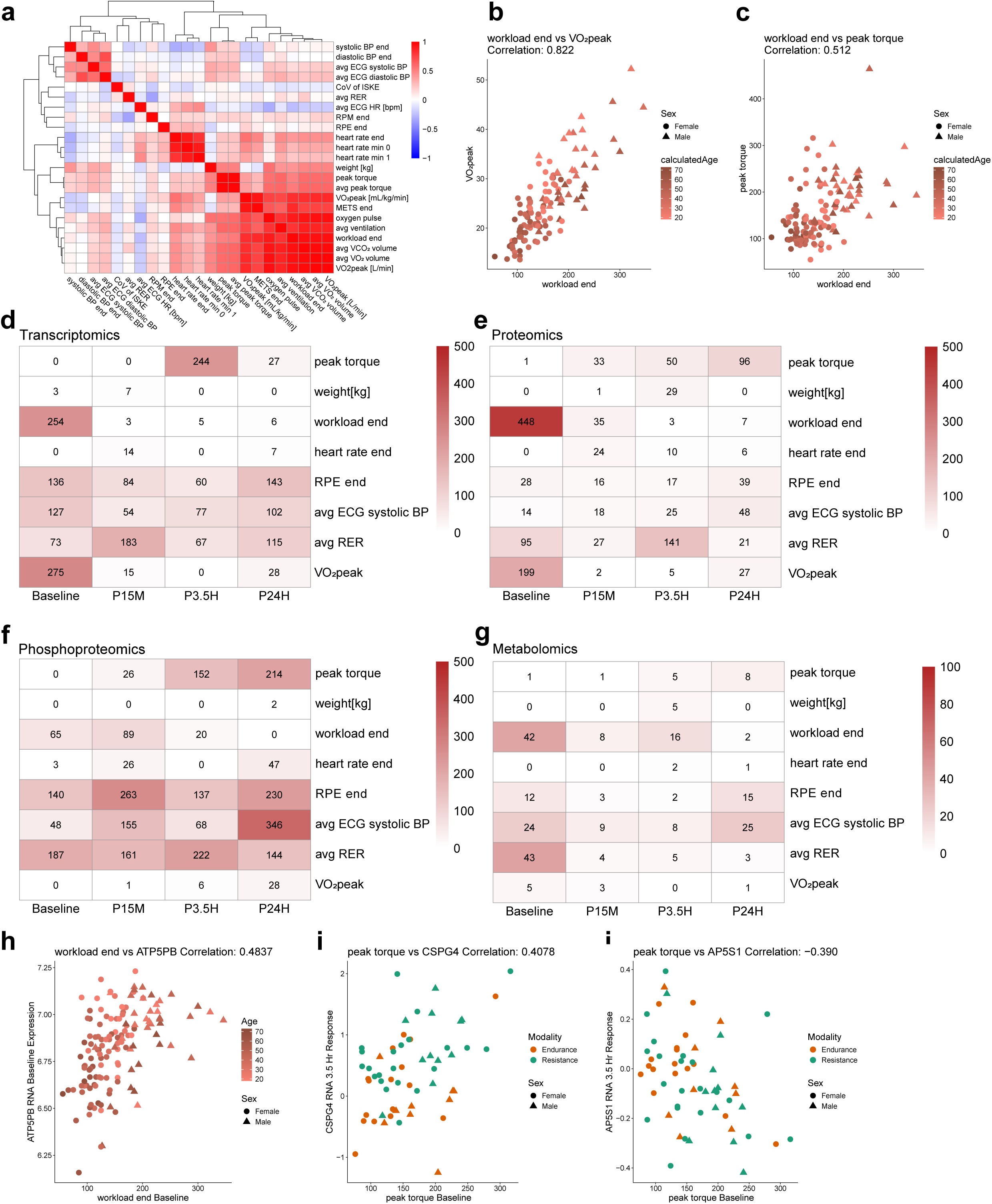
Multi-omic associations with baseline physiological outcomes, related to Figures 1 and 2. **a.** Heatmap of Pearson correlation values between the clinical features included in the clinical x omics comparisons **b.** Scatter plot of screening test workload (Watts) vs screening test VO₂peak, with Pearson correlation included in the figure title. Dots are shaped by sex and colored by age. **c.** Scatter plot of screening test workload (Watts) vs screening test ISKE peak torque, with Pearson correlation included in the figure title. Dots are shaped by sex and colored by age. **d-g**. Heatmaps of the counts of significant clinically-associated (adj. p-val < 0.05) omic features for each linear model comparison in the transcriptomics (D), proteomics (E), phosphoproteomics (F), and metabolomics (G) data. Each row is a distinct clinical feature from the curated clinical screening data of baseline physical fitness for each subject. The first column reflects the model comparing baseline clinical metrics against pre-exercise feature abundance while the remaining three columns reflect the model comparing baseline clinical metrics against feature exercise response at each post-exercise time point. **h.** Scatter plot of screening test workload (Watts) vs pre-exercise gene expression for ATP5PB, with Pearson correlation included in the figure title. Dots are shaped by sex and colored by age. **i-j**. Scatter plots of screening test ISKE peak torque vs 3.5 hour exercise response for genes CSPG4 (I) and AP5S1 (J), with Pearson correlation included in the figure title. Dots are shaped by sex and colored by subject exercise modality.

**Extended Data Figure 3.**
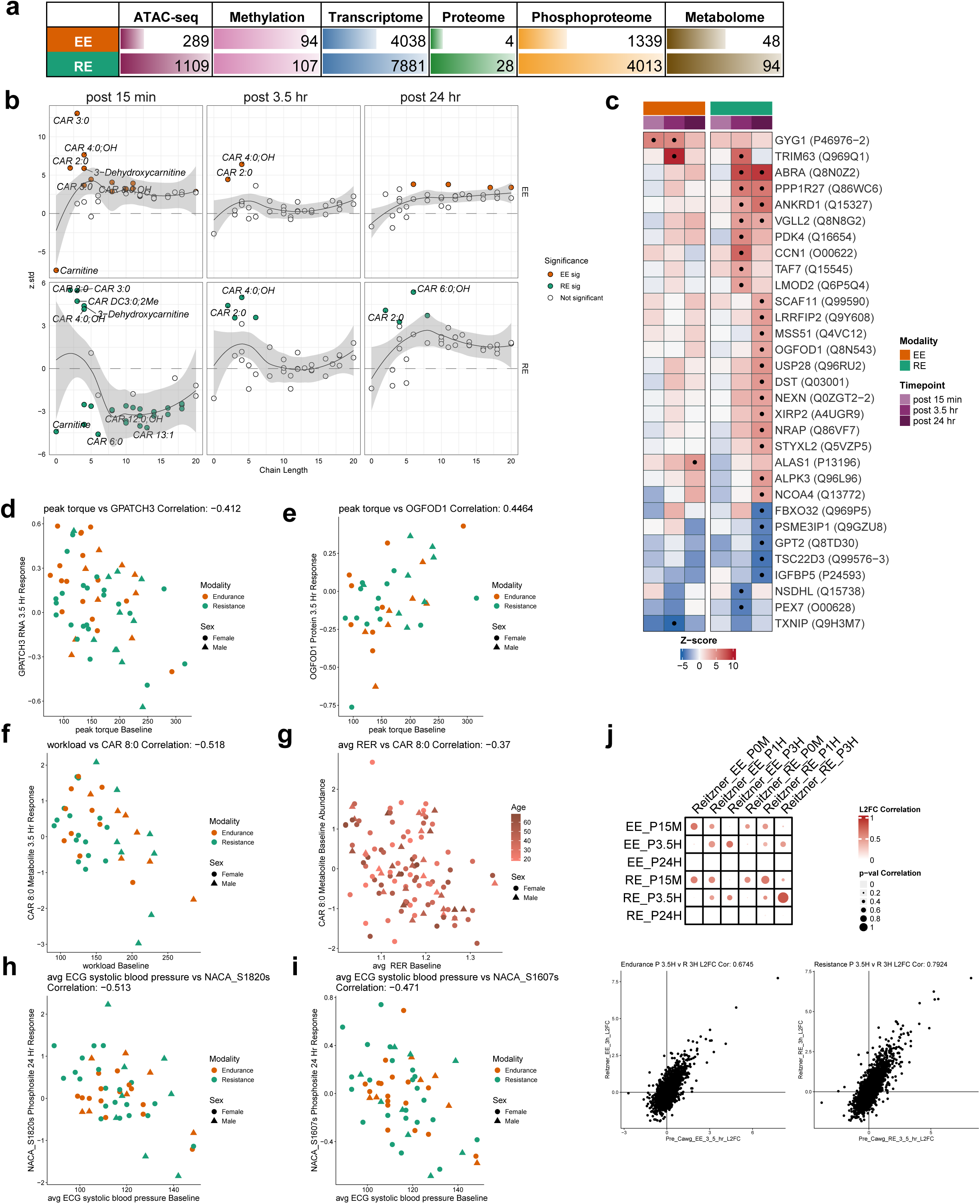
Differential regulation of multi-omics dataset, related to Figure 2. **a.** Total number of differentially abundant (DA) features across the post-exercise timepoints for each ome and exercise modality. Numbers represent the total DA features across the time points; bars are colored using ome-specific colors. **b.** Z-scores of acyl carnitines (CAR) plotted by chain length, shown separately for EE and RE across post-exercise timepoints. **c.** Heatmap displaying temporal dynamics of all DA proteins across post-exercise timepoint (n = 4 in EE, n = 31 in RE). Asterisks indicate significance at adj. P-value < 0.05. **d.** Scatter plot of peak torque from isokinetic test vs 3.5 hour exercise response for gene GPATCH3, with Pearson correlation included in the figure title. Dots are shaped by sex and colored by subject exercise modality. **e.** Scatter plot of peak torque from isokinetic test vs 3.5 hour exercise response for protein OGFOD1, with Pearson correlation included in the figure title. Dots are shaped by sex and colored by subject exercise modality. **f.** Scatter plot of workload at the end of CPET (Watts) vs 3.5 hour exercise response for metabolite CAR 8:0, with Pearson correlation included in the figure title. Dots are shaped by sex and colored by subject exercise modality. **g.** Scatter plot of average respiratory exchange ratio (RER) from CPET vs pre-exercise metabolite abundance for CAR 8:0, with Pearson correlation included in the figure title. Dots are shaped by sex and colored by age. **h.** Scatter plot of systolic blood pressure at the end of CPET vs 24 hour exercise response for phosphosite NACA_S1820s, with Pearson correlation included in the figure title. Dots are shaped by sex and colored by subject exercise modality. **i.** Scatter plot of systolic blood pressure at the end of CPET vs 24 hour exercise response for phosphosite NACA_S1607s, with Pearson correlation included in the figure title. Dots are shaped by sex and colored by subject exercise modality. **j.** Comparison of study transcriptomic results with previous^62^. Top: the bubble heatmap, each grid point describes correlations of L2FC and adjusted p-value between post exercise time points in each study. Size of the bubble reflects Pearson correlation of adjusted p-value of exercise versus control comparisons and the color of the bubble reflects Pearson correlation of L2FC of exercise vs control comparisons. Bottom: scatter plots are also included plotting L2FC of this study versus Reitzner et al study comparing EE vs Control (left) and RE vs Control (right) at the 3.5 hour time point, with Pearson correlations of L2FC values listed above each plot.

**Extended Data Figure 4.**
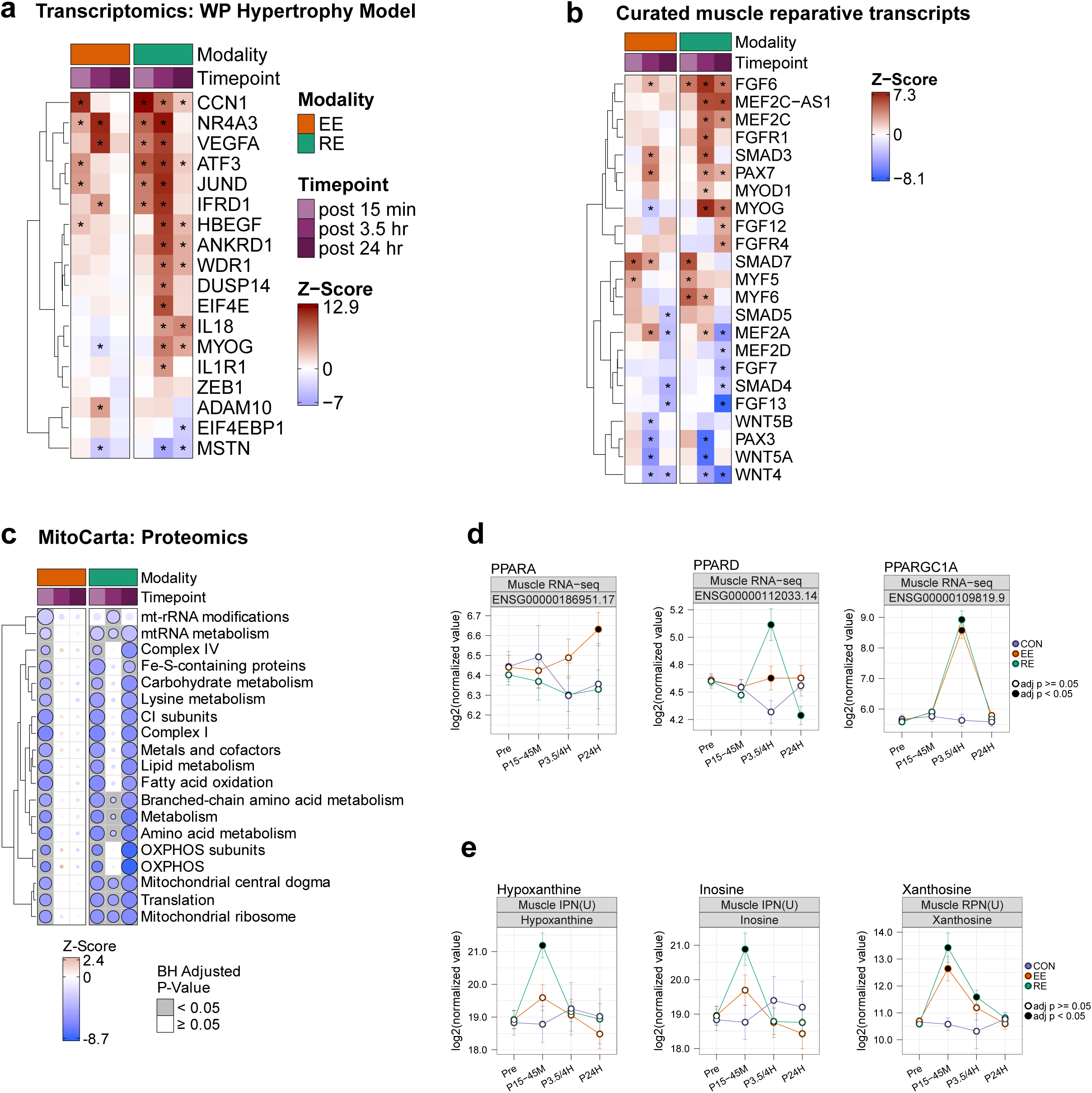
Pathway enrichment analysis, related to Figure 3. **a.** Feature level heatmap of enriched transcripts in the WP Hypertrophy Model pathway, presented in Figure 3a. Asterisks indicate individual feature significance with adj. P-value < 0.05. **b.** Feature level heatmap of curated transcripts related to skeletal muscle regeneration and repair. Asterisks indicate individual feature significance with adj. P-value < 0.05. **c.** Heatmap of CAMERA-PR enrichment of proteomics dataset using MitoCarta 3.0 database displaying significantly enriched terms (adj. P-value < 0.05). **d.** Temporal trajectory of select metabolites in the skeletal muscle. EE = Endurance exercise (orange line), RE = resistance exercise (green line), CON = control group (purple), Pre = pre-exercise, D15M =15 minutes post exercise, P3.5H = 3.5 hours post exercise, P24H = 24 hours post exercise. Significance (adj. P value < 0.05) indicated by black dot per timepoint per group. **e.** Temporal trajectories of PPAR (Peroxisome Proliferator-Activated Receptor) transcripts: PPARA, PPARD, and PPARGC1A. EE = Endurance exercise (orange line), RE = resistance exercise (green line), CON = control group (purple), Pre = pre-exercise, D15M =15 minutes post exercise, P3.5H = 3.5 hours post exercise, P24H = 24 hours post exercise. Significance (adj. P value < 0.05) indicated by black dot per timepoint per group.

**Extended Data Figure 5.**
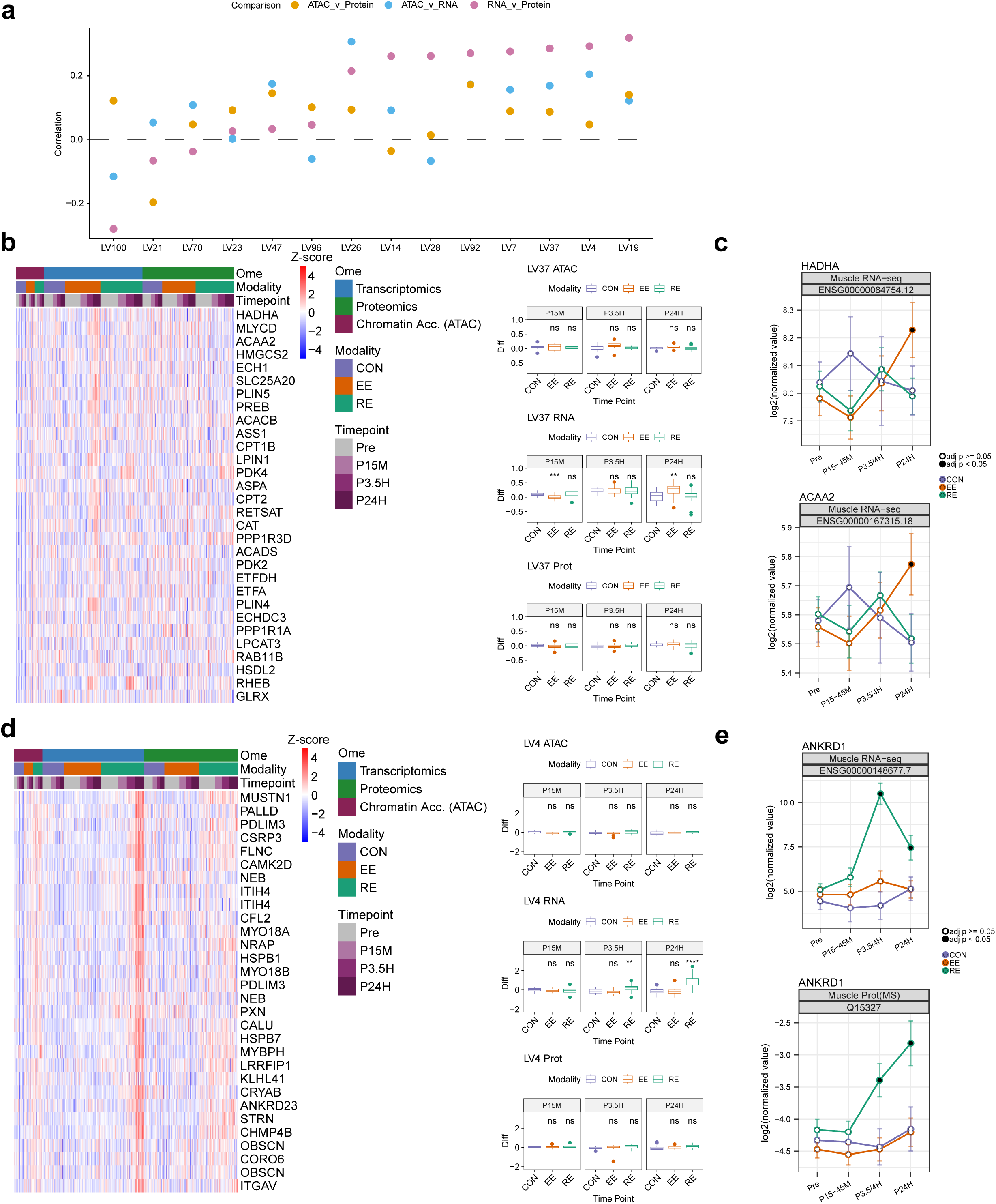
Cross-omic coordinated responses to exercise highlighting changes in protein abundance, related to Figure 4. **a.** The cross-ome Pearson correlation for each of the RNAseq LVs described in (Fig. 4b) **b.** Heatmap and associated boxplot for LV37, where the temporal pattern of omic enrichments are indicated above each panel. Heatmaps indicate within-ome z-scores of the top 30 features for each participant at every ome-time point for a given LV. Each heatmap is paired with boxplots detailing the within-subject expression level difference relative to pre-exercise of each LV in ATAC, RNA, and protein at each post-exercise time point within each modality. Significance is calculated by a t-test measuring EE or RE post- vs pre-exercise differences relative to control subject post- vs pre-exercise differences in LV level. Significance labels represent: ns: p > 0.05, *: 5e-03 < p <= 0.05, **: 5e-04 < p <= 5e-03, ***: 5e-05 < p <= 5e-04, ****: p < 5e-05. **c.** Line plots describing HADHA (top panel) and ACAA2 (bottom panel) response at the RNAseq level. Time points are colored black if differential analysis of a given feature is significant (adj p-val < 0.05). **d.** Heatmap and associated boxplot for LV4, where the temporal pattern of omic enrichments are indicated above each panel. Heatmaps indicate within-ome z-scores of the top 30 features for each participant at every ome-time point for a given LV. Each heatmap is paired with boxplots detailing the within-subject expression level difference relative to pre-exercise of each LV in ATAC, RNA, and protein at each post-exercise time point within each modality. Significance is calculated by a t-test measuring EE or RE post- vs pre-exercise differences relative to control subject post- vs pre-exercise differences in LV level. Significance labels represent: ns: p > 0.05, *: 5e-03 < p <= 0.05, **: 5e-04 < p <= 5e-03, ***: 5e-05 < p <= 5e-04, ****: p < 5e-05. **e.** Temporal trajectory of ANKRD1 (Ankyrin repeat domain 1) transcript expression (top) and protein abundance (bottom). EE = Endurance exercise (orange line), RE = resistance exercise (green line), CON = control group (purple), Pre = pre-exercise, D15M =15 minutes post exercise, P3.5H = 3.5 hours post exercise, P24H = 24 hours post exercise. Y-axis shows Log2 transformed raw means. Significance (adj. P value < 0.05) indicated by black dot per timepoint per group.

**Extended Data Figure 6.**
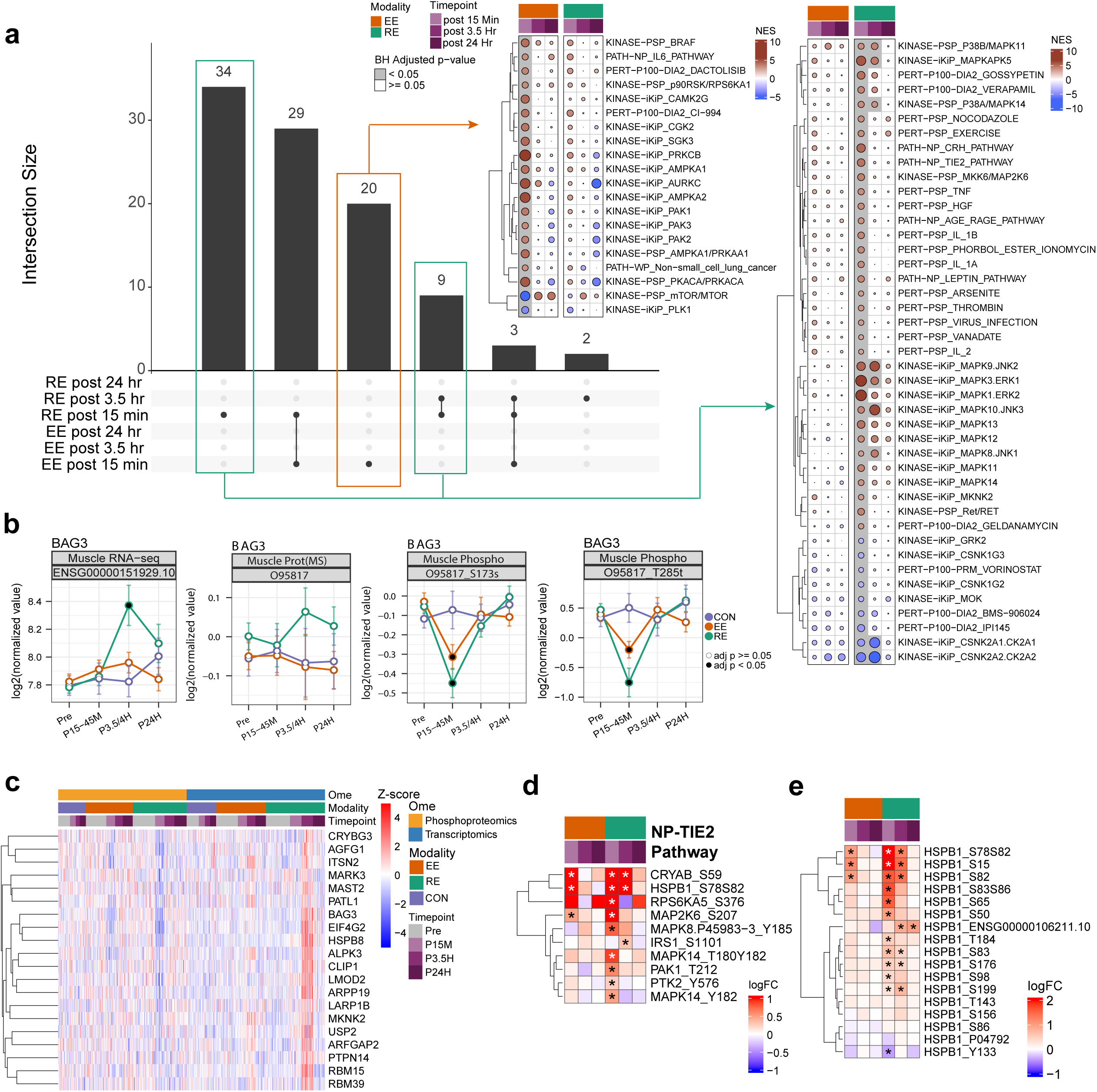
Phosphoproteomics response to acute exercise, related to Figure 5. **a.** Upset plot (left panel) illustrating overlap of significant PTM-SEA (Post Translational Modification Signature Enrichment Analysis) signatures across the exercise modalities and time points. Heatmaps show signatures that are significant only in EE 15 minute time point (middle panel) and only in RE at 15 minute time point and shared between 15 minute and 3.5 hour time points (right panel). **b.** Temporal trajectory of BAG3 (BAG family molecular chaperone regulator 3) transcript, protein and pS173 and pT285 phosphosites. EE = Endurance exercise (orange line), RE = resistance exercise (green line), CON = control group (purple), Pre = pre-exercise, D15M =15 minutes post exercise, P3.5H = 3.5 hours post exercise, P24H = 24 hours post exercise. Significance (adj. P value < 0.05) indicated by black dot per timepoint per group. **c.** Heatmap of one of the LVs in the PLIER analysis with the transcriptome and phosphoproteome. Heatmap indicates within-ome z-scores of the top 15 features for each participant at every ome and time point. **d.** Heatmap of NP-TIE2 pathway. Asterisks indicate significance in the phosphoproteome dataset with adj. P-value < 0.05. **e.** Heatmap of quantified HSPB1 features (transcript and several phosphosites). Asterisks indicate individual feature significance with adj. P-value < 0.05.

**Extended Data Figure 7.**
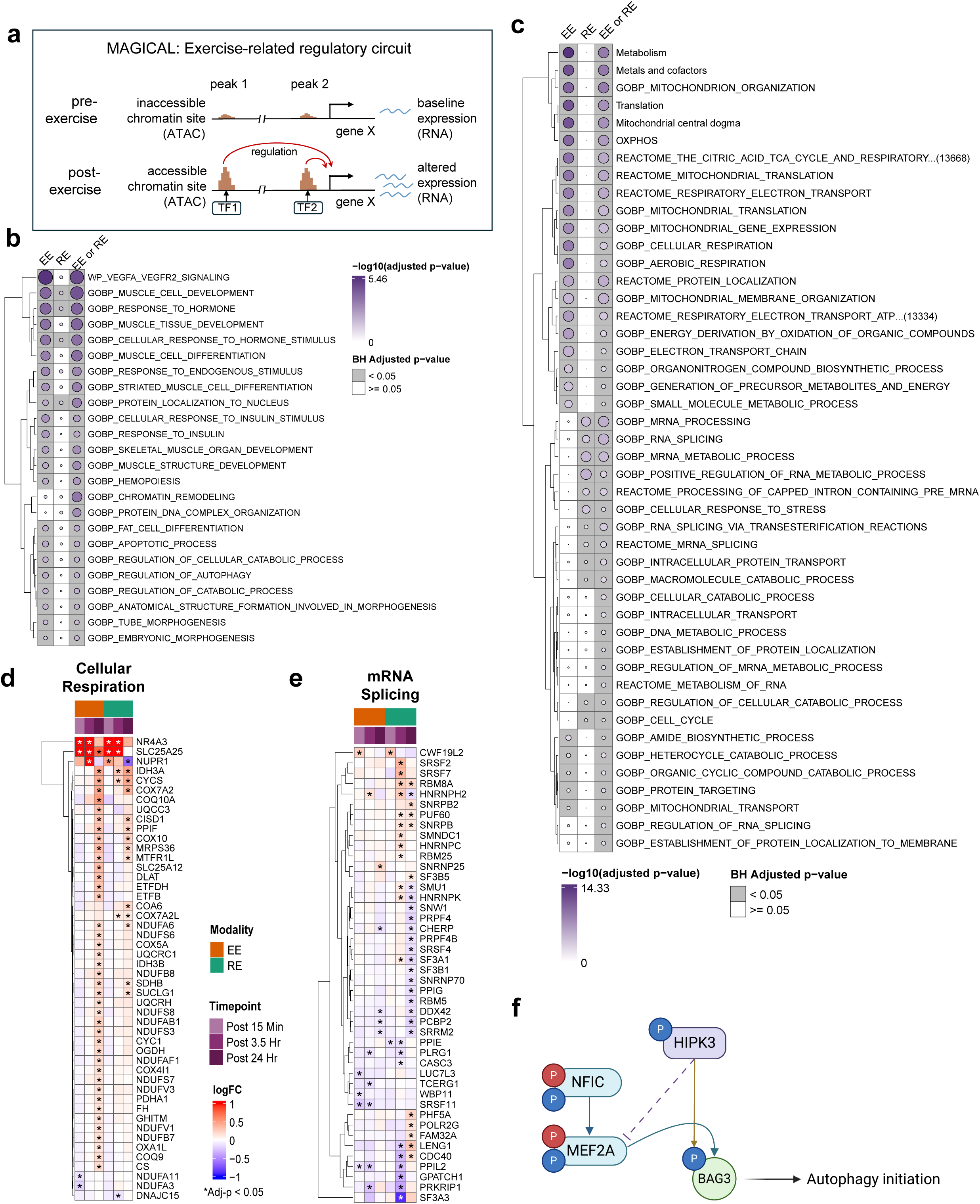
Integrative network analysis identifies NFIC and MEF2A as exercise response hubs, related to Figure 6. **a.** Example of an exercise-regulatory circuit based on ATAC-seq and transcriptome data that would be inferred by MAGICAL. **b.** ORA on ChIP-supported MEF2A targets inferred using EE data, RE data, or either (EE or RE). Circle size and color represent -log_10_(adjusted p-value) and gray squares indicate adjusted p-value < 0.05. **c.** ORA on ChIP-supported NFIC targets inferred using EE data, RE data, or either (EE or RE). Circle size and color represent -log_10_(adjusted p-value) and gray squares indicate adjusted p-value < 0.05. **d.** Transcriptome expression of ChIP-supported NFIC targets which are in the GOBP Cellular Respiration gene set. Asterisks indicate individual feature significance with adj. P-value < 0.05. **e.** Transcriptome expression of ChIP-supported NFIC targets which are in the GOBP Macromolecule Catabolic Process gene set. Asterisks indicate individual feature significance with adj. P-value < 0.05. **f.** Regulatory network between MEF2A, HIPK3, and BAG3. Phosphosite colors represent response to exercise (red: up, blue: down). NFIC transcriptionally regulates MEF2A, which in turn transcriptionally regulates BAG3. In parallel, HIPK3 phosphorylates BAG3. This coordinated regulation is one of many factors that contributes to initiation of autophagy following exercise.

**Extended Data Figure 8.**
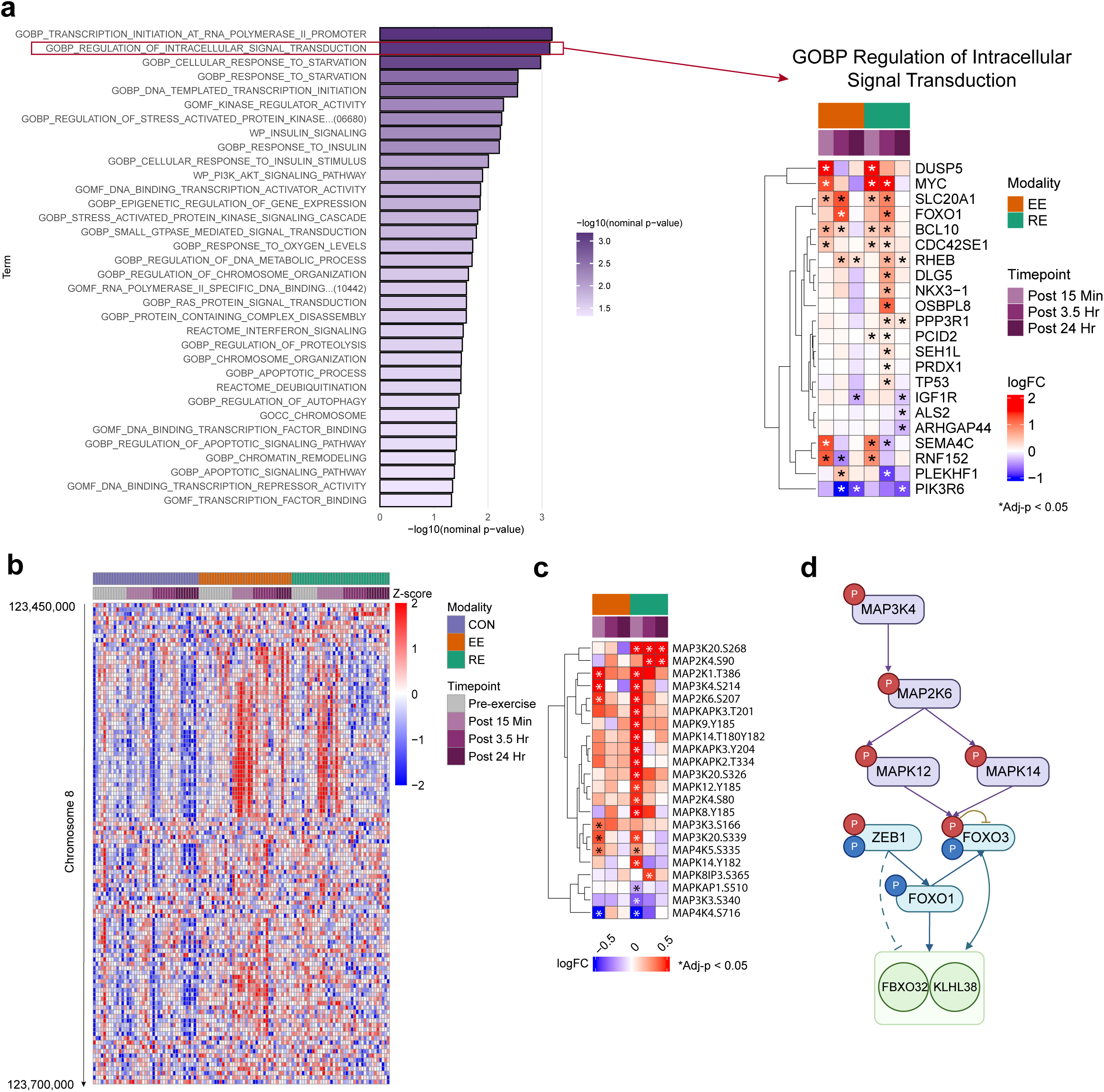
Coordinated regulation of muscle protein turnover, related to Figure 7. **a.** ORA of differentially expressed ChIPseq validated ZEB1 transcriptional targets affected by exercise (left) and exemplary feature level heatmap of GO-BP Regulation of Intracellular Signal Transduction displaying modality shared and divergent ZEB1-targeted gene expression (right); asterisks indicate individual feature significance with adj. P-value < 0.05. **b.** Heatmap of z-scored chromatin accessibility across all muscle samples for ATACseq peaks on chromosome 8 between base pairs 123,450,000 and 123,700,000. Samples are annotated by exercise modality and time point. Highly correlated peaks (as seen in Fig. 7D) annotated to FBXO32 show strong increases in accessibility 15 minutes post EE and RE. **c.** Heatmap of all MAPK phosphosites that are significantly changing (adjusted p-value < 0.05) in at least one time point following acute exercise. **d.** Proposed pathway detailing the regulation of atrophy-linked genes FBXO32 and KLHL38. Phosphosite colors represent response to exercise (red: up, blue: down). MAP3K4 phosphorylates MAP2K6, which phosphorylates MAPK12 and MAPK14, both of which phosphorylate a site on FOXO3 known to inhibit and degrade FOXO3 activity. We identify a ZEB1 to FOXO1 to FOXO3 cascade, all three of which act on FBXO32 and KLHL38. The combination of RE-induced ZEB1 inhibition and the degradation of FOXO3’s enhancement of the target genes leads to the decrease of FBXO32 and KLHL38 expression post RE, despite their increase post EE.

## Supplementary Information

Supplementary Table 1. Participant numbers, related to Figure 1.

Supplementary Table 2. Omics association with baseline physiological features, related to Figures 1 and 2.

Supplementary Table 3. Feature level results, related to Figure 2.

Supplementary Table 4. CAMERA-PR enrichment analysis, related to Figure 3.

Supplementary Table 5. PLIER results, related to Figure 4.

Supplementary Table 6. Phosphoproteome analysis, related to Figure 5.

Supplementary Table 7. SC-ION results, related to Figure 6.

Supplementary Table 8. Protein turnover analysis, related to Figure 7.

## Methods

### Experimental model and study participant details

#### Institutional Review Board

The Molecular Transducers of Physical Activity Consortium (MoTrPAC) (NCT03960827) is a multicenter study designed to isolate the effects of structured exercise training on the molecular mechanisms underlying the health benefits of exercise and physical activity. Methods are described here in sufficient detail to interpret results, with references to supplemental material and prior publications for detailed descriptions. The present work includes data from prior to the suspension of the study in March 2020 due to the Covid-19 pandemic. The study was conducted in accordance with the Declaration of Helsinki and approved by the Institutional Review Board of Johns Hopkins University School of Medicine (IRB protocol # JHMIRB5; approval date: 05/06/2019). All MoTrPAC participants provided written informed consent for the MoTrPAC study indicating whether they agreed to open data sharing and at what level of sharing they wanted. Participants could choose to openly share all de-identified data, with the knowledge that they could be reidentified, and they could also choose to openly share limited de-identified individual level data, which is lower risk of re-identification. All analyses and resulting data and results are shared in compliance with the NIH Genomic Data Sharing (GDS) policy and Data Safety Monitoring Board (DSMB) requirements for the randomized study.

#### Participant Characteristics

Volunteers were screened to: (a) ensure they met all eligibility criteria; (b) acquire phenotypic assessments of the study population; and (c) verify they were healthy and medically cleared to be formally randomized into the study (^70^).^71^ Participants were then randomized into intervention groups stratified by clinical site and completed a pre-intervention baseline acute test. A total of 176 participants completed the baseline acute test (EE=66, RE=73, and CON=37), and among those, 175 (99%) had at least one biospecimen sample collected (adipose, blood, muscle) and 174 had muscle tissue collected.

#### Participant Assessments

As described in *Clinical Landscape and protocol paper,*^14^ prior to randomization, participant screening was completed via questionnaires, and measurements of anthropometrics, resting heart rate and blood pressure, fasted blood panel, cardiorespiratory fitness, muscular strength, and free-living activity and sedentary behavior.(ref clinical landscape)^71^ Cardiorespiratory fitness was assessed using a cardiopulmonary exercise test (CPET) on a cycle ergometer (Lode Excalibur Sport Lode BV, Groningen, The Netherlands) with indirect calorimetry. Quadriceps strength was determined by isometric knee extension of the right leg at a 60° knee angle using a suitable strength device.^71^ Grip strength of the dominant hand was obtained using the Jamar Handheld Hydraulic Dynamometer (JLW Instruments, Chicago, IL). See *Clinical Landscape* for participant assessment results.

#### Acute Exercise Intervention

The pre-intervention baseline EE acute bouts (Fig. 1a) were composed of three parts: (1) a 5min warm up at 50% of the estimated power output to elicit 65% VO_2_peak, (2) 40min cycling at ∼65% VO_2_peak, and (3) a 1min recovery at ∼25 W. The pre-intervention baseline RE acute bouts were composed of a 5min warm-up at 50-60% heart rate reserve (HRR) followed by completion of three sets each of five upper (chest press, overhead press, seated row, triceps extension, biceps curl) and three lower body (leg press, leg curl, leg extension) exercises to volitional fatigue (∼10RM) with 90sec rest between each set, with an isokinetic leg lift as the last exercise before biopsy. Participants randomized to CON did not complete exercise during their acute tests. Participants rested supine for 40min to mirror the EE and RE acute test schedule. See *Clinical Landscape* for acute bout results.

#### Biospecimen Collection

To standardize conditions prior to the acute test, participants were instructed to comply with a variety of controls related to COX-inhibitors, biotin, caffeine, alcohol, exercise, and final nutrition consumption.^71^ Muscle samples were collected for the acute test at specific time points before and after exercise (Fig. 1a).

Participants arrived fasted in the morning, and rested supine for at least 30min prior to pre-exercise biospecimen collections. re-exercise biospecimens were obtained from all participants.The subsequent post-exercise collection time points varied for each training group and their randomized temporal profile (early, middle, late, all). Muscle tissue samples were collected at 15 minutes (early), 3.5 hours (middle) and 24 hours (late) post exercise time points.The post-exercise time point began at the completion of the 40 minutes of cycling, third set of leg extension, or 40 minutes of rest, depending on the randomized group. Participants rested supine or seated for biospecimen collections. The collection and processing protocols for each tissue were previously described in the human clinical protocol design paper.^71^ See *Clinical Landscape* for biospecimen collection success results^70^.

The number of biospecimen samples that were profiled on any given assay varied by time point, and randomized intervention group for feasibility reasons. Multi-omic coverage is summarized in Extended Data Fig.1a and Supplementary Table 1.

### Molecular Quantification Methods

#### ATAC-sequencing

ATAC-seq (Assay for Transposase Accessible Chromatin using sequencing) assays of muscle tissue were done at Icahn School of Medicine at Mount Sinai. Muscle tissue samples were processed for ATAC-seq by study group (CON, EE, and RE). Eight to 12 samples were processed at a time for nuclear extraction and library preparation, and all the samples in the study group were sequenced together. Nuclei from aliquoted tissue samples (10 mg for Muscle tissues) were extracted using the Omni-ATAC protocol with modifications. Two tissue-specific consortium reference standards were included for sample processing QC. 500-600 ul 1X homogenization buffer with protease inhibitor was added to the sample micronic tubes plus three 1.8 mm ceramic bead media. Tubes were vortexed for 5-10 seconds 3 times and kept on ice. The mixture was homogenized using Bead Ruptor Elite (Bead Mill Homogenizer) and Omni BR Cryo Cooling Unit at the settings: Tube 1.2 ml, Speed: 1.0, Cycles: 2, Time: 20 Sec, Dwell Time: 20 Sec, Temperature: 4-8⁰C. The homogenized mixture was filtered with Mini 20 um Pluristrainer and waited for 20-30 seconds. The homogenate tubes were centrifuged with filters attached briefly if the homogenate did not pass through the filter easily. Nuclei were stained with Trypan blue dye 0.4% and counted using Countess II Automated Cell Counter. Depending upon the counts, the number of nuclei to use for transposition was calculated. For Human Pre-covid muscle samples, 250K-300K nuclei were used. Calculated nuclei were added to 1 ml RSB in 2.0 ml Eppendorf tubes, mixed well and centrifuged at 510 RCF for 10 min on fixed angle centrifuge at 4⁰C. 70% supernatant was removed without disturbing pellet at the bottom and nuclei were again centrifuged at 510 RCF for 5 minutes. Almost all of the supernatant was removed. A 50 μl transposition mix was added to the pellet, and mixed by pipetting 6 times, then incubated at 37⁰C on a thermomixer with 1000 rpm for 30 minutes and immediately transferred to ice after incubation. The transposed DNA was purified using Zymo DNA Clean and Concentrator kit (Zymo research, D4014). The DNA product was amplified using NEBnext High-Fidelity 2x PCR Master Mix (NEB, M0541L) and custom indexed primers. Purified PCR libraries were amplified using Ampure XP beads, double-sided bead purification (sample to beads ratios 0.5X and 1.3X).

#### ATAC-seq library sequencing and data processing

Pooled libraries were sequenced on an Illumina NovaSeq 6000 platform (Illumina, San Diego, CA, USA) using a paired-end 100 base-pair run configuration to a target depth of 25 million read pairs (50 million paired-end reads) per sample. Reads were demultiplexed with bcl2fastq2 (v2.20.0) (Illumina, San Diego, CA, USA). Data was processed with the ENCODE ATAC-seq pipeline (v1.7.0) (https://github.com/ENCODE-DCC/atac-seq-pipeline). Samples from a single sex, group and exercise time point, e.g., endurance exercising males 15 minutes post exercise bout, were analyzed together as biological replicates in a single workflow. Briefly, adapters were trimmed with cutadapt (v2.5) and aligned to human genome hg38 with Bowtie 2 (v2.3.4.3).^72^ Duplicate reads and reads mapping to the mitochondrial chromosome were removed. Signal files and peak calls were generated using MACS2 v2.2.4,^73^ both from reads from each sample and pooled reads from all biological replicates. Pooled peaks were compared with the peaks called for each replicate individually using Irreproducibility Discovery Rate and thresholded to generate an optimal set of peaks. The cloud implementation of the ENCODE ATAC-seq pipeline and source code for the post-processing steps are available at https://github.com/MoTrPAC/motrpac-atac-seq-pipeline. Optimal peaks (overlap.optimal_peak.narrowPeak.bed.gz) from all workflows were concatenated, trimmed to 200 base pairs around the summit, and sorted and merged with bedtools v2.29.0 to generate a master peak list.^74^ This peak list was intersected with the filtered alignments from each sample using bedtools coverage with options -nonamecheck and -counts to generate a peak by sample matrix of raw counts. The remaining steps were applied separately on raw counts from each tissue. Peaks from non-autosomal chromosomes were removed, as well as peaks that did not have at least 10 read counts in six samples, which aided in increasing significant peak output in differential analysis. Principal component analysis (PCA) was conducted to find sample outliers to be excluded from downstream analysis using the *call_pca_outliers* command (found in the https://github.com/MoTrPAC/MotrpacRatTraining6mo/ R package).^75^ No PCA outliers were identified. Filtered raw counts were then quantile-normalized with variancePartition::voomWithDreamWeights,^76^ which is akin to limma::voom,^77^ with the added ability to include random effects in the formula. Two samples were flagged as potential sample mislabels based on Somalier analysis^78^ and were removed from downstream analysis. See Supplementary Results of the landscape manuscript ^15^ for an overview of experimental limitations in the muscle ATAC-seq data.

#### Methylation Capture sequencing

##### Data production

Plates for methylation capture were batched for methylation capture processing by exercise group. Samples were processed in subject-level batches of 4 to make 1 library pool, and all timepoints for a subject were pooled together when possible. Library pools (4 samples) were randomized into sequencing pools (40 total samples) to distribute the study groups.

Genomic DNA from muscle was extracted using the GenFind v3 kit (Beckman Coulter) in an automated workstation (Biomek Fxp) according to the manufacturer’s instructions. Tissue specific control samples (provided by the Consortium) were included to monitor the sample processing QC. The DNA was quantified by Qubit assay (dsDNA BR assay, Thermo Fisher Scientific) and the quality was determined by the Nanodrop A260/280 and A260/230 ratios. A260/280 and A260/230 ratios were used to determine the quality of the DNA. DNA samples with a A260/A230 ratio above 1.8 were used for the assay.

The Methyl-Cap sequencing libraries were prepared using the Illumina TruSeq Methyl Capture EPIC Library Prep Kit (Cat# FC-151-1003) according to the manufacturer’s instructions (https://www.illumina.com/products/by-type/sequencing-kits/library-prep-kits/truseq-methyl-capture-epic.html). Briefly,1 μg of genomic DNA in 50 μL was fragmented using a Covaris E220 ultrasonicator (Covaris, USA) and the fragment size was assessed by running 1.0 μL of the samples on a Bioanalyzer High Sensitivity DNA chip (Agilent Technologies). Samples with DNA fragments between 100-300 bp with a peak between 155–175 bp were used for library preparation. Fragmented DNA was end repaired into dA-tailed fragments and ligated with custom xGen™ Methyl UDI-UMI Adapters (Integrated DNA Technologies). Adapter-ligated DNA was purified by magnetic bead separation. The DNA samples were then pooled in groups of 4 in equal volume (10 μL/samples, 40 μL final volume) and hybridized with Illumina EPIC probe sets (covering >3.3 million targeted CpG sites). The DNA fragments hybridized with probes were captured by streptavidin-magnetic beads and purified. The enriched DNA underwent a second round of hybridization and purification to ensure high specificity of the captured regions. Bisulfite conversion was performed on the captured DNA libraries and PCR amplified. The purified libraries were analyzed with the 2100 Bioanalyzer High Sensitivity DNA assay (Agilent Technologies) and quantified by Qubit assay (ThermoFisher). The libraries contained fragment size between 230 - 450 bp with a peak around 250 - 300 bp.

The libraries were pooled and 15% PhiX DNA (Illumina) was added to each library pool to boost diversity. DNA sequencing of paired-end reads (2 x 100 bp reads) were performed on an Illumina Novaseq 6000 system (Illumina Inc.) at a depth of approximately 60 million reads per library. In order to capture the 9-base UMIs, libraries were sequenced using 17 cycles for i7 index read and 8 cycles for the i5 index read.

#### Methylation data processing, quantification and QC

The 17 cycles for the i7 index read in the libraries consist of 8 cycles of sample i7 barcode index and 9 cycles of UMI. Using the mask options --use-bases-mask Y*,I8Y*,I*,Y* --mask-short-adapter-reads 0 --minimum-trimmed-read-length 0 in bcl2fastq command to extract the two read (R1,R2) FASTQ files, and index FASTQ files that contain the UMI information. The three FASTQ files are then fed into the methylcap processing pipeline as described in https://github.com/MoTrPAC/motrpac-methyl-capture-pipeline. Briefly, the pipeline first attaches the 9 cycles of the UMI index FASTQ file as part of the read names in both R1 and R2 FASTQ files and then uses trimgalore (via cutadapt v1.18)^79^ to remove the adapters. The FASTQ files are aligned with bismark (v0.20.0)^80^ and the resulting bam files are deduplicated using the bismark command deduplicate_bismark with option “--barcode”. The bismark coverage from the adjacent neighboring positions of the two opposite strands are merged with the bismark command coverage2cytosine with the option “--merge_CpG”.

NGScheckmate (v1.01) was applied to the deduplicated bam files from the Bismark pipeline to check the within-subject genotype consistency.^81^ The percentage of reads mapped to chrX and chrY are also checked against with the sex annotation of the sample. The inconsistent samples from genotype and sex checking and bismark QC metrics (two low read depth, two high %CHH) outliers were removed from further analysis.

The bismark coverage from all samples are combined into a single R data file after removing CpG sites with 5X coverage in less than 50% of the samples. To merge highly correlated nearby sites into a region, Markov Cluster Algorithm (MCL) is then applied on the bismark coverage R data file for each tissue separately,^75,82,83^ with the following decision algorithm: a) For each sample, compute nCpG10, the number of CpG sites with at least 10X coverage. b) Filter out samples with less than 20 percentile of the nCpG10. c) apply MCL on the CpG sites with at least 10X coverage on all of the remaining samples. d) also apply the CpG merging results from c) to the filtered out samples in b).

Principal component analysis (PCA) was conducted to find sample outliers to be excluded from downstream analysis using the *call_pca_outliers* command (found in the https://github.com/MoTrPAC/MotrpacRatTraining6mo/ R package). No additional outliers were detected from PCA in this ome.

#### Methylation Statistical analysis

To accommodate the within subject mixed effects model for the differential methylation analysis, the coverage of all of the CpG sites (or CpG regions with merged CpG sites obtained from the MCL) is applied to Malax to obtain the p-value.^84^ The statistical comparison of interest compares the change from pre-exercise to a during or post-exercise timepoint in one of the two exercise groups (RE and EE) to the same change in control. Due to the limited sample sizes in the methylation data, only sex and age are the additional covariates in the statistical model. In order to reduce the computational time, the CpG sites/regions are divided into 100 small data sets for the Malax analysis using cluster computers for parallel computing and the adjusted p-values are computed on the final merged dataset. After hypothesis testing using Malax, the change in absolute methylation percentage is calculated using edgeR.^85^

#### Transcriptomics

RNA Sequencing (RNA-Seq) was performed at Stanford University and the Icahn School of Medicine at Mount Sinai. Processing randomization was done according to https://github.com/MoTrPAC/clinical-sample-batching.

#### Extraction of total RNA

Tissues (∼10 mg for muscle) were disrupted in Agencourt RNAdvance tissue lysis buffer (Beckman Coulter, Brea, CA) using a tissue ruptor (Omni International, Kennesaw, GA, #19-040E). Total RNA was extracted in a BiomekFX automation workstation according to the manufacturer’s instructions for tissue-specific extraction. Tissue-specific consortium reference standards were included to monitor the sample processing QC. The RNA was quantified by NanoDrop (ThermoFisher Scientific, # ND-ONE-W) and Qubit assay (ThermoFisher Scientific), and the quality was determined by either Bioanalyzer or Fragment Analyzer analysis.

#### mRNA Sequencing Library Preparation

Universal Plus mRNA-Seq kit from NuGEN/Tecan (# 9133) were used for generation of RNA-Seq libraries derived from poly(A)-selected RNA according to the manufacturer’s instructions. Universal Plus mRNA-Seq libraries contain dual (i7 and i5) 8 bp barcodes and an 8 bp unique molecular identifier (UMI), which enable deep multiplexing of NGS sequencing samples and accurate quantification of PCR duplication levels. Approximately 500ng of total RNA was used to generate the libraries for muscle. The Universal Plus mRNA-Seq workflow consists of poly(A) RNA selection, RNA fragmentation and double-stranded cDNA generation using a mixture of random and oligo(dT) priming, end repair to generate blunt ends, adaptor ligation, strand selection, AnyDeplete workflow to remove unwanted ribosomal and globin transcripts, and PCR amplification to enrich final library species. All library preparations were performed using a Biomek i7 laboratory automation system (Beckman Coulter). Tissue-specific reference standard provided by the consortium was included with RNA isolations to QC the RNA.

#### RNA Sequencing, quantification, and normalization

RNA sequencing, quantification, and normalization Pooled libraries were sequenced on an Illumina NovaSeq 6000 platform (Illumina, San Diego, CA, USA) to a target depth of 40 million read pairs (80 million paired-end reads) per sample using a paired-end 100 base pair run configuration. In order to capture the 8-base UMIs, libraries were sequenced using 16 cycles for the i7 index read and 8 cycles for the i5 index read. Reads were demultiplexed with bcl2fastq2 (v2.20.0) using options --use-bases-mask Y*,I8Y*,I*,Y* --mask-short-adapter-reads 0 --minimum-trimmed-read-length 0 (Illumina, San Diego, CA, USA), and UMIs in the index FASTQ files were attached to the read FASTQ files. Adapters were trimmed with cutadapt (v1.18),^72^ and trimmed reads shorter than 20 base pairs were removed. FastQC (v0.11.8) was used to generate pre-alignment QC metrics. STAR (v2.7.0d)^86^ was used to index and align reads to release 38 of the Ensembl Homo sapiens (hg38) genome and Gencode (Version 29). Default parameters were used for STAR’s genomeGenerate run mode; in STAR’s alignReads run mode, SAM attributes were specified as NH HI AS NM MD nM, and reads were removed if they did not contain high-confidence collapsed splice junctions (--outFilterType BySJout). RSEM (v1.3.1)^87^ was used to quantify transcriptome-coordinate-sorted alignments using a forward probability of 0.5 to indicate a non-strand-specific protocol. Bowtie 2 (v2.3.4.3)^88^ was used to index and align reads to globin, rRNA, and phix sequences in order to quantify the percent of reads that mapped to these contaminants and spike-ins. UCSC’s gtfToGenePred was used to convert the hg38 gene annotation (GTF) to a refFlat file in order to run Picard CollectRnaSeqMetrics (v2.18.16) with options MINIMUM_LENGTH=50 and RRNA_FRAGMENT_PERCENTAGE=0.3. UMIs were used to accurately quantify PCR duplicates with NuGEN’s “nodup.py” script (https://github.com/tecangenomics/nudup). QC metrics from every stage of the quantification pipeline were compiled, in part with multiQC (v1.6). The openWDL-based implementation of the RNA-Seq pipeline on Google Cloud Platform is available on Github (https://github.com/MoTrPAC/motrpac-rna-seq-pipeline). Filtering of lowly expressed genes and normalization were performed separately in each tissue. RSEM gene counts were used to remove lowly expressed genes, defined as having 0.5 or fewer counts per million in at least 10% of samples. These filtered raw counts were used as input for differential analysis with the variancePartition::dream,^76^ as described in the statistical analysis methods section. To generate normalized sample-level data for downstream visualization, filtered gene counts were TMM-normalized using edgeR::calcNormFactors, followed by conversion to log2 counts per million with edgeR::cpm.

Principal Component Analysis and calculation of the variance explained by variables of interest were used to identify and quantify potential batch effects. Based on this analysis, processing batch, Clinical Site, percentage of UMI duplication, and RNA Integrity Number (RIN) technical effects were regressed out of the TMM-normalized counts via linear regression using limma::RemoveBatchEffect function in R.^89^ A design matrix including age, sex, and a combination of group and timepoint was used during batch effect removal to avoid removing variance attributable to biological effects.

#### Proteomics and Phosphoproteomics

##### Study design

LC-MS/MS analysis of 379 muscle samples encompassing baseline and 3 post-intervention timepoints from all three groups (control, endurance and resistance) was performed at the Broad Institute of MIT and Harvard (BI) and Pacific Northwest National Laboratories (PNNL). Samples were split evenly across the two sites, and a total of 14 samples were processed and analyzed at both sites to serve as cross-site replicates for evaluation of reproducibility. Additionally, 46 adipose tissue samples representing baseline and 4-hours post-intervention from all three groups were analyzed at PNNL.

##### Generation of common reference

Tissue-specific common reference material was generated from bulk human samples. The common reference sample for muscle consisted of bulk tissue digest from 5 individuals at 2/3 ratio of female/male. Samples were split equally between BI and PNNL, digested at each site following sample processing protocol described below, then mixed all digests from both sites and centrally aliquoted at PNNL. Common reference for adipose tissue was generated at PNNL using bulk tissue from 6 individuals representing a 4:2 ratio of female:male. 250 μg aliquots of both tissue specific common reference samples were made to be included in each multiplex (described below) and additional aliquots are stored for inclusion in future MoTrPAC phases to facilitate data integration.

##### Sample processing

Proteomics analyses were performed using clinical proteomics protocols described previously.^90,91^ Muscle tissue samples were lysed in ice-cold, freshly-prepared lysis buffer (8 M urea (Sigma-Aldrich, St. Louis, Missouri), 50 mM Tris pH 8.0, 75 mM sodium chloride, 1 mM EDTA, 2 μg/ml Aprotinin (Sigma-Aldrich, St. Louis, Missouri), 10 μg/ml Leupeptin (Roche CustomBiotech, Indianapolis, Indiana), 1 mM PMSF in EtOH, 10 mM sodium fluoride, 1% phosphatase inhibitor cocktail 2 and 3 (Sigma-Aldrich, St. Louis, Missouri), 10 mM Sodium Butyrate, 2 μM SAHA, and 10 mM nicotinamide and protein concentration was determined by BCA assay. Protein lysate concentrations were normalized, and protein was reduced with 5 mM dithiothreitol (DTT, Sigma-Aldrich) for 1 hour at 37°C with shaking at 1000 rpm on a thermomixer, alkylated with iodoacetamide (IAA, Sigma-Aldrich) in the dark for 45 minutes at 25°C with shaking at 1000 rpm, followed by dilution of 1:4 with Tris-HCl, pH 8.0 prior to adding digestion enzymes. Proteins were first digested with LysC endopeptidase (Wako Chemicals) at a 1:50 enzyme:substrate ratio (2 hours, 25 °C, 850 rpm), followed by digestion with trypsin (Promega) at a 1:50 enzyme:substrate ratio (or 1:10 ratio for adipose tissue; 14 hours, 25 °C, 850 rpm). The next day formic acid was added to a final concentration of 1% to quench the reaction. Digested peptides were desalted using Sep-Pac C18 columns (Waters), concentrated in a vacuum centrifuge, and a BCA assay was used to determine final peptide concentrations. 250μg aliquots of each sample were prepared, dried down by vacuum centrifugation and stored at -80°C.

Tandem mass tag (TMT) 16-plex isobaric labeling reagent (ThermoFisher Scientific) was used for this study. Samples were randomized across the first 15 channels of TMT 16-plexes, and the last channel (134N) of each multiplex was used for a common reference that was prepared prior to starting the study (see above). Randomization of samples across the plexes within each site was done using https://github.com/MoTrPAC/clinical-sample-batching, with the goal to have all timepoints per participant in the same plex, and uniform distribution of groups (endurance, resistance, control), sex and sample collection clinical site across the plexes.

Peptide aliquots (250 μg per sample) were resuspended to a final concentration of 5 μg/μL in 200 mM HEPES, pH 8.5 for isobaric labeling. TMT reagent was added to each sample at a 1:2 peptide: TMT ratio, and labeling proceeded for 1 hour at 25°C with shaking at 400 rpm. The labeling reaction was diluted to a peptide concentration of 2 µg/µL using 62.5 μL of 200 mM HEPES and 20% ACN. 3 μL was removed from each sample to quantify labeling efficiency and mixing ratio. After labeling QC analysis, reactions were quenched with 5% hydroxylamine and samples within each multiplex were combined and desalted with Sep-Pac C18 columns (Waters).

Combined TMT multiplexed samples were then fractionated using high pH reversed phase chromatography on a 4.6mm ID x 250mm length Zorbax 300 Extend-C18 column (Agilent) with 5% ammonium formate/2% Acetonitrile as solvent A and 5% ammonium formate/90% acetonitrile as solvent B. Samples were fractionated with 96min separation gradient at flow rate of 1mL/min and fractions were collected at each minute onto a 96-well plate. Fractions are then concatenated into 24 fractions with the following scheme: fraction 1 = A1+C1+E1+G1, fraction 2 = A2+C2+E2+G2, fraction 3 = A3+C3+E3+G3, all the way to fraction 24 = B12+D12+F12+H12 following the same scheme. 5% of each fraction was removed for global proteome analysis, and the remaining 95% was further concatenated to 12 fractions for phosphopeptide enrichment using immobilized metal affinity chromatography (IMAC).

Phosphopeptide enrichment was performed through immobilized metal affinity chromatography (IMAC) using Fe 3+ -NTA-agarose beads, freshly prepared from Ni-NTA-agarose beads (Qiagen, Hilden, Germany) by sequential incubation in 100 mM EDTA to strip nickel, washing with HPLC water, and incubation in 10 mM iron (III) chloride). Peptide fractions were resuspended to 0.5 μg/uL in 80% ACN + 0.1% TFA and incubated with beads for 30 minutes in a thermomixer set to 1000 rpm at room temperature. After 30 minutes, beads were spun down (1 minute, 1000 rcf) and supernatant was removed and saved as flow-through for subsequent enrichments. Phosphopeptides were eluted off IMAC beads in 3x 75 μL of agarose bead elution buffer (500 mM K2HPO4, pH 7.0), desalted using C18 stage tips, eluted with 50% ACN, and lyophilized. Samples were then reconstituted in 3% ACN / 0.1% FA for LC-MS/MS analysis (9 μL reconstitution / 4 μL injection at the BI; 12 μL reconstitution / 5 μL injection at PNNL).

##### Data acquisition

Broad Institute: Both proteome and phosphoproteome samples were analyzed on 75um ID Picofrit columns packed in-house with ReproSil-Pur 120 Å, C18-AQ, 1.9 µm beads to the length of 20-24cm. Online separation was performed on Easy nLC 1200 systems (ThermoFisher Scientific) with solvent A of 0.1% formic acid/3% acetonitrile and solvent B of 0.1% formic acid/90% acetonitrile, flow rate of 200nL/min and the following gradient: 2-6% B in 1min, 6-20% B in 52min, 20-35% B in 32min, 35-60% B in 9min, 60-90% B in 1min, followed by a 5min hold at 90%, and 9min hold at 50%. Proteome fractions were analyzed on a Q Exactive Plus mass spectrometer (ThrmoFisher Scientific) with MS1 scan across the 300-1800 m/z range at 70,000 resolution, AGC target of 3x10^6^ and maximum injection time of 5ms. MS2 scans of most abundant 12 ions were performed at 35,000 resolution with AGC target of 1x10^5^ and maximum injection time of 120ms, isolation window of 0.7m/z and normalized collision energy of 27. Phosphoproteome fractions were analyzed on Q Exactive HFX MS system (ThermoFisher Scientific) with the following parameters: MS1 scan across 350-1800 m/z mass range at 60,000 resolution, AGC target of 3x10^6^ and maximum injection time of 10ms. 20 most abundant ions are fragmented in MS2 scans with AGC of 1x10^5^, maximum injection time of 105ms, isolation width of 0.7 m/z and NCE of 29. In both methods dynamic exclusion was set to 45sec.

PNNL: For mass spectrometry analysis of the global proteome of muscle samples, online separation was performed using a nanoAcquity M-Class UHPLC system (Waters) and a 25 cm x 75 μm i.d. picofrit column packed in-house with C18 silica (1.7 μm UPLC BEH particles, Waters Acquity) with solvent A of 0.1% formic acid/3% acetonitrile and solvent B of 0.1% formic acid/90% acetonitrile, flow rate of 200nL/min and the following gradient: 1% B for 8min, 8-20% B in 90min, 20-35% B in 13min, 35-75% B in 5min, 75-95% B in 3min, followed by a 6min hold at 90%, and 9min hold at 50%. Proteome fractions were analyzed on a Q Exactive Plus mass spectrometer (ThermoFisher Scientific) with MS1 scan across the 300-1800 m/z range at 60,000 resolution, AGC target of 3x10^6^ and maximum injection time of 20ms. MS2 scans of most abundant 12 ions were performed at 30,000 resolution with AGC target of 1x10^5^ and maximum injection time of 100ms, isolation window of 0.7m/z and normalized collision energy of 30.

For mass spectrometry analysis of the phosphoproteome, online separation was performed using a Dionex Ultimate 3000 UHPLC system (ThermoFisher) and a 25 cm x 75 μm i.d. picofrit column packed in-house with C18 silica (1.7 μm UPLC BEH particles, Waters Acquity) with solvent A of 0.1% formic acid/3% acetonitrile and solvent B of 0.1% formic acid/90% acetonitrile, flow rate of 200nL/min and the following gradient: 1-8% B in 10min, 8-25% B in 90min, 25-35% B in 10min, 35-75% B in 5min, and 75-5% in 3min. Phosphoproteome fractions were analyzed on a Q Exactive HF-X Plus mass spectrometer (ThermoFisher Scientific) with MS1 scan across the 300-1800 m/z range at 60,000 resolution, AGC target of 3x10^6^ and maximum injection time of 20ms. MS2 scans of most abundant 12 ions were performed at 45,000 resolution with AGC target of 1x10^5^ and maximum injection time of 100ms, isolation window of 0.7m/z and normalized collision energy of 30.

##### Data searching

Proteome and phosphoproteome data from both BI and PNNL were searched at the Bioinformatics Center (BIC) using the MSGF+ cloud-based pipeline previously described ^92^ against a composite protein database. This database comprised UniProt canonical sequences (downloaded 2022-09-13; 20383 sequences), UniProt human protein isoforms (downloaded 2022-09-13; 21982 sequences), and common contaminants (261 sequences), resulting in 42,626 sequences.

##### QC/Filtering/Normalization

The log2 Reporter ion intensity (RII) ratios to the common reference were used as quantitative values for all proteomics features (proteins and phosphosites). Datasets were filtered to remove features identified from contaminant proteins and decoy sequences. Datasets were visually evaluated for sample outliers by looking at top principal components, examining median feature abundance and distributions of RII ratio values across samples, and by quantifying the number of feature identifications within each sample. No outliers were detected in the muscle tissue dataset. The Log_2_ RII ratio values were normalized within each sample by median centering to zero. Principal Component Analysis and calculation of the variance explained by variables of interest were used to identify and quantify potential batch effects. Based on this analysis, TMT plex, Clinical Site, and Chemical Analysis Site (muscle only, where samples were analyzed at both BI and PNNL) batch effects were removed using Linear Models Implemented in the *limma::RemoveBatchEffect()* function in R.^89^ A design matrix including age, sex, and group_timepoint was used during batch effect removal in order to preserve the effect of all variables included in later statistical analysis. Correlations between technical replicates analyzed within and across CAS (where applicable) were calculated to evaluate intra- and inter-site reproducibility; the data from technical replicates were then averaged for downstream analysis. Finally, features with quantification in less of 30% of all samples were removed. For specific details of the process, see available code.

#### Metabolomics and Lipidomics

The metabolomic analysis was performed by investigators across multiple *Chemical Analysis Sites* that employed different technical platforms for data acquisition. At the highest level, these platforms were divided into two classes: *Targeted* and *Untargeted*. The data generated by the untargeted platforms were further divided into Named (confidently identified chemical entities) and Unnamed (confidently detected, but no chemical name annotation) compounds.

Targeted metabolomics data were generated at 3 analysis Sites: Duke University, the Mayo Clinic, and Emory University. Duke quantified metabolites belonging to the metabolite classes Acetyl-CoA, Keto Acids, and Nucleic Acids ("acoa", "ka", "nuc"), Mayo quantified amines and TCA intermediates (“amines” and “tca”), while Emory quantified Oxylipins (“oxylipneg”).

The untargeted metabolomics data were generated at 3 analysis Sites: the Broad Institute, the University of Michigan, and the Georgia Institute of Technology (Georgia Tech). The Broad Institute applied Hydrophilic interaction liquid chromatography (HILIC) in the positive ion mode (hilicpos), Michigan applied reverse phase liquid chromatography in both positive and negative ion modes (“rppos” and “rpneg”) and ion-pairing chromatography in the negative mode (“ionpneg”), and Georgia Tech performed lipidomics assays using reverse phase chromatography in both positive and negative ion modes (“lrppos” and “lrpneg”).

##### LC-MS/MS analysis of branched-chain keto acids

Duke University conducted targeted profiling of branched-chain keto acid metabolites. Tissue homogenates were prepared at 100 mg/ml in 3M perchloric acid containing isotopically labeled ketoleucine (KIC)-d3, ketoisovalerate (KIV)-^13^C_5_ (from Cambridge Isotope Laboratories), and 3-methyl-2-oxovalerate (KMV)-d8 (from Toronto Research Chemicals, Canada) internal standards and 200 μL was centrifuged.

Next, 200 μL of 25 M o-phenylenediamine (OPD) in 3M HCl were added to tissue supernatants. The samples were incubated at 80°C for 20 minutes. Keto acids were then extracted using ethyl acetate, following a previously described protocol.^93,94^ The extracts were dried under nitrogen, reconstituted in 200 mM ammonium acetate, and subjected to analysis on a Xevo TQ-S triple quadrupole mass spectrometer coupled to an Acquity UPLC (Waters), controlled by the MassLynx 4.1 operating system.

The analytical column used was a Waters Acquity UPLC BEH C18 Column (1.7 μm, 2.1 × 50 mm), maintained at 30°C. 10 μL of the sample were injected onto the column and eluted at a flow rate of 0.4 ml/min. The gradient consisted of 45% mobile phase A (5 mM ammonium acetate in water) and 55% mobile phase B (methanol) for 2 minutes. This was succeeded by a linear gradient to 95% B from 2 to 2.5 minutes, holding at 95% B for 0.7 minutes, returning to 45% A, and finally re-equilibrating the column at initial conditions for 1 minute. The total run time was 4.7 minutes.

In positive ion mode, mass transitions of m/z 203 → 161 (KIC), 206 → 161 (KIC-d3), 189 → 174 (KIV), 194 → 178 (KIV-^5^C_13_), 203 → 174 (KMV), and 211 → 177 (KMV-d8) were monitored. Endogenous keto acids were quantified using calibrators prepared by spiking dialyzed fetal bovine serum with authentic keto acids (Sigma-Aldrich).

##### Flow injection MS/MS analysis of acyl CoAs

Duke University conducted targeted profiling of acyl CoAs. Here, 500 μL of tissue homogenate, prepared at a concentration of 50 mg/ml in isopropanol/0.1 M KH2PO4 (1:1), underwent extraction with an equal volume of acetonitrile. The resulting mixture was then centrifuged at 14,000 x g for 10 minutes, following a previously established procedure.^95,96^

The supernatants were acidified with 0.25 ml of glacial acetic acid, and the acyl CoAs were subjected to additional purification through solid-phase extraction (SPE) using 2-(2-pyridyl) ethyl functionalized silica gel (Sigma-Aldrich), as outlined in Minkler et al. (2008). Prior to use, the SPE columns were conditioned with 1 ml of acetonitrile/isopropanol/water/glacial acetic acid (9/3/4/4 : v/v/v/v). After application and flow-through of the supernatant, the SPE columns underwent a washing step with 2 ml of acetonitrile/isopropanol/water/glacial acetic acid (9/3/4/4 : v/v/v/v). Acyl CoAs elution was achieved with 2 ml of methanol/250 mM ammonium formate (4/1 : v/v), followed by analysis using flow injection MS/MS in positive ion mode on a Xevo TQ-S triple quadrupole mass spectrometer (Waters). The mobile phase used was methanol/water (80/20, v/v) with 30 mM ammonium hydroxide. Spectra were acquired in the multichannel acquisition mode, monitoring the neutral loss of 507 amu (phosphoadenosine diphosphate) and scanning from m/z 750 to 1100. As an internal standard, heptadecanoyl CoA was employed.

Quantification of endogenous acyl CoAs was carried out using calibrators created by spiking tissue homogenates with authentic Acyl CoAs (Sigma-Aldrich) that covered saturated acyl chain lengths from C0 to C18. Empirical corrections for heavy isotope effects, particularly ^13^C, to the adjacent m+2 spectral peaks in a specific chain length cluster were made by referring to the observed spectra for the analytical standards.

##### LC-MS/MS analysis of nucleotides

Duke University conducted targeted profiling of nucleotide metabolites. 300 μL of tissue homogenates, prepared at a concentration of 50 mg/ml in 70% methanol, underwent spiking with nine internal standards: ^13^C^10^,^15^N^5^-adenosine monophosphate, ^13^C^10^,^15^N^5^-guanosine monophosphate, ^13^C^10^,^15^N^2^-uridine monophosphate, ^13^C^9^,^15^N^3^-cytidine monophosphate, ^13^C^10^-guanosine triphosphate, ^13^C^10^-uridine triphosphate, ^13^C^9^-cytidine triphosphate, ^13^C^10^-adenosine triphosphate, and nicotinamide-1, N^6^-ethenoadenine dinucleotide (eNAD) (all from Sigma-Aldrich).

Nucleotides were extracted using an equal volume of hexane, following the procedures outlined by Cordell et al. and Gooding et al..^97,98^ After vortexing and centrifugation at 14,000 x g for 5 minutes, the bottom layer was subjected to another centrifugation. Chromatographic separation and MS analysis of the supernatants were performed using an Acquity UPLC system (Waters) coupled to a Xevo TQ-XS quadrupole mass spectrometer (Waters).

The analytical column employed was a Chromolith FastGradient RP-18e 50-2mm column (EMD Millipore, Billerica, MA, USA), that was maintained at 40°C. The injection volume was 2 μL. Nucleotides were separated using a mobile phase A consisting of 95% water, 5% methanol, and 5 mM dimethylhexylamine adjusted to pH 7.5 with acetic acid. Mobile phase B comprised 20% water, 80% methanol, and 10 mM dimethylhexylamine. The flow rate was set to 0.3 ml/min. The 22-minute gradient (t=0, %B=0; t=1.2, %B=0; t=22, %B=40) was followed by a 3-minute wash and 7-minute equilibration. Nucleotides were detected in the negative ion multiple reaction monitoring (MRM) mode based on characteristic fragmentation reactions. Endogenous nucleotides were quantified using calibrators prepared by spiked-in authentic nucleotides obtained from Sigma-Aldrich.

##### LC-MS/MS analysis of amino metabolites

The Mayo Clinic conducted LC-MS-based targeted profiling of amino acids and amino metabolites, as previously outlined.^99,100^ In summary, 5 mg of tissue homogenates were supplemented with an internal standard solution comprising isotopically labeled amino acids (U-^13^C^4^ L-aspartic acid, U-^13^C_3_ alanine, U-^13^C_4_ L-threonine, U-^13^C L-proline, U-^13^C_6_ tyrosine, U-^13^C_5_ valine, U-^13^C_6_ leucine, U-^13^C_6_ phenylalanine, U-^13^C_3_ serine, U-^13^C_5_ glutamine, U-^13^C_2_ glycine, U-^13^C_5_ glutamate, U-^13^C_6_,^15^N_2_ lysine, U-^13^C_5_,^15^N methionine, 1,1U-^13^C_2_ homocysteine, U-^13^C_6_ arginine, U-^13^C_5_ ornithine, ^13^C_4_ asparagine, ^13^C_2_ ethanolamine, D_3_ sarcosine, D_6_ 4-aminobutyric acid).

The supernatant was promptly derivatized using 6-aminoquinolyl-N-hydroxysuccinimidyl carbamate with a MassTrak kit (Waters). A 10-point calibration standard curve underwent a similar derivatization procedure after the addition of internal standards. Both the derivatized standards and samples were subjected to analysis using a Quantum Ultra triple quadrupole mass spectrometer (ThermoFischer) coupled with an Acquity liquid chromatography system (Waters). Data acquisition utilized selected ion monitoring (SRM) in positive ion mode. The concentrations of 42 analytes in each unknown sample were calculated against their respective calibration curves.

##### GC-MS analysis of TCA metabolites

The Mayo Clinic conducted GC-MS-targeted profiling of tricarboxylic acid (TCA) metabolites, as previously detailed,^101,102^ with some modifications. In summary, 5 mg of tissue were homogenized in 1X PBS using an Omni bead ruptor (Omni International, Kennesaw, GA), followed by the addition of 20 μL of an internal solution containing U-^13^C labeled analytes (^13^C_3_ sodium lactate, ^13^C_4_ succinic acid, ^13^C_4_ fumaric acid, ^13^C_4_ alpha-ketoglutaric acid, ^13^C_4_ malic acid, ^13^C_4_ aspartic acid, ^13^C_5_ 2-hydroxyglutaric acid, ^13^C_5_ glutamic acid, ^13^C_6_ citric acid, ^13^C_2_,^15^N glycine, ^13^C_2_ sodium pyruvate). For plasma, 50 μL were used.

The proteins were precipitated out using 300 μL of a mixture of chilled methanol and acetonitrile solution. After drying the supernatant in the speedvac, the sample was derivatized first with ethoxime and then with MtBSTFA + 1% tBDMCS (N-Methyl-N-(t-Butyldimethylsilyl)-Trifluoroacetamide + 1% t-Butyldimethylchlorosilane) and then analyzed on an Agilent 5977B GC/MS (Santa Clara, California) under single ion monitoring conditions using electron ionization. Concentrations of lactic acid (m/z 261.2), fumaric acid (m/z 287.1), succinic acid (m/z 289.1), ketoglutaric acid (m/z 360.2), malic acid (m/z 419.3), aspartic acid (m/z 418.2), 2-hydroxyglutaratic acid (m/z 433.2), cis-aconitic acid (m/z 459.3), citric acid (m/z 591.4), isocitric acid (m/z 591.4), and glutamic acid (m/z 432.4) were measured against 7-point calibration curves that underwent the same derivatization procedure.

##### Targeted lipidomics of low-level lipids

Lipid targeted profiling was conducted at Emory University following established methodologies as previously described.^103,104^ In summary, 20 mg of powdered tissue samples were homogenized in 100 μL PBS using Bead Ruptor (Omni International, Kennesaw, GA). Homogenized tissue samples were diluted with 300 μL 20% methanol and spiked with a 1% BHT solution to a final BHT concentration of 0.1% and pH of 3.0 by acetic acid addition. After centrifugation (10 minutes, 14000 rpm), the supernatants were transferred to 96-well plates for further extraction.

The supernatants were loaded onto Isolute C18 solid-phase extraction (SPE) columns that had been conditioned with 1000 μL ethyl acetate and 1000 μL 5% methanol. The SPE columns were washed with 800 μL water and 800 μL hexane, and the oxylipins were then eluted with 400 μL methyl formate. The SPE process was automated using a Biotage Extrahera (Uppsala, Sweden). The eluate was dried with nitrogen and reconstituted with 200 μL acetonitrile before LC-MS analysis. Sample blanks, pooled extract samples used as quality controls (QC), and consortium reference samples were prepared for analysis using the same methods. All external standards were purchased from Cayman Chemical (Ann Arbor, Michigan) at a final concentration in the range 0.01-20 μg/ml and consisted of: prostaglandin E2 ethanolamide (Catalog No. 100007212), oleoyl ethanolamide (Catalog No. 90265), palmitoyl ethanolamide (Catalog No. 10965), arachidonoyl ethanolamide (Catalog No. 1007270), docosahexaenoyl ethanolamide (Catalog No. 10007534), linoleoyl ethanolamide (Catalog No. 90155), stearoyl ethanolamide (Catalog No. 90245), oxy-arachidonoyl ethanolamide (Catalog No. 10008642), 2-arachidonyl glycerol (Catalog No. 62160), docosatetraenoyl ethanolamide (Catalog No. 90215), α-linolenoyl ethanolamide (Catalog No. 902150), oleamide (Catalog No. 90375), dihomo-γ-linolenoyl ethanolamide (Catalog No. 09235), decosanoyl ethanolamide (Catalog No. 10005823), 9,10 DiHOME (Catalog No. 53400), prostaglandin E2-1-glyceryl ester (Catalog No. 14010), 20-HETE (Catalog No. 10007269), 9-HETE (Catalog No. 34400), 14,15 DiHET (Catalog No. 10007267), 5(S)-HETE (Catalog No. 34210), 12(R)-HETE (Catalog No. 10007247), 11(12)-DiHET (Catalog No. 10007266), 5,6-DiHET (Catalog No. 10007264), thromboxane B2 (Catalog No. 10007237), 12(13)-EpOME (Catalog No. 52450), 13 HODE (Catalog No. 38600), prostaglandin F2α (Catalog No. 10007221), 14(15)-EET (Catalog No. 10007263), 8(9)-EET (Catalog No. 10007261), 11(12)-EET (Catalog No. 10007262), leukotriene B4 (Catalog No. 20110), 8(9)-DiHET (Catalog No. 10007265), 13-OxoODE (Catalog No. 38620), 13(S)-HpODE (Catalog No. 48610), 9(S)-HpODE (Catalog No. 48410), 9(S)-HODE (Catalog No. 38410), resolvin D3 (Catalog No. 13834), resolvin E1 (Catalog No. 10007848), resolvin D1 (Catalog No. 10012554), resolvin D2 (Catalog No. 10007279), 9(S)HOTrE (Catalog No. 39420), 13(S)HOTrE (Catalog No. 39620), 8-iso Progstaglandin F2α (Catalog No. 25903), maresin 1 (Catalog No. 10878), maresin 2 (Catalog No. 16369).

LC-MS data were acquired using an Agilent 1290 Infinity II chromatograph from Agilent (Santa Clara, CA), equipped with a ThermoFisher Scientific AccucoreTM C18 column (100 mm × 4.6, 2.6 µm particle size), and coupled to Agilent 6495 mass spectrometer for polarity switch multiple reaction monitoring (MRM) scan. The mobile phases consisted of water with 10 mM ammonium acetate (mobile phase A) and acetonitrile with 10 mM ammonium acetate (mobile phase B). The chromatographic gradient program was: 0.5 minutes with 95% A; 1 minute to 2 minutes with 65% A; 2.1 minutes to 5.0 minutes with 45% A; 7 minutes to 20 minutes with 25% A; and 21.1 minutes until 25 minutes with 95% A. The flow rate was set at 0.40 ml/min. The column temperature was maintained at 35°C, and the injection volume was 6 μL. For MS analysis, the MRM analysis is operated at gas temperature of 290 °C, gas flow of 14 L/min, Nebulizer of 20 psi, sheath gas temperature of 300 °C, sheath gas flow of 11 L /min, capillary of 3000 V for both positive and negative ion mode, nozzle voltage of 1500 V for both positive and negative ion mode, high pressure RF of iFunnel parameters of 150V for both positive and negative ion mode, and low pressure of RF of iFunnel parameters of 60 V for both positive and negative ion mode. Skyline (version 25.1.0.142)^105^ was utilized to process raw LC-MS data. Standard curves were constructed for each oxylipin/ethanolamide and scrutinized to ensure that all concentration points fell within the linear portion of the curve with an R-squared value not less than 0.9. Additionally, features exhibiting a high coefficient of variation (CV) among the quality control (QC) samples were eliminated from the dataset. Pearson correlation among the QCs for each tissue type was computed using the Hmisc R library, and the figures documented in the QC report were generated and visualized with the corrplot R library.^106,107^

##### Hydrophilic interaction LC-MS metabolomics

The untargeted analysis of polar metabolites in the positive ionization mode was conducted at the Broad Institute of MIT and Harvard. The LC-MS system consisted of a Shimadzu Nexera X2 UHPLC (Shimadzu Corp., Kyoto, Japan) coupled to a Q-Exactive hybrid quadrupole Orbitrap mass spectrometer (Thermo Fisher Scientific). Tissue (10 mg) homogenization was performed at 4°C using a TissueLyser II (QAIGEN) bead mill set to two 2 min intervals at 20 Hz in 300 μL of 10/67.4/22.4/0.018 v/v/v/v water/acetonitrile/methanol/formic acid valine-d8 and phenylalanine-d8 internal standards. The column was eluted isocratically at a flow rate of 250 μL/min with 5% mobile phase A (10 mM ammonium formate and 0.1% formic acid in water) for 0.5 minute, followed by a linear gradient to 40% mobile phase B (acetonitrile with 0.1% formic acid) over 10 minutes, then held at 40% B for 4.5 minutes. MS analyses utilized electrospray ionization in the positive ion mode, employing full scan analysis over 70-800 m/z at 70,000 resolution and 3 Hz data acquisition rate. Various MS settings, including sheath gas, auxiliary gas, spray voltage, and others, were specified for optimal performance. Data quality assurance was performed by confirming LC-MS system performance with a mixture of >140 well-characterized synthetic reference compounds, daily evaluation of internal standard signals, and the analysis of four pairs of pooled extract samples per sample type inserted in the analysis queue at regular intervals. One sample from each pair was used to correct for instrument drift using “nearest neighbor” scaling while the second reference sample served as a passive QC for determination of the analytical coefficient of variation of every identified metabolite and unknown. Raw data processing involved the use of TraceFinder software (v3.3, Thermo Fisher Scientific) for targeted peak integration and manual review, as well as Progenesis QI (v3.0, Nonlinear Dynamics, Waters) for peak detection and integration of both identified and unknown metabolites. Metabolite identities were confirmed using authentic reference standards.

##### Reversed Phase-High Performance & Ion-pairing LC metabolomics

Reversed-phase and ion pairing LC-MS profiling of polar metabolites was conducted at the University of Michigan. LC-MS grade solvents and mobile phase additives were procured from Sigma-Aldrich, while chemical standards were obtained from either Sigma-Aldrich or Cambridge Isotope Labs. Frozen muscle tissue samples were rapidly weighed into pre-tared, pre-chilled Eppendorf tubes and extracted in 1:1:1:1 v:v methanol:acetonitrile:acetone:water containing the following internal standards diluted from stock solutions to yield the specified concentrations: ^13^C_3_-lactic acid, 12.5 µM; ^13^C_5_-oxoglutaric acid, 125 nM;^13^C_5_-citric acid,1.25 µM; ^13^C_4_-succinic acid,125 nM; ^13^C_4_-malic acid, 125 nM; U-^13^C amino acid mix (Cambridge Isotope CLM-1548-1), 2.5 µg/mL; ^13^C_5_-glutamine, 6.25 µM;^15^N_2_-asparagine, 1.25 µM;^15^N_2_-tryptophan, 1.25 µM; ^13^C_6_-glucose, 62.5 µM; D_4_-thymine, 1 µM; ^15^N-anthranillic acid, 1 µM; gibberellic acid,1 µM; epibrassinolide, 1 µM. The extraction solvent also contained a 1:400 dilution of Cambridge Isotope carnitine/acylcarnitine mix NSK-B, resulting in the following concentrations of carnitine/acylcarnitine internal standards: D_9_-L-carnitine, 380 nM; D_3_-L-acetylcarnitine, 95 nM; D_3_-L-propionylcarnitine, 19 nM; D_3_-L-butyrylcarnitine, 19 nM; D_9_-L-isovalerylcarnitine, 19 nM; D_3_-L-octanoylcarnitine, 19 nM; D_9_-L-myristoylcarnitine, 19 nM; D_3_-L-palmitoylcarnitine, 38 nM. Sample extraction was performed by adding chilled extraction solvent to tissue sample at a ratio of 1 ml solvent to 50 mg wet tissue mass. Immediately after solvent addition, the sample was homogenized using a Branson 450 probe sonicator. Subsequently, the tubes were mixed several times by inversion and then incubated on ice for 10 minutes. Following incubation, the samples were centrifuged and 300 μL of the supernatant was carefully transferred to two autosampler vials with flat-bottom inserts and dried under a constant stream of nitrogen gas. A QC sample was created by pooling residual supernatants from multiple samples. This QC sample underwent the same drying and reconstitution process as described for individual samples. Dried supernatants were stored at -80 C until ready for instrumental analysis. On the day of analysis, samples were reconstituted in 60 μL of 8:2 v:v water:methanol and then submitted to LC-MS.

Reversed phase LC-MS samples were analyzed on an Agilent 1290 Infinity II / 6545 qTOF MS system with a JetStream electrospray ionization (ESI) source (Agilent Technologies, Santa Clara, California) using a Waters Acquity HSS T3 column, 1.8 µm 2.1 x 100 mm equipped with a matched Vanguard precolumn (Waters Corporation). Mobile phase A was 100% water with 0.1% formic acid and mobile phase B was 100% methanol with 0.025% formic acid. The gradient was as follows: Linear ramp from 0% to 100% B from 0-10 minutes, hold 100% B until 17 minutes, linear return to 0% B from 17 to 17.1 minutes, hold 0% B until 20 minutes. The flow rate was 0.45 ml/min, the column temperature was 55°C, and the injection volume was 5 μL. Each sample was analyzed twice, once in positive and once in negative ion mode MS, scan rate 2 spectra/sec, mass range 50-1200 m/z. Source parameters were: drying gas temperature 350°C, drying gas flow rate 10 L/min, nebulizer pressure 30 psig, sheath gas temperature 350°C and flow 11 L/minute, capillary voltage 3500 V, internal reference mass correction enabled. A QC sample run was performed at minimum every tenth injection.

Ion-pair LC-MS samples were analyzed on an identically-configured LC-MS system using an Agilent Zorbax Extend C18 1.8 µm RRHD column, 2.1 x 150 mm ID, equipped with a matched guard column. Mobile phase A was 97% water, 3% methanol. Mobile phase B was 100% methanol. Both mobile phases contained 15 mM tributylamine and 10 mM acetic acid. Mobile phase C was 100% acetonitrile. Elution was carried out using a linear gradient followed by a multi-step column wash including automated (valve-controlled) backflushing, detailed as follows: 0-2min, 0%B; 2-11 min, linear ramp from 0-99%B; 12-16 min, 99%B, 16-17.5min, 99-0%B. At 17.55 minutes, the 10-port valve was switched to reverse flow (back-flush) through the column. From 17.55-20.45 min the solvent was ramped from 99%B to 99%C. From 20.45-20.95 min the flow rate was ramped up to 0.8 mL/min, which was held until 22.45 min, then ramped down to 0.6mL/min by 22.65 min. From 22.65-23.45 min the solvent was ramped from 99% to 0% C while flow was simultaneously ramped down from 0.6-0.4mL/min. From 23.45 to 29.35 min the flow was ramped from 0.4 to 0.25mL/min; the 10-port valve was returned to restore forward flow through the column at 25 min. Column temperature was 35°C and the injection volume was 5 μL. MS acquisition was performed in negative ion mode, scan rate 2 spectra/sec, mass range 50-1200 m/z. Source parameters were: drying gas temperature 250°C, drying gas flow rate 13 L/min, nebulizer pressure 35 psig, sheath gas temp 325°C and flow 12 L/min, capillary voltage 3500V, internal reference mass correction enabled. A QC sample run was performed at minimum every tenth injection.

Iterative MS/MS data was acquired for both reverse phase and ion pairing methods using the pooled sample material to enable compound identification. Eight repeated LC-MS/MS runs of the QC sample were performed at three different collision energies (10, 20, and 40) with iterative MS/MS acquisition enabled. The software excluded precursor ions from MS/MS acquisition within 0.5 minute of their MS/MS acquisition time in prior runs, resulting in deeper MS/MS coverage of lower-abundance precursor ions.

Feature detection and alignment was performed utilizing a hybrid targeted/untargeted approach. Targeted compound detection and relative quantitation was performed by automatic integration followed by manual inspection and correction using Profinder v8.0 (Agilent Technologies, Santa Clara, CA.) Non-targeted feature detection was performed using custom scripts that automate operation of the “find by molecular feature” workflow of the Agilent Masshunter Qualitative Analysis (v7) software package. Feature alignment and recursive feature detection were performed using Agilent Mass Profiler Pro (v8.0) and Masshunter Qualitative Analysis (“find by formula” workflow), yielding an aligned table including m/z, RT, and peak areas for all features.

###### Data Cleaning and Degeneracy Removal

The untargeted features and named metabolites were merged to generate a combined feature table. Features missing from over 50% of all samples in a batch or over 30% of QC samples were then removed prior to downstream normalization procedure. Next, the software package Binner was utilized to remove redundancy and degeneracy in the data.^108^ Briefly, Binner first performs RT-based binning, followed by clustering of features by Pearson’s correlation coefficient, and then assigns annotations for isotopes, adducts or in-source fragments by searching for known mass differences between highly correlated features.

###### Normalization and Quality Control

Data were then normalized using a Systematic Error Removal Using Random Forest (SERRF) approach,^109^ which helps correct for drift in peak intensity over the batch using data from the QC sample runs. When necessary to correct for residual drift, peak area normalization to closest-matching internal standard was also applied to selected compounds. Both SERRF correction and internal standard normalization were implemented in R. Parameters were set to minimize batch effects and other observable drifts, as visualized using principal component analysis score plots of the full dataset. Normalization performance was also validated by examining relative standard deviation values for additional QC samples not included in the drift correction calculations.

###### Compound identification

Metabolites from the targeted analysis workflow were identified with high confidence (MSI level 1)^110^ by matching retention time (+/- 0.1 minute), mass (+/- 10 ppm) and isotope profile (peak height and spacing) to authentic standards. MS/MS data corresponding to unidentified features of interest from the untargeted analysis were searched against a spectral library (NIST 2020 MS/MS spectral database or other public spectral databases) to generate putative identifications (MSI level 2) or compound-class level annotations (MSI level 3) as described previously.^111^

##### LC-MS/MS untargeted lipidomics

###### Sample preparation

Non-targeted lipid analysis was conducted at the Georgia Institute of Technology. Powdered tissue samples (10 mg) were extracted in 400 μL isopropanol containing stable isotope-labeled internal standards (IS) with bead homogenization with 2mm zirconium oxide beads (Next Advance) using a TissueLyser II (10min, 30 Hz). Samples were then centrifuged (5 min, 21,100xg), and supernatants were transferred to autosampler vials. Sample blanks, pooled extract samples used as quality controls (QC), and consortium reference samples, were prepared for analysis using the same methods. The IS mix consisted of PC (15:0-18:1(d7)), Catalog No. 791637; PE (15:0-18:1(d7)), Catalog No. 791638; PS (15:0-18:1(d7)), Catalog No. 791639; PG(15:0-18:1(d7)), Catalog No. 791640; PI(15:0-18:1(d7)), Catalog No. 791641; LPC(18:1(d7)), Catalog No. 791643; LPE(18:1(d7)); Catalog No. 791644; Chol Ester (18:1(d7)), Catalog No. 700185; DG(15:0-18:1(d7)), Catalog No. 791647; TG(15:0-18:1(d7)-15:0), Catalog No. 791648; SM(18:1(d9)), Catalog No. 791649; Cholesterol (d7), Catalog No. 700041. All internal standards were purchased from Avanti Polar Lipids (Alabaster, Alabama) and added to the extraction solvent at a final concentration in the 0.1-8 μg/ml range.

###### Data collection

Lipid LC-MS data were acquired using a Vanquish (ThermoFisher Scientific) chromatograph fitted with a ThermoFisher Scientific AccucoreTM C30 column (2.1 × 150 mm, 2.6 µm particle size), coupled to a high-resolution accurate mass Q-Exactive HF Orbitrap mass spectrometer (ThermoFisher Scientific) for both positive and negative ionization modes. For positive mode analysis, the mobile phases were 40:60 water:acetonitrile with 10 mM ammonium formate and 0.1% formic acid (mobile phase A), and 10:90 acetonitrile:isopropyl alcohol, with 10 mM ammonium formate and 0.1% formic acid (mobile phase B). For negative mode analysis, the mobile phases were 40:60 water:acetonitrile with 10 mM ammonium acetate (mobile phase A), and 10:90 acetonitrile:isopropyl alcohol, with 10 mM ammonium acetate (mobile phase B). The chromatographic method used for both ionization modes was the following gradient program: 0 minutes 80% A; 1 minute 40% A; 5 minutes 30% A; 5.5 minutes 15% A; 8 minutes 10% A; held 8.2 minutes to 10.5 minutes 0% A; 10.7 minutes 80% A; and held until 12 minutes. The flow rate was set at 0.40 ml/min. The column temperature was set to 50°C, and the injection volume was 2 μL.

For analysis of the organic phase the electrospray ionization source was operated at a vaporizer temperature of 425°C, a spray voltage of 3.0 kV for positive ionization mode and 2.8 kV for negative ionization mode, sheath, auxiliary, and sweep gas flows of 60, 18, and 4 (arbitrary units), respectively, and capillary temperature of 275°C. The instrument acquired full MS data with 240,000 resolution over the 150-2000 m/z range. LC-MS/MS experiments were acquired using a DDA strategy. MS2 spectra were collected with a resolution of 120,000 and the dd-MS2 were collected at a resolution of 30,000 and an isolation window of 0.4 m/z with a loop count of top 7. Stepped normalized collision energies of 10%, 30%, and 50% fragmented selected precursors in the collision cell. Dynamic exclusion was set at 7 seconds and ions with charges greater than 2 were omitted.

###### Data processing

Data processing steps included peak detection, spectral alignment, grouping of isotopic peaks and adduct ions, drift correction, and gap filling. Compound Discoverer V3.3 (ThermoFisher Scientific) was used to process the raw LC-MS data. Drift correction was performed on each individual feature, where a Systematic Error Removal using Random Forest (SERRF) method builds a model using the pooled QC sample peak areas across the batch and was then used to correct the peak area for that specific feature in the samples. Detected features were filtered with background and QC filters. Features with abundance lower than 5x the background signal in the sample blanks and that were not present in at least 50% of the QC pooled injections with a coefficient of variance (CV) lower than 80% (not drift corrected) and 50% (drift corrected) were removed from the dataset. Lipid annotations were accomplished based on accurate mass and relative isotopic abundances (to assign elemental formula), retention time (to assign lipid class), and MS2 fragmentation pattern matching to local spectral databases built from curated experimental data. Lipid nomenclature followed that described previously.^112^

###### Quality control procedures

System suitability was assessed prior to the analysis of each batch. A performance baseline for a clean instrument was established before any experiments were conducted. The mass spectrometers were mass calibrated, mass accuracy and mass resolution were checked to be within manufacturer specifications, and signal-to-noise ratios for the suite of IS checked to be at least 75% of the clean baseline values. For LC-MS assays, an IS mix consisting of 12 standards was injected to establish baseline separation parameters for each new column. The performance of the LC gradient was assessed by inspection of the column back pressure trace, which had to be stable within an acceptable range (less than 30% change). Each IS mix component was visually evaluated for chromatographic peak shape, retention time (lower than 0.2 minute drift from baseline values) and FWHM lower than 125% of the baseline measurements. The CV of the average signal intensity and CV of the IS (<=15%) in pooled samples were also checked. These pooled QC samples were used to correct for instrument sensitivity drift over the various batches using a procedure similar to that described by the Human Serum Metabolome (HUSERMET) Consortium (ref). To evaluate the quality of the data for the samples themselves, the IS signals across the batch were monitored, PCA modeling for all samples and features before and after drift correction was conducted, and Pearson correlations calculated between each sample and the median of the QC samples.

##### Metabolomics/Lipidomics Data filtering and normalization

The untargeted metabolomics datasets were categorized as either “named”, for chemical compounds confidently identified, or “unnamed”, for compounds with specific chemical properties but without a standard chemical name. While the preprocessing steps were performed on the named and unnamed portions together, only the named portions were utilized for differential analysis. The following steps are performed:

- Average rows that have the same metabolite ID.
- Merge the “named” and “unnamed” subparts of the untargeted datasets.
- Convert negative and zero values to NAs.
- Remove features with > 20% missing values.
- Features with < 20% missing values are imputed either using K-Nearest Neighbor imputation (for datasets with > 12 features) or half-minimum imputation (for datasets with < 12 features).
- All data are log2-transformed, and the untargeted data are median-MAD normalized if neither sample medians nor upper quartiles were significantly associated with sex or sex-stratified training group (Kruskal-Wallis p-value < 0.01). Note that for all log2 calculations, 1 is added to each value before log-normalization. This allows metabolite values that fall between 0 and 1 to have a positive log2 value.

Outlier detection was performed by examining the boxplot of each Principal Component and extending its whiskers to the predefined multiplier above and below the interquartile range (5x the IQR). Samples outside this range are flagged. All outliers were reviewed by Metabolomics CAS, and only confirmed technical outliers were removed.

##### Redundant Metabolite/Lipid Management

To address metabolites measured on multiple platforms (e.g. a metabolite measured on the HILIC positive and RP positive platforms), and metabolites with the same corresponding RefMet ID (e.g. alpha-Aminoadipic-acid, Aminoadipic acid both correspond to RefMet name ‘Aminoadipic acid’), we utilize the set of internal standards described above to make a decision on which platform’s measurement of a given feature to include in further analysis. For each tissue, based on whichever platform has the lowest coefficient of variation across all included reference standards for a given refmet id, that metabolite was chosen. The other platforms or metabolites, for this tissue, corresponding to this refmet id were removed from further downstream analysis. Data for all metabolites removed, including information about the coefficient of variation in the standards, normalized data, or differential analysis results, is available in the R Package *MotrpacHumanPreSuspensionAnalysis*, but is not loaded by default.

### Unsupervised multi-omic clustering

Non-negative matrix factorization (NMF)-based multi-omic clustering was used for participant-level clustering as previously described^113,114^. Data from RNA expression, proteome, and phosphoproteome from 152 baseline (pre-exercise) samples were used as input for the integrative multi-omic clustering. All data tables were concatenated and all rows with missing values were removed. To remove uninformative features from the dataset prior to NMF clustering, features with the lowest standard deviation (bottom 5th percentile) across all samples were removed. Each column in the data matrix was further scaled and standardized such that all features from different data types were represented as z-scores. The resulting data matrix of z-scores was converted into a non-negative input matrix, as required by NMF, as follows:

1. Create one data matrix with all negative numbers zeroed.
2. Create another data matrix with all positive numbers zeroed and the signs of all negative numbers removed.
3. Concatenate both matrices resulting in a data matrix twice as large as the original, but containing only positive values and zeros and hence appropriate for NMF.

The resulting matrix was then subjected to NMF analysis leveraging the NMF R-package^115^ and using the factorization method described in^116^. To determine the optimal factorization rank *k* (number of clusters) for the multi-omic data matrix, a range of clusters between *k* = 3 and 8 was tested. For each *k*, the matrix *V* was factorized using 50 iterations with random initializations of *W* and *H*. To determine the optimal factorization rank two metrics for each *k* were calculated: 1) the cophenetic correlation coefficient measuring how well the intrinsic structure of the data was recapitulated after clustering and 2) the dispersion coefficient of the consensus matrix as defined in^114^, measuring the reproducibility of the clustering across 50 iterations. The optimal *k* (*k =* 4) was defined as the maximum of the product of both metrics for cluster numbers between *k* = 3 and 8.

Having determined the optimal factorization rank *k*, in order to achieve robust factorization of the multi-omic data matrix *V*, the NMF analysis was repeated using 250 iterations with random initializations of *W* and *H*, and performed the partitioning of samples into clusters as described above.

A subset of samples, termed core samples, were identified for each cluster. For each sample, the difference between its highest cluster membership score and all other cluster membership scores was calculated. Core samples were classified as those samples for which the minimum cluster membership difference was greater than 1/*k* (1/4). Only core samples were used for the downstream statistical analyses. To test the association of the resulting clusters to clinical variables, Fisher’s exact test (R function fisher.test) was used to test for overrepresentation in the set of samples defining the cluster core.

The entire workflow described above has been implemented as a module in PANOPLY^117^, which runs on Broad’s Cloud platform Terra (https://app.terra.bio/). The docker container encapsulating the source code and required R-packages for NMF clustering has been submitted to Dockerhub (broadcptac/panoply_mo_nmf:1_6).

Enrichment of relevant gene sets was computed by projecting the matrix of signed multi-omic feature weights (Wsigned) onto Hallmark pathway gene sets^118^ using CAMERA-PR (see Analysis of Molecular Signatures). To derive a single weight for each gene measured across multiple omics data types, the weight with maximal absolute amplitude was retained.

### Metabolome association analysis

One-vs-rest association analysis^119^ was performed to identify differential metabolites across the baseline, mutli-omic NMF clusters. Fold-changes were computed for each one-vs-rest comparison, and then an F-test was performed across the four fold changes using limma. Metabolites with BH adjusted p-value < 0.01 were considered significant. The docker container encapsulating the source code and required R-packages for association analysis has been submitted to Dockerhub (broadcptac/panoply_association:1_6)

### Baseline and post-exercise response clinical associations

To identify significant associations between baseline clinical features and the baseline and exercise responses of omic features, we first limited our scope to human subjects in the EE or RE cohorts. We conducted this analysis on the transcriptomic, proteomic, phosphoproteomic, and metabolomic datasets. We constructed a matrix with rows the EE and RE subjects and the columns, 23 baseline clinical features measured prior to the acute exercise bout to gage each subject’s underlying physical fitness^70^.

For each ome analyzed, we also generated four separate omic value matrices with rows the omic features and columns the RE and EE subjects. One matrix represented baseline (pre-exercise) expression values for each feature in each EE and RE subject. The other three matrices reflected the three post-exercise timepoints, where values were the difference between pre and post-exercise for each subject. Exercise response matrices were limited to only include the subjects that contained both that time point and the pre-exercise time point for a given ome.

For each clinical feature x omic feature comparison, we constructed a linear model with covariates sex, age, exercise modality, BMI and clinical site ID using the R function *lm* to calculate association significance for a given comparison. To reduce the number of multiple hypotheses tested, we limited comparisons to features that already satisfied an unadjusted p-value of 0.01 for a two-sided Pearson correlation test, using the R function *cor.test*. p-values were adjusted using a Benjamini Hochberg correction with the R function *p.adjust*. Subjects with missing values for a given feature are removed from the clinical and omics data for the correlation testing and linear modeling.

We identified peak torque and end Watts as two clinical features that were most frequently significantly associated with gene expression, endwatts with respect to baseline gene expression and peaktorq with respect to 3.5 hour post exercise response. For each of these two comparisons, we separated the significant genes into those positively and negatively correlated and then used the *run_ORA* function in the *MotrpacHumanPreSuspensionAnalysis* R package to calculate pathway enrichment relative to a background set of the expressed genes in the muscle dataset.

### Statistical analysis

#### Differential analysis

To model the effects of both exercise modality and time, relative to non exercising control, each measured molecular feature was treated as an outcome in a linear mixed effects model accounting for fixed effects of exercise group (RE, EE, or CON), time point, as well as demographic and technical covariates (see covariate selection below for more info). Participant identification was treated as a random effect. For each molecular feature, a cell-means model is fit to estimate the mean of each exercise group-timepoint combination, and all hypothesis tests are done comparing the means of the fixed effect group-timepoint combinations. In order to model the effects of exercise against non-exercising control, a difference-in-changes model was used which compared the change from pre-exercise to a during or post-exercise timepoint in one of the two exercise groups to the same change in control. This effect is sometimes referred to as a “delta-delta” or “difference-in-differences” model. All comparisons were structured using ‘variancePartition::makeContrastsDream’.^76^ For full details on model and covariate selection, please refer to the multi-omics Landscape paper; in preparation/submission.

##### Missingness

Data missingness varied for multiple reasons including experimental design (i.e. temporal randomization which randomized some participants to have samples obtained at a subset of time points, see assay limitations (metabolomics and proteomics can have missingness as described in their individual sections). For some omic sets, imputation to alleviate missingness could be performed (metabolomics) but for others was not, including proteomics. Analysis demonstrated proteomics missingness was approximately at random (data not shown).

Given the above issues affecting the ultimate sample size for effect estimation, for a feature to be included in the analysis, a minimum number of 3 participants were required to have a paired pre-exercise sample for all groups and all during/post-exercise time points. Thus, all group comparisons (e.g. RE vs CON at 3.5 hours post-exercise) required 3 participants with matched pre- and post-exercise samples in each group. This requirement was satisfied in all omes and tissues except a subset of MS-acquired proteomics and phosphoproteomics features in skeletal muscle. Features not meeting this criterion in any group-time point set were excluded from differential analysis entirely, though raw and QC-normalized values are available.

##### Specific omic-level statistical considerations

Models for all omes other than methylation were fit using ‘variancePartition::dream’.^76^ For the transcriptomics and ATAC-seq, which are measured as numbers of counts, the mean-variance relationship was measured using ‘variancePartition::voomWithDreamWeights’ as previously described.^76^ For all other omes the normalized values were used directly as input to the statistical model.

The full implementation of the statistical models for each of non-methylation datasets can be found in the R Package *MotrpacHumanPreSuspensionAnalysis*. Methylation data was processed separately, and the methods can be found in the methylation methods section.

##### Significance thresholds

For each of the above contrasts, p-values were adjusted for multiple comparisons for each unique contrast-group-tissue-ome-time point combination separately using the Benjamini-Hochberg method to control False Discovery Rate (FDR).^120^ Features were considered significant at a FDR of 0.05 unless otherwise stated.

#### Human feature to gene mapping

The feature-to-gene map links each feature tested in differential analysis to a gene, using Ensembl version 105 (mapped to GENCODE 39)^121^ as the gene identifier source. Proteomics feature IDs (UniProt IDs) were mapped to gene symbols and Entrez IDs using UniProt’s mapping files.^122^ Epigenomics features were mapped to the nearest gene using the ChIPseeker::annotatePeak()^123,124^ function with Homo sapiens Ensembl release 105 gene annotations. Gene symbols, Entrez IDs, and Ensembl IDs were assigned to features using biomaRt version 2.58.2 (Bioconductor 3.18).^125–127^

For ATACseq and methyl capture features, relationship to gene and custom annotation provide gene proximity information and custom ChIPseeker annotations, respectively. Relationship to gene values indicate the distance of the feature from the closest gene, with 0 indicating overlap. Custom annotation categories include Distal Intergenic, Promoter (<=1kb), Exon, Promoter (1-2kb), Downstream (<5kb), Upstream (<5kb), 5’ UTR, Intron, 3’ UTR, and Overlaps Gene.

Metabolite features were mapped to KEGG IDs using KEGGREST, RefMet REST API, or web scraping from the Metabolomics Workbench, built on August 4th, 2026.^128,129^

#### Enrichment analysis

##### Preparation of differential analysis results

The differential analysis results tables for each combination of tissue and ome were converted to matrices of z-scores (derived from the limma moderated t-statistics) with either gene symbols, RefMet metabolite/lipid IDs, or phosphorylation flanking sequences as rows and contrasts as columns. These matrices serve as input for the enrichment analyses. For the transcriptomics, TMT proteomics, and Olink proteomics results, transcripts and proteins were first mapped to gene symbols. To resolve cases where multiple features mapped to a single gene, only the most extreme z-score for each combination of tissue, contrast, and gene was retained. Metabolite and lipid identifiers were standardized using the Metabolomic Workbench Reference List of Metabolite Names (RefMet) database^130^. For the CAMERA-PR enrichment of phosphoproteomics results, since peptides could be phosphorylated at multiple positions, phosphorylation sites were separated into single sites with identical row information (e.g., protein_S1;S2 becomes protein_S1 and protein_S2, both with the same data). For any combinations of contrast and phosphosite that were not uniquely defined, only the most extreme z-score (maximum absolute value) was selected for inclusion in the matrix. Singly phosphorylated sites were required to use the curated kinase–substrate relationship information available from PhosphoSitePlus (PSP),^131^ described below.

##### Gene set selection

Gene sets were obtained from the MitoCarta3.0 database, CellMarker 2.0 database [PMID: 36300619], and the C2-CP (excluding KEGG_LEGACY) and C5 collections of the human Molecular Signatures Database (MSigDB; <u>v2023.2.Hs</u>)^118,132,133^. Metabolites and lipids were grouped according to chemical subclasses from the RefMet database^130^. This includes subclasses such as “Acyl carnitines” and “Saturated fatty acids”.Flanking sequences for protein phosphorylation sites were grouped according to their known human protein kinases provided in the *Kinase_Substrate_Dataset* file from PhosphoSitePlus (PSP; v6.7.1.1; https://www.phosphosite.org/staticDownloads.action; last modified 2023-11-17).^131^ In addition to these kinase sets, sets from the directional Post-Translational Modification Signatures Database (PTMsigDB) were included for analysis.^134^

For each ome, molecular signatures were filtered to only those genes, metabolites/lipids, or flanking sequences that appeared in the differential analysis results. After filtering, all molecular signatures were required to contain at least 5 features, with no restriction on the maximum size of sets. Additionally, gene sets were only kept if they retained at least 70% of their original genes to increase the likelihood that the genes that remain in a given set are accurately described by the set label.

##### Analysis of Molecular Signatures

Analysis of molecular signatures was carried out with the pre-ranked Correlation Adjusted MEan RAnk (CAMERA-PR) gene set test using the z-score matrices described in the “Preparation of differential analysis results” section to summarize the differential analysis results for each contrast at the level of molecular signatures.^16,134^ Z-scores were selected as the input statistics primarily to satisfy the normality assumption of CAMERA-PR, and the analysis was carried out with the *cameraPR.matrix* function from the TMSig R/Bioconductor package (https://www.bioconductor.org/packages/release/bioc/html/TMSig.html).

##### Statistical significance thresholds

For both CAMERA-PR and ORA results, p-values were adjusted within each combination of tissue, ome, contrast, and broad molecular signature collection (MitoCarta3.0, CellMarker 2.0, C2, C5, RefMet, PSP, and PTMsigDB) using the Bejamini-Hochberg method. Gene sets and RefMet subclasses were declared significant if their adjusted p-values were less than 0.05. This threshold was raised to 0.1 for kinase sets.

##### Visualization methods

Bubble heatmaps were generated from the enrichment analysis results using the *enrichmap* function from the TMSig R/Bioconductor package (https://www.bioconductor.org/packages/release/bioc/html/TMSig.html).^135,136^

#### PLIER methods

To delve into cross-omic temporal trends in muscle acute exercise response, we applied PLIER^137^, which takes an expression matrix of features vs samples as input and outputs patterns of activity, in the form of latent variables (LVs), and connects these patterns with user-provided prior information, such as reactome or KEGG pathways. We use the PLIER package R function *num.pc* to identify the number of significant principle components for the singular value decomposition (SVD) of the data and specify double this number as the desired LV count. If *num.pc* is computationally infeasible, we use a default of 100 LVs for a PLIER run, with rigorous testing of alternative LV depth to ensure coverage of the significant discoverable exercise patterns. The ATAC-RNA-Protein cross-omic PLIER analysis specified the default 100 LVs, as did the RNA-Phosphoproteome cross-omic PLIER analysis. PLIER works to identify a set of LVs that best account for all patterns identified in the dataset, optimizing individual LV trajectories over the samples and feature associations to minimize the difference between normalized input data and PLIER LV output, while also accounting for feature associations with prior knowledge in the form of pathway enrichments.

We ran PLIER separately on each cross-omic combination. For the ATAC-RNA-Protein cross-omic PLIER, we identified proteins present in the protein abundance data that were also included in the RNAseq data and contained at least one annotated promoter peak in the ATACseq data. For the ATACseq data, we generated a cumulative accessibility score for the promoter region of each protein’s associated gene. For the RNA-Phosphoproteome cross-omic PLIER, we identified proteins with phosphosites present in the phosphoproteomic data that were also included in the RNAseq data. Similar to the ATACseq data, we generated a cumulative phosphorylation score for each protein. In each cross-omic PLIER case, we then generated a cross-omic matrix of exercise response, z-scoring the normalized data matrix for each ome. Following by-ome z-scoring, the z-scored matrices are concatenated into one large matrix with the rows corresponding to the consensus features across all omes, and columns the cumulative samples from each ome. Significance of an LV to differentiate resistance or endurance exercise from control at a given time point relative to pre-exercise is determined by comparing the differences in LV value post-exercise vs pre-exercise for each subject at that time point separately between resistance and control and endurance and control within each ome and adjusting for multiple hypotheses using the *compare_means* R function.

#### PTM-SEA

PTM-SEA is the name given to single sample gene set enrichment analysis (ssGSEA) used in conjunction with PTMsigDB.^134^ Each signature is assigned an enrichment score (ES), which classically quantifies the degree of association with a particular sample profile, though we applied PTM-SEA at the contrast level. It utilizes permutation of feature labels to calculate p-values and normalized enrichment scores (NES).

PTM-SEA was performed on the z-scores from muscle and adipose phosphoproteomics separately. The dataset was filtered for confidently localized sites (confident_score > 17) prior to this analysis. The first step of PTM-SEA creates a single-site dataset from the initial multi-site dataset using the flanking sequence and protein grouping information (preprocessGCT). This was run with the following parameters: id_type_out = “seqwin”, organism = “human”, mode = “abs.max”, seqwin_column = “flanking_sequence”, gene_symbol_col = “gene_symbol”, id_type = “sm”, acc_type_in = “uniprot”, residue = “S|T|Y”, ptm = “p”, and localized = FALSE.

After pre-processing, PTM-SEA was performed on the single-site level datasets using PTMSigDB v2.0 with the following parameters: weight = 0, statistic = ”area.under.RES”, output.score.type = “NES”, nperm = 1000, global.fdr = FALSE, and min.overlap = 5. Since weight = 0, parameters sample.norm.type and correl.type do not affect the analyses.

PTM-SEA was run using the v0.5.2 Dockerfile (gcr.io/broadcptac/ptm-sea:0.5.2). All source code and definitions of parameters are available on Github: https://github.com/broadinstitute/ssGSEA2.0.

#### SC-ION methods

Spatiotemporal Clustering and Inference of Omics Networks (SC-ION, https://github.com/nmclark2/SCION)^45^ was applied to the dataset to infer unsupervised multi-omics networks of exercise response. The source code for SC-ION was adapted for MoTrPAC such that the SC-ION analysis can be repeated or repurposed for additional data within this study (see Package assembly and distribution). The exact parameters used to generate results for the figures presented in the manuscript are found in the figures folder in the motrpac-human-presuspension-acute repository.

##### Selecting regulator and target data

SC-ION takes as input two data matrices where one matrix is assigned as the “regulator” matrix and the other is assigned as the “target” matrix. Phosphosites on transcription factors (TFs) in were used as the regulators, and DA transcripts were used as the targets. EE and RE samples were separated, and one network was inferred for each modality. Control samples were excluded from the SC-ION analysis.

##### Preparation of input data

The normalized values for the muscle phosphoproteome were used as the regulator matrix. Since SC-ION cannot incorporate missing values, the muscle phosphoproteome was filtered for phosphosites with no more than 40% missing values, and then missing values were imputed with Multiple Imputation by Chained Equations (MICE)^138^ by using the average imputed value from 15 multiple imputations.This matrix was filtered for phosphosites located on TFs using “The Human Transcription Factors” database (https://humantfs.ccbr.utoronto.ca/). The normalized values for the transcriptome were used as the target matrix. The target matrix was filtered for DA transcripts (adjusted p-value < 0.05 in any exercise-with-control contrast). All matrices were split into EE and RE samples prior to running SC-ION.

##### Clustering

SC-ION incorporates an optional clustering step to group together features with similar expression profiles prior to inferring the network. For this implementation, c-means clustering was used to cluster only the regulator and target features used as inputs to each network. The z-score matrices described in the Enrichment Analysis Methods were used as input for fuzzy c-means (FCM) clustering.^139,140^ The regulator and target matrices were stacked to form a single matrix. The rows of this matrix were then divided by their sample standard deviations, which were calculated after two columns of zeros were temporarily included to represent the pre-exercise timepoints; these zero columns were discarded before clustering. These values were zero-scaled prior to clustering. The Mfuzz *R* package was used to perform FCM clustering with the parameter *m* set to the value of *Mfuzz::mestimate* for the normalized and scaled matrix^136,141,142^. The minimum distance between centroids was used as the cluster validity index to determine the optimal number of clusters, which was determined to be 12.

##### Edge trimming

All of the edges in the inferred networks are assigned a weight which indicates the relative confidence in the prediction. To trim the networks based on edge weights, a permutation analysis was performed by randomly shuffling the regulator and target matrices and running SC-ION 100 times. The edge weights were then ranked and assigned a *p*-value based on a permutation test. The *p*-values were then adjusted using Benjamini-Hochberg (BH). The networks were originally trimmed to edge weights with a BH-adjusted *p*-value < 0.05 as determined by the permutation test. However, this resulted in overly dense networks, so a more conservative edge weight cutoff (edge weight > 0.1) was chosen based on the edge weight distribution.

#### MAGICAL methods

MAGICAL^143^ was used to infer cis-regulatory circuits of differentially expressed genes in response to acute exercise. A cis-regulatory circuit consists of a transcription factor (TF), a cis-regulatory region on the chromatin where the TF binds to, and a target gene whose expression is regulated by the TF binding to the cis-regulatory region. MAGICAL infers cis-regulatory circuits with a Bayesian framework which identifies correlated patterns of chromatin accessibility and gene expression from paired ATAC-seq and RNA-seq of the same participants across time points. MAGICAL also infers transcription factor activities and their binding to the identified cis-regulatory regions. For each gene, MAGICAL identifies cis-regulatory regions within a genomic window of +/- 100kb around the transcription start site of the gene. To focus the analysis on identifying cis-regulatory circuits relevant to physical exercise, MAGICAL was used only on the chromatin peaks with differential accessibility (uncorrected p-value < 0.005) and genes with differential expression (adjusted p-value < 0.05) in at least one timepoint. The less stringent cutoff for the differential peaks was due to the lower strength of signal in the ATAC-seq assay; additionally, this analysis focuses on identifying cross-individual correlation patterns and does not rely on a stringent differential cutoff.

#### Integration of SC-ION and MAGICAL networks

##### Network integration

The EE SC-ION, RE SC-ION, and MAGICAL networks were integrated by taking the union of all inferred edges. Therefore, any edge inferred by SC-ION or MAGICAL is retained in the final network. If the same edge is predicted by both SC-ION or MAGICAL, it is reported twice (Table S6).

##### Importance score calculation

A Network Motif Score (NMS) was calculated for each feature in the combined SC-ION+MAGICAL network as described in Clark et al.^45^ Feed-forward loop (FFL), diamond, and 3-chain motifs were used as input for the NMS (Figure 6A). The total number of motifs, as well as the features within each motif, were determined using the NetMatchStar app^144^ in Cytoscape. The NMS for each network feature is reported in Table S6.

##### Visualization

All network visualization on the combined SC-ION+MAGICAL network was performed in Cytoscape.

### Quantification and Statistical Analysis

#### Statistical parameter reporting

All relevant statistical parameters for the analyses performed in this study such as sample size (*N*), center and spread (e.g. mean/median, standard deviation/error), statistical methodology (e.g. mixed-effects linear model) and significance cutoffs (e.g. adjusted *p*-value < 0.05) are reported in the main text, figure legends, STAR Methods, and/or supplementary information. Where appropriate, methods used to determine the validity of certain statistical assumptions are discussed.

#### Statistical limitations

This report presents analyses and results for 176 participants randomized before the suspension of the MoTrPAC study due to the COVID-19 pandemic. The main post-suspension MoTrPAC study will include over 1,500 participants randomized under a slightly modified protocol.^14^ The current paper focuses on evaluating feasibility and generating hypotheses for the main study. Results should be interpreted with caution for several reasons: 1) small sample sizes reduce the power to detect even moderate effects;^145^ 2) simple randomization of small groups can create imbalances in both known and unknown confounders;^146,147^ and 3) NIH initiatives have long emphasized caution in interpreting small studies to enhance reproducibility.^148^

Although blocked randomization with site-based stratification was employed, discrepancies in regulatory approvals, start-up times, and pandemic-related interruptions resulted in imbalanced and small sample sizes. Specific limitations in the baseline (pre-intervention) data include: 1) subgroup sample sizes based on intervention group, sex, and timepoint as small as 4 participants; 2) limited generalizability, as 50% of controls with biosamples were randomized at two of the ten sites, and 74% of participants at four sites; and 3) "sex differences" may reflect "body composition differences" due to insufficient data to disentangle sex from body composition, and variations in DXA machine operators and brands across sites.

Testing for heterogeneity of response in small subgroups was largely avoided. As Brookes et al.^149^ show, interaction effects must be at least double the main effect to achieve 80% power. While the EE and RE groups each had around 70 participants with biosamples, providing 80% power to detect a 0.5 effect size (difference in means/SD) using a two-sample t-test (alpha = 0.05, two-sided), for tests of interaction effects to have 80% power an interaction effect size > 1 would be required, which is considered large by Cohen.^150^ Consequently, heterogeneity of response was explored only descriptively within the larger randomized groups (EE/RE), with some inferential statistics (e.g., confidence intervals, p-values) emphasizing interval estimation. Our aim was to present the results from a hypothesis-generating perspective, following Ioannidis’s cautionary guidance,^14^ and to lay a solid foundation for future analyses in the main MoTrPAC study.

### Additional Resources

#### Package assembly and distribution

The *MotrpacHumanPreSuspensionAnalysis* R package (https://github.com/MoTrPAC/MotrpacHumanPreSuspensionAnalysis) contains functions to generate visualizations, as well as data objects for differential abundance analysis, summary statistics for normalized expression data, feature to gene mapping, and molecular signature datasets as described in the methods.

The *MotrpacHumanPreSuspensionData* R package (https://github.com/MoTrPAC/MotrpacHumanPreSuspensionData) is a package containing individual level data, which cannot be made openly available and can be accessed with approval from a data access committee. See motrpac-data.org for details.

Data used in the preparation of this article were obtained from the Molecular Transducers of Physical Activity Consortium (MoTrPAC) database, which is available for public access at motrpac-data.org. The specific collection versions are: quant-id/human-precovid/c1.0, analysis/human-precovid-sed-adu/c2.0, phenotype/humanprecovid-sed-adu/c3.0.

Lastly, the MotrpacPreSuspensionAcute GitHub repository (https://github.com/MoTrPAC/motrpac-human-presuspension-acute) contains code and individual parameters for each figure panel, utilizing the MotrpacHumanPreSuspensionAnalysis and MotrpacHumanPreSuspensionData packages.

## Acknowledgements

The MoTrPAC Study is supported by NIH grants U24OD026629 (Bioinformatics Center), U24DK112349, U24DK112342, U24DK112340, U24DK112341, U24DK112326, U24DK112331, U24DK112348 (Chemical Analysis Sites), U01AR071133, U01AR071130, U01AR071124, U01AR071128, U01AR071150, U01AR071160, U01AR071158 (Clinical Centers), U24AR071113 (Consortium Coordinating Center), U01AG055133, U01AG055137, U01AG055135, U01AG070959, U01AG070960, and U01AG070928 (Pre-Clinical Animal Sites).

## Author Information

### MoTrPAC Study Group

Hiba Abou Assi, Cheehoon Ahn, David Amar, Mary Anne S. Amper, Brian J. Andonian, Euan A. Ashley, Isaac K. Attah, Jacob L. Barber. Alicia Belangee, Bryan C. Bergman, Daniel H. Bessesen, Kevin Bonanno, Gerard A. Boyd, Anna R. Brandt, Nicholas T. Broskey, Thomas W. Buford, Toby L. Chambers, Clarisa Chavez Martinez, Maria Chikina, Alex Claiborne, Zachary S. Clayton, Clary B. Clish, Katherine A. Collins-Bennett, Dan M. Cooper, Tiffany M. Cortes, Gary R. Cutter, Matthew Douglass, Sara E. Espinoza, Charles R. Evans, Facundo M. Fernandez, Johanna Y. Fleischman, Daniel E. Forman, Will A. Fountain, David A. Gaul, Yongchao Ge, Robert E. Gerszten, Catherine Gervais, Aaron H. Gouw, Kevin J. Gries, Marina A. Gritsenko, Fadia Haddad, Joshua R. Hansen, Trevor Hastie, Zhenxin Hou, Fang-Chi Hsu, Ryan P. Hughes, Chelsea M. Hutchinson-Bunch, Olga Ilkayeva, Byron C. Jaeger, John M. Jakicic, Catherine M. Jankowski, Pierre M. Jean-Beltran, David Jimenez-Morales, Neil M. Johannsen, Johanna L. Johnson, Maureen T. Kachman, Erin E. Kershaw, Wendy M. Kohrt, Dillon J. Kuszmaul, Damon T. Leach, Bridget Lester, Minghui Lu, Colleen E. Lynch, Nada Marjanovic, Sandra T. May, Edward L. Melanson, Nikhil Milind, Michael E. Miller, Matthew E. Monroe, Cristhian Montenegro, Stephen B. Montgomery, Ronald J. Moore, Kerrie L. Moreau, Charles C. Mundorff, Venugopalan D. Nair, Masatoshi Naruse, Bradley C Nindl, German Nudelman, Nora-Lovette Okwara, Eric A. Ortlund, Vladislav A. Petyuk, Paul D. Piehowski, Hanna Pincas, David Popoli, Wei-Jun Qian, Shlomit Radom-Aizik, Tuomo Rankinen, Prashant Rao, Abraham Raskind, Alexander Raskind, Blake B. Rasmussen, Ulrika Raue, Eric Ravussin, R. Scott Rector, W. Jack Rejeski, Joseph Rigdon, Stas Rirak, Ethan Robbins, Jeremy M. Robbins, Margaret Robinson, Kaitlyn R. Rogers, Renee J. Rogers, Jessica L. Rooney, Tyler J. Sagendorf, Irene E. Schauer, Robert S. Schwartz, Courtney G. Simmons, Chad M. Skiles, Kevin S. Smith, Michael P. Snyder, Tanu Soni, Maja Stefanovic-Racic, Cynthia L. Stowe, Andrew M. Stroh, Yifei Sun, Kristen J. Sutton, Robert Tibshirani, Russell Tracy, Todd A. Trappe, Mital Vasoya, Nikolai G. Vetr, Caroline S. Vincenty, Elena Volpi, Alexandria Vornholt, Martin J. Walsh, Matthew T. Wheeler, Katie L. Whytock, Angela Wiggins, Yilin Xie, Gilhyeon Yoon, Jay Yu, Xuechen Yu, Elena Zaslavsky, Bingqing Zhao, Jimmy Zhen

## MoTrPAC Acknowledgements

Nicole Adams, Abdalla Ahmed, Andrea Anderson,Carter Asef, Arianne Aslamy, Marcas M. Bamman, Jerry Barnes, Susan Barr, Kelsey Belski, Will Bennett, Kevin Bonanno, Amanda Boyce, Brandon Bukas, Emily Carifi, Chih-Yu Chen,Haiying Chen, Shyh-Huei Chen, Samuel Cohen, Audrey Collins, Gavin Connolly, Elaine Cornell, Julia Dauberger, Carola Ekelund, Shannon S. Emilson, Karyn A. Esser, Jerome Fleg, Nicole Gagne, Mary-Catherine George, Ellie Gibbons, Jillian Gillespie, Laurie Goodyear, Aditi Goyal, Bruce Graham, Xueyun Gulbin, Jere Hamilton, Leora Henkin, Andrew Hepler, Andrea Hevener, Lidija Ivic, Ronald Jackson, Andrew Jones, Lyndon Joseph, Leslie Kelly, Ian Lanza, Gary Lee, Jun Li, Adrian Loubriel, Kristal M. Maner-Smith, Ryan Martin, Padma Maruvada, Alyssa Mathews, Curtis McGinity, Lucas Medsker, Kiril Minchev, Samuel G. Moore, Michael Muehlbauer, K Sreekumaran Nair, Anne Newman, John Nichols, Concepcion R. Nierras, George Papanicolaou, Lorrie Penry, June Pierce, Megan Reaves, Eric W. Reynolds, Jeremy Rogers, Scott Rushing, Santiago Saldana, Rohan Shah, Samiya M. Shimly, Cris Slentz, Deanna Spaw, Debbie Steinberg, Suchitra Sudarshan, Alyssa Sudnick, Jennifer W. Talton, Christy Tebsherani, Nevyana Todorova, Mark Viggars, Jennifer Walker, Michael P. Walkup, Anthony Weakland, Gary Weaver, Christopher Webb, Sawyer Welden, John P. Williams, Marilyn Williams, Leslie Willis, Yi Zhang

