## Supplementary Results for "Integrative Multi-omics Analysis of the Human Skeletal Muscle Response to Endurance or Resistance Exercise: Findings from the Molecular Transducers of Physical Activity Consortium (MoTrPAC)"

### *Multi-omic associations with baseline physiological features*

We examined molecular enrichment in each NMF cluster using Pre-ranked Correlation-Adjusted MEan-RAnk (CAMERA-PR)[16] (Extended Data Fig. 1c). Clusters 3 and 4, skewed for younger age, were positively enriched in oxidative phosphorylation compared to the older clusters. Although clusters 1 and 2 separate into older age groups, cluster 1 displayed positive enrichment in inflammatory pathways, including complement and innate immune signaling pathways. Additionally, these inflammatory pathways trended lower in cluster 2 (older women) compared to cluster 1 (inflammation enriched) and cluster 4 (younger women).

Baseline metabolomic profiles also differed across the NMF clusters (Extended Data Fig. 1d). Several amino acids, energy-associated metabolites, and lipid species showed the strongest separation. Cluster 3 (younger men) had higher baseline abundance of muscle- and metabolism-related metabolites such as carnosine, citrulline, branched-chain amino acids (leucine/isoleucine), histidine-related metabolites (His, His-Ala), carnitine-related metabolites (carnitine, 3-Dehydroxycarnitine), and other polar metabolites such as ectoine and imidazolepropionic acid. Conversely, several triglyceride species were relatively depleted in Cluster 3 and showed higher abundance in Clusters 1 and 2. Cluster 2 (older women) showed higher abundance of lipid remodeling metabolites, particularly ether PCs and several triglyceride species, as well as nicotinamide-associated metabolites such as 1-methylnicotinamide and trigonelline. Cluster 4 (younger women) showed a distinctive lipid profile with higher abundance of ether PEs, lysophospholipids, diacylglycerols, and selected ceramide species. .

In ome to clinical feature association analysis (see Methods) we identified 4251 genes (25.7%), 2368 proteins (38.0%), 5529 phosphosites (29.8%), and 376 metabolites (41.7%) that, for at least one timepoint comparison, were significantly associated with a clinical feature (Extended Data Fig. 2d-g, Supplementary Table 2). Fifty-nine of the 182 positively associated genes (32.4%, hypergeometric test p-value $< 0.0001$) were also positively associated in the proteomics data, 35 of which were directly annotated to aerobic respiration, including ATP5PB (Extended Data Fig. 2h). Genes negatively correlated with Watts were instead enriched for semaphoring signaling and RHO GTPASE effectors.

### *Differential Abundance Analysis*

The 15 min post-exercise time point displayed the greatest number of EE and RE concurrent features, and consisted primarily of phosphoproteomic, followed by transcriptomic and epigenomic (ATAC-seq) features. Subsequently, the number of modality convergent features decreased throughout the post-exercise time course and was driven mainly by the transcriptome. Overrepresentation analysis (ORA) of the shared EE and RE DA features revealed early (15 min post-exercise) enrichments in muscle tissue development and contraction terms, followed by inflammatory and immune system-related pathways at 3.5 hours and mitochondrial pathways at 24 hours post-exercise (Fig. 2b; Supplementary Table 3).

Further, consistent with a more robust multi-omic response to RE, the majority of DA features were specific to the three post RE time points.

Modality divergent changes in skeletal muscle acylcarnitines highlight the underlying differences in metabolic pathways, i.e., glycolytic vs. oxidative to support RE and EE, respectively. Yet, both modalities induced an increase in short chain (C3-C5) acylcarnitine species (Extended Data Fig. 3b), that are derived from non-lipid sources as products of metabolism of branched chain amino acids (BCAAs) and other amino acids (e.g. CAR 3, CAR 5) and ketones (CAR 4;OH).[18] Both EE and RE led to reductions in triacylglycerol (TG) species at 3.5 hours which was sustained in EE at 24 hours, consistent with muscle TG oxidation during acute exercise[151] and an enhanced capacity for lipid catabolism following EE (Fig. 3d). Hydroxy fatty acids rose at 3.5 hours following both exercise modalities (Fig. 3f) which could reflect increased ketone body utilization reflecting the fasting state following the exercise bout.

Overall, the 24-hour time point revealed distinct, modality-specific changes in the proteome, although relatively few proteins were significantly altered. Of the 31 proteins that changed, only four proteins were specific to EE, including ALAS1, a mitochondrial enzyme essential for initiating heme biosynthesis, which directly supports oxidative phosphorylation capacity[152] (Extended Data Fig. 3c). Glycogenin-1 (GYG1), a glycogen synthesis protein, was the only protein that increased early and stayed upregulated until the 3.5 hour time point following EE only. RE-specific proteins included CCN family member 1 (CCN1), a protein with roles in wound healing and angiogenesis [153] as well as several involved in muscle development, differentiation, and striated muscle contraction including ALPK3 and NEXN.[154,155] Notably, Ankyrin repeat domain 1 (ANKRD1), a mechanosensitive transcription co-regulator protein and part of titin-N2A complex, that is implicated in muscle remodeling, displayed delayed but prolonged upregulation at both the gene and protein levels following RE (Extended Data Fig. 5e).[156,157] In summary, the multi-omic analysis suggests that later time points exhibit greater exercise modality divergence, and that RE exerts a generally larger and prolonged influence on the multi-omic landscape of skeletal muscle.

We anticipated that a portion of these DA features would be associated with physiological phenotypes measured during an initial screening test prior to the acute exercise bout (Extended Data Fig. 2a). We found that 2183 identified DEGs were also significantly associated with a clinical feature, including 116 whose 3.5 hour exercise response was linked with knee extension peak torque. This includes GPATCH3 (Extended Data Fig. 3d), a critical alternative splicing co-regulator along with DHX15[158], both of which are DEGs 3.5 hours post RE. The 3.5 hour protein abundance response to OGFOD1, a promoter of cardiac hypertrophy[159] and a differential protein post RE, was also associated with knee extension peak torque (Extended Data Fig. 3e). Fifty-eight identified differential metabolites were also significantly associated with a clinical feature, 24 of which were acyl carnitines. The baseline expression of 19 of these acyl carnitines were associated with initial screening average respiratory exchange ratio (RER) and the 3.5 hour exercise response of 14 of the acyl carnitines were associated with peak power output (Watts), including CAR 8:0 which was significantly associated in each comparison (Extended Data Fig. 3f,g). Among the 1044 overlapping differential phosphosites and those significantly associated to clinical features, 43 of them were located on NACA, a muscle-specific transcription factor and co-activator of JUN[160], one of the early response DEGs post EE and RE in muscle. The 24 hour exercise response of 11 of the NACA phosphosites were associated with screening test end systolic BP (Extended Data Fig. 3h,i).

Comparing the muscle transcriptomic results of this study with a previous study of the gene expression changes following both EE and RE[62] identified a large degree of time point-specific consistency in exercise response for both modalities (Extended Data Fig. 3j). While the exact post-exercise time points were not the same, we observed high correlation between the changes at 15 minute and 3.5 hour time points in our study with the changes at the immediate post-exercise and 3 hour time points in the comparison study,[62] respectively.

### *Modality-specific responses to acute exercise are linked to metabolic and mechanical stimuli*

From the transcriptomic CAMERA-PR enrichment analyses, despite numerous EE and RE shared enrichments, the magnitude and duration of responses were often greater with RE (Fig. 3a). Early EE transcriptomic enrichments were commonly  attenuated by the intermediate timepoint compared to prolonged upregulation with RE, as exemplified by angiogenic VEGFR2 signaling and the integrin 3 pathways (Fig. 3a). Integrin 3 regulates muscle repair following injury by promoting myogenesis and regulating macrophage polarization,[161,162] its prolonged enrichment with RE suggests its greater muscle remodeling signature. Despite early overlap in transcriptomic enrichments, the majority of enriched pathways were modality-divergent at 24 hours, where EE resulted in a robust enrichment in mitochondrial metabolism-related pathways including cristae formation, FAO and mitochondrial fission (Fig. 3a). To further examine mitochondria-specific responses, we queried the MitoCarta 3.0[133] database for transcript enrichments (Fig. 3b, Supplementary Table 4). Both modalities induced an early downregulation of mitochondrial transcriptomic pathways reflecting earlier studies[62]. Enrichment analysis using MitoCarta 3.0  further highlighted EE induced a widespread increase in mitochondrial transcriptomics pathways including, lipid and branched chain amino acid metabolism, cristae formation, and detoxification (Fig. 3b). We did observe some mitochondrial transcriptomic enrichments following RE at 24 hours, albeit much attenuated compared to EE (Fig. 3b). Modality-specific transcriptomic enrichments included unique transcriptomic enrichment of autophagy, contractile fiber, antiviral mechanisms via IFN and translation initiation at 3.5 hours following RE, whereas EE induced unique enrichment of FAO at 3.5 hours, and regulation of mitochondrial gene expression and glycolysis/gluconeogenesis at 24 hours post-exercise (Fig. 3a). Even for shared pathways, the genes driving enrichments often differed between modalities. For example, in the hypertrophy model pathway (Extended Data Fig. 4a), the key skeletal muscle transcriptional regulator *NR4A3* was upregulated in response to both EE and RE, whereas myogenin (*MYOG*), a transcription factor that promotes myocyte fusion and activates muscle-specific gene expression,[163] was increased in RE at 3.5 and 24 hours, but decreased with EE at 3.5 hours [15] (Extended Data Fig. 4a).

At 3.5 hour post exercise, both exercise modalities induced enrichment of terms related to class III histone deacetylases (HDAC3) and PPAR gamma coactivator 1-alpha (*PPARGC1A*) activation by phosphorylation (Fig. 3a), key pathways involved in fatty acid partitioning.[164] While both modalities increased *PPARGC1A* expression, EE induced unique upregulation of PPAR-alpha while RE induced a more robust upregulation of PPAR-delta at 3.5 hours that uniquely decreased below baseline levels at 24 hours (Extended Data Fig. 4e).

Probing the MitoCarta 3.0 enrichment database, we further observed that RE suppressed mitochondrial proteomic pathway enrichments throughout th epost-exercise recovery period up to 24 hours, whereas mitochondrial protein enrichments were only suppressed immediately following EE (Extended Data Fig. 4c). Together these data suggest that mitochondrial transcriptional adaptations begin during the intermediate post-exercise timepoint, and that changes in protein abundance likely occur beyond 24 hours, given that mitochondrial transcriptomic enrichments peaked at this late timepoint (Fig. 3b, Extended Data Fig. 4c).

### *PTM-SEA enrichment analysis of the phosphoproteome*

Post translational modification signature enrichment analysis (PTM-SEA) showed that EE and RE signatures were broadly shared at the early timepoint and included IL-11, oxytocin, TWEAK and T-cell receptor pathway enrichments, EGF perturbagen signal, and kinase signatures such as PRKA and PRKD, among others (Fig. 5a). EE-specific signatures included CAMK2G, shown to promote MEF2A-mediated GLUT4 transcription with exercise[165], and the IL-6 pathway. IL-6 secretion following acute exercise is linked to glycogen depletion and has been previously documented to be more pronounced following acute EE[166,167] pairing with CAMK2G enrichment (Extended Data Fig. 6a). RE specific signatures included multiple proinflammatory cytokines such as TNF, IL-1B, and IL-1A, and the stress-associated corticotropin-releasing hormone (CRH) pathway among multiple stress-inducible MAPK pathways. RE also induced an early increase in the angiogenic NP TIE2 pathway (Extended Data Fig. 6d), which included HSPB1 (HSP27) (Extended Data Fig. 6e), whose stronger increase in the phosphoproteome in RE is consistent with prior studies,[168] indicating greater induction of stress response signatures following RE[15]. These findings may represent a compensatory response to RE in the presence of mechanical stress and decreased oxygen availability, linking these phosphoproteomic changes to our observed differential substrate utilization pathways in the acute EE or RE response.

### *Central regulatory nodes of the acute exercise response connect with modality divergent transcriptional effects*

MEF2A was identified as a key regulator of the exercise response consistent with previous exercise studies.[169] In exercise training, recurrent MEF2A activation promotes mitochondrial adaptations and coordinates myoblast fate commitment, differentiation and remodelling.[170,171] We filtered the predicted targets of MEF2 in the combined network to only those which show evidence of MEF2A binding via ChIP-seq (Fig. 6b) (211/632 predicted targets, 33.4%).[50,172] The majority of these targets are predicted to be regulated by one of two different forms of phosphorylated MEF2A: the singly-phosphorylated at S98 which decreased at 15 minutes facilitates its transcriptional activity in a cell dependent context, where its dephosphorylation favors MEF activation in myoblasts,[173] yet the doubly-phosphorylated S98T108 which increases at 15 minutes (Fig. 6c). We performed ORA on the MEF2A targets whose transcript expression significantly changed in response to EE, RE, or both modalities at any timepoint and found several significant terms including pathways related to muscle cell development (Extended Data Fig. 7b), consistent with known roles of MEF2 family TF in skeletal muscle repair[174]. Several MEF2A-targeted genes related to autophagy and VEGFA/VEGFR2 signaling were upregulated following acute exercise in both EE and RE, although the magnitude and trajectories of gene expression differed between modalities, with RE upregulating a greater number of both autophagy and VEGFR2-related genes (Fig. 6d,e). For example, the MEF2A-targeted autophagy genes induced by RE included *BAG3*, which contained HIPK3-associated phosphosites (Fig. 5d), and cyclic AMP-dependent transcription factor (*ATF6*) which regulates the unfolded protein response. Additionally, within VEGFR2 signaling enrichment of CCN1 was conserved across exercise modalities, yet displaying a prolonged enrichment with RE. The role of CCN1 as an exkerine is further explored in (ref landscape). At the transcriptomic level, RE also displayed prolonged enrichment in pathways related to protein folding chaperones, supportive of greater protein remodeling in the 24 hr recovery from acute RE (Fig. 3a). Together, this suggests that MEF2A regulates coordination of angiogenesis, remodeling, and cellular clearance in the post-acute exercise period, with modality-specific transcriptional programming.

### *Supplementary Discussion*

We observed EE-dominant CAMK signaling activation, which is induced with exercise and facilitates MEF2 transcriptional activation through the repression of the inhibitory role of class II HDACs,[175] which may be a contributory driver of mitochondrial transcriptional programming with EE. The decrease in ceramide species coupled with a more robust increase in the ceramidase, *ASAH1*, with EE may be linked to training-induced improvements in insulin sensitivity; reductions with acute exercise in this data suggest mechanisms by which a single bout of acute EE can enhance insulin sensitivity.[176,177] The increased drive of fatty acid metabolism following EE were likely reflected by the increased expression of genes related to FAO and mitochondrial components at 3.5 and 24 hours following the acute bout, respectively, and an EE specific induction of *PPARA* at 24 hours, which promotes FAO.[178]

MEF2A activation has been suggested to be a primary regulator of skeletal muscle remodeling, for example we previously observed it to be a key TF in striated muscle responses to exercise training in rats.[179] Additionally, regulatory network analysis identified other known central regulators of exercise responses, including HES1, ATF3 and ESRRA.[62,180,181] Through regulatory network analysis (SC-ION), we identify autophagy and VEGFA/VEGFR2 as central targets for MEF2A-mediated transcription, with NFIC driving divergent transcriptional programming responses to EE and RE–with pronounced changes of mitochondria NFIC target genes following EE and translation genes with RE (Fig. 7g).
